# Inhibition of lagging strand replication by G-rich telomeric DNA and the shelterin subunit POT1

**DOI:** 10.1101/2025.11.13.688213

**Authors:** Ciara Leonard-Booker, Giulia Mazzucco, Charlotte E.L. Fisher, Karim Hussain, Tom D. Deegan, Ylli Doksani, Max E. Douglas

**Affiliations:** Telomere Biology Laboratory, The Institute of Cancer Research, 237 Fulham Road, London SW3 6JB, UK; IFOM ETS-The AIRC Institute of Molecular Oncology, Milan, Italy; MRC Human Genetics Unit, University of Edinburgh, EH4 2XU, UK

## Abstract

Telomeres preserve stable eukaryotic chromosomes by protecting the natural chromosome ends from DNA repair but pose a persistent challenge to the replication machinery and an endogenous source of replication stress. Different features have been implicated in causing this effect but how the canonical replication process is altered at telomeres in mechanistic terms remains poorly understood. To address this question, we have reconstituted telomere replication with purified human proteins. This system reveals that while G-rich telomeric DNA directly and specifically blocks lagging strand replication in a manner counteracted by BLM helicase, shelterin unexpectedly acts as an additional lagging strand barrier. Biochemical experiments and electron microscopy imaging show that POT1-containing shelterin complexes induce Okazaki fragment skipping by binding the lagging strand template, generating large single-stranded gaps that remain unreplicated. Our study defines how the core components of telomeres interfere with the canonical replication process, identifying several potential sources of replication stress.

## INTRODUCTION

In human cells, stable linear chromosomes are maintained by telomeres composed of 10-15 kilobases of the repetitive sequence TTAGGG and shelterin, a six-subunit protein complex comprising TRF1, TRF2, RAP1, TIN2, TPP1 and POT1^1^. Shelterin binds tightly to double-stranded telomeric DNA through TRF1 and TRF2 and single-stranded G-rich repeats via POT1, forming a nucleoprotein assembly that blocks inappropriate DNA repair processes including homologous recombination (HR), non-homologous end joining (NHEJ) and checkpoint activation^2^. Loss of these functions at even a single chromosome end can disrupt chromosomal structure in human cells, driving cell cycle arrest or cell death^3^.

In proliferating cells, telomeres are maintained through semiconservative DNA replication. Recent advances have established a canonical view of the replication process in which the replicative <u>C</u>dc45, <u>M</u>cm2-7 and <u>G</u>INS (CMG) helicase and its associated proteins Claspin, Tipin-timeless and AND-1 unwind parental DNA by translocating 3’-5’ along the leading strand template^4^. Continuous leading strand synthesis is mediated by polymerase ε (Pol ε)^5^, while discontinuous lagging strand synthesis by Polymerase α/primase (Pol α/prim)^6^ and Polymerase δ (Pol δ)^4^ is followed by maturation of the lagging strand into a continuous product through a set of factors including Flap Endonuclease 1 (FEN1) and DNA Ligase 1 (LIG1)^4^.

In order for complete duplication of the genome, these processes must traverse complex templates bound to proteins, assembled into R-loops or containing secondary structures that can induce copying defects termed replication stress^7^. In mammalian cells, stress-prone regions have been identified as ‘fragile sites’ characterised by sensitivity to low concentrations of replication inhibitors^8^. Telomeres are recognised as fragile sites that are prone to fork stalling and replication stress in mouse and human cells^9,10^. As fork arrest is also observed at telomeres in budding and fission yeasts^11,12^, telomere replication may pose conserved challenges across many eukaryotic species, with the potential for loss of telomeric DNA unless these challenges are handled effectively.

An extensive number of studies have implicated a range of telomeric properties in this effect (reviewed in^13,14^), including the components that endow telomeres with their protective functions: TTAGGG repeats can fold into G-quadruplex (G4) structures that hinder replicative polymerases and the CMG helicase in vitro^15–18^, and mutations in G4 unwinding helicases or treatment with G4-stabilising ligands cause fragile telomeres in mouse and human cells consistent with sequence playing an inhibitory role^19–24^. However, whether telomeric DNA can directly influence the replication process and which stages are affected remains to be determined. The shelterin complex, which is involved in essentially all protective functions of telomeres, binds directly to telomeric repeats and TRF1 and 2 can block an SV40 replisome in cell extracts^25,26^, suggesting that in addition to recruiting factors that promote telomere replication^9,27^, shelterin may also prove inhibitory. Yet, how the human replisome is affected by the tight binding of shelterin to single- and double-stranded telomeric DNA remains unclear.

Addressing these questions requires a system in which the impact of different telomeric components on specific aspects of DNA replication can be studied in isolation and in molecular detail. Here, we achieve this goal by reconstituting telomere replication with purified human proteins. Our results reveal distinct inhibitory effects of the core components of telomeres on the canonical replication process, uncovering a potent inhibitory effect of POT1 on copying of the lagging strand.

## RESULTS

### Telomeric DNA does not appreciably slow or stall leading strand synthesis

To determine how the canonical replication process is altered at telomeres, we set out to combine a previously reconstituted assay for human DNA replication with templates containing different telomeric properties (Fig. S1a). In this system^28^, a set of eight purified replication fork components (Fig. 1a) is used to build a human replisome on a forked DNA template that supports CMG- and Pol ε-dependent leading strand synthesis visualised through the incorporation of α-^32^P-labelled dCTP (Fig. 1c). Consistent with a previous study^28^, pulse chase assays containing a full complement of leading strand replication factors revealed that leading strand synthesis proceeds at a rate of 3.8 kb/min in vitro (Fig. S1b), and non-pulse chase assays showed that replication forks slowed significantly without either AND-1, Claspin or Tipin-Timeless (Fig. S1c).

**Figure 1.**
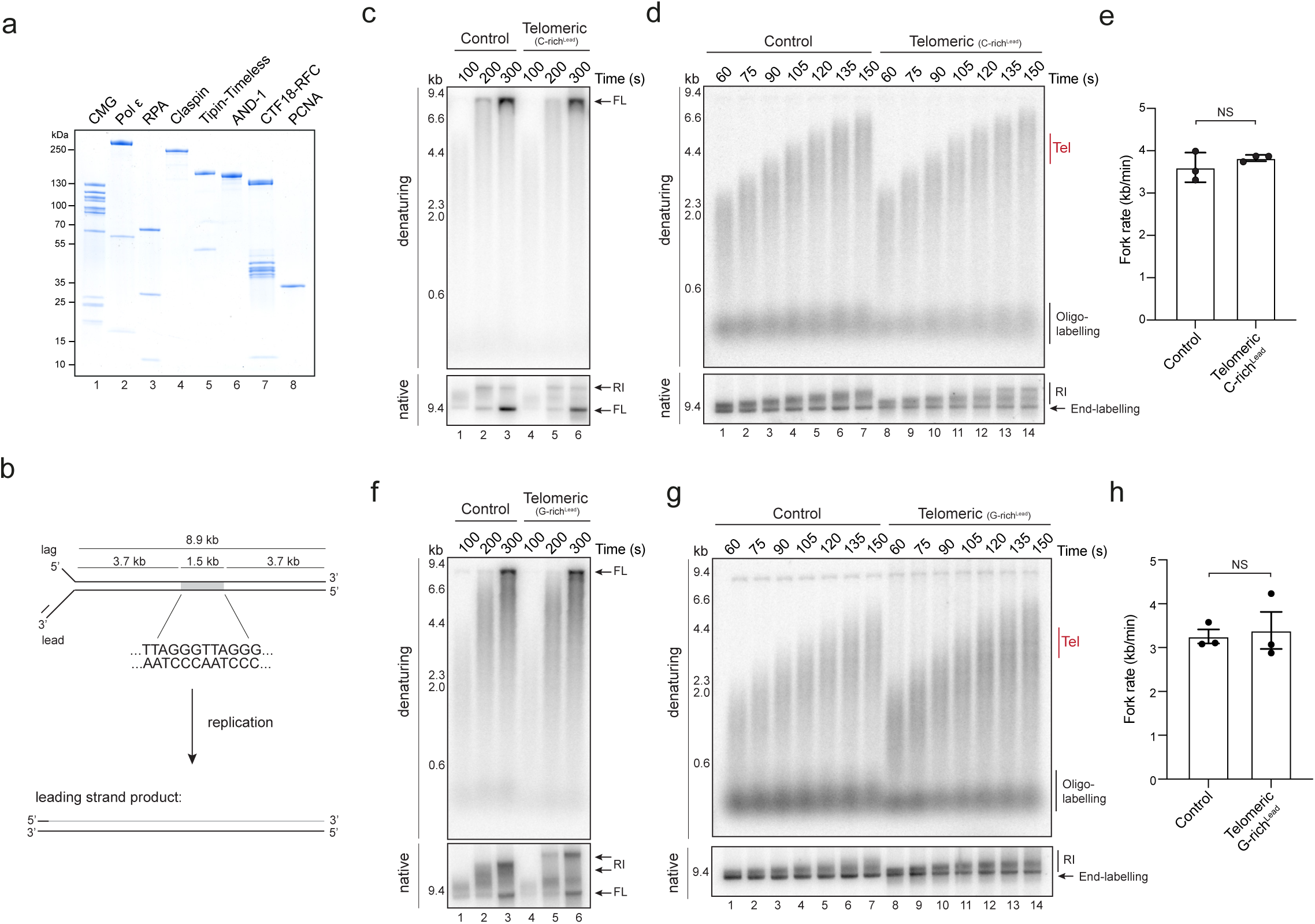
Human telomeric DNA does not inhibit leading strand synthesis or replisome progression. **(a)** SDS-PAGE analysis of purified human replisome factors. **(b)** Schematic showing the telomeric template used in replication reactions and expected leading and lagging strand products. The position and orientation of the telomeric DNA is highlighted. **(c)** Replication reactions performed with control and telomeric templates for the indicated times analysed by native and denaturing alkaline agarose electrophoresis as indicated. Full-length (FL) and replication intermediates (RI). **(d)** Pulse chase analysis with control and telomeric templates. Chase added after 50 s and samples taken at the indicated times after initiation and analysed by native and denaturing alkaline agarose electrophoresis as indicated. Site of telomeric insert indicated (Tel). **(e)** Quantification showing maximal fork rates calculated from three independent pulse chase experiments with control or telomeric templates as indicated. Error bars show the S.E.M. Unpaired t-test shows P > 0.05 **(f-h)** As for c-e, except comparing control and flipped telomeric templates, where G-rich telomeric DNA is the sequence for leading strand replication.

To adapt this system to study telomeres, the first feature we chose to incorporate was telomeric DNA. The repetitive, GC-rich nature of telomeres has been proposed to inhibit DNA replication, but whether this is the case and how different replisome functions are affected by telomeric sequence remains unclear. A 1.5 kb tract of TTAGGG repeats was inserted into the replication substrate oriented with C-rich telomeric DNA as the leading strand template, reflecting the orientation of most telomere replication events in vivo (Fig. 1b)^9,29^. Figure 1c shows that relative to a non-telomeric control, this template was replicated efficiently, with no apparent stalling or pausing of leading strand synthesis at the position of telomeric repeats (Fig. 1c, compare lanes 1-3 and 4-6). Pulse chase assays in which a subset of replication forks is followed along the template confirmed that bulk replication rate was unaffected by TTAGGG repeats with no appreciable pausing or stalling of synthesis (Figs. 1d and e, Fig. S1d). Thus, up to 1.5 kb of C-rich telomeric DNA can be copied efficiently as the leading strand without apparently stalling the replication fork.

DNA replication occasionally initiates within telomeric repeats, leading to instances in which CMG translocates along G-rich telomeric DNA^19,29^. Without the BLM helicase, this scenario is reported to inhibit fork progression due to G-quadruplexes forming on the leading strand template^19^. Reversing the orientation of telomeric repeats did not reveal a strong stalling effect of G-rich DNA in our system, with no significant change in bulk replication rate compared with the control template (Figs. 1f-h and Fig. S1e). G-rich sequences on the leading strand template have previously been shown to block a reconstituted budding yeast replisome in the presence of G4 stabilising compounds^30^. Consistently, the hydroquinone compound phenDC3 induced a mild accumulation of leading strands on telomeric DNA (Fig. S1f, compare lanes 5 and 6). However, most replication events nonetheless proceeded to the end of the template unless high phenDC3 concentrations that also inhibit C-strand replication were used (Figs. S1f and g), suggesting G-rich telomeric DNA alone is unlikely to be a strong leading strand barrier.

### G-rich telomeric DNA prevents complete replication of the lagging strand

In most instances, G-rich telomeric DNA is the template for the lagging strand. This orientation may be particularly problematic because unlike the leading strand, which is synthesised directly behind CMG^5^, Okazaki fragment (OF) synthesis by Pol δ is uncoupled from DNA unwinding, potentially providing more opportunities for DNA structures to form.

To test this idea, we included the machinery for lagging strand synthesis into our reactions (Fig. 2a). As reported previously^28^, addition of RFC1-5 and Pol α/prim led to the synthesis of OFs 200-500 nt long that were extended in the presence of Pol δ (Fig. S2a). Purified FEN1 and LIG1 were then added to mature OFs into a continuous lagging strand product. Figure S2b shows that together but not individually, FEN1 and LIG1 caused an increase in unit length products and the disappearance of OFs, suggesting they had been processed and ligated together successfully (Fig. 2b and S2b and c). To confirm that this was the case, replication products were treated with a site specific nickase primarily targeting the leading strand, revealing a full length lagging strand product only in the presence of maturation factors (Fig. S2d, see Fig. 2c for nicking positions). This data demonstrates that as for the budding yeast replisome^31^, FEN1 and LIG1 represent a minimal set of components for lagging strand maturation at a human replication fork.

**Figure 2.**
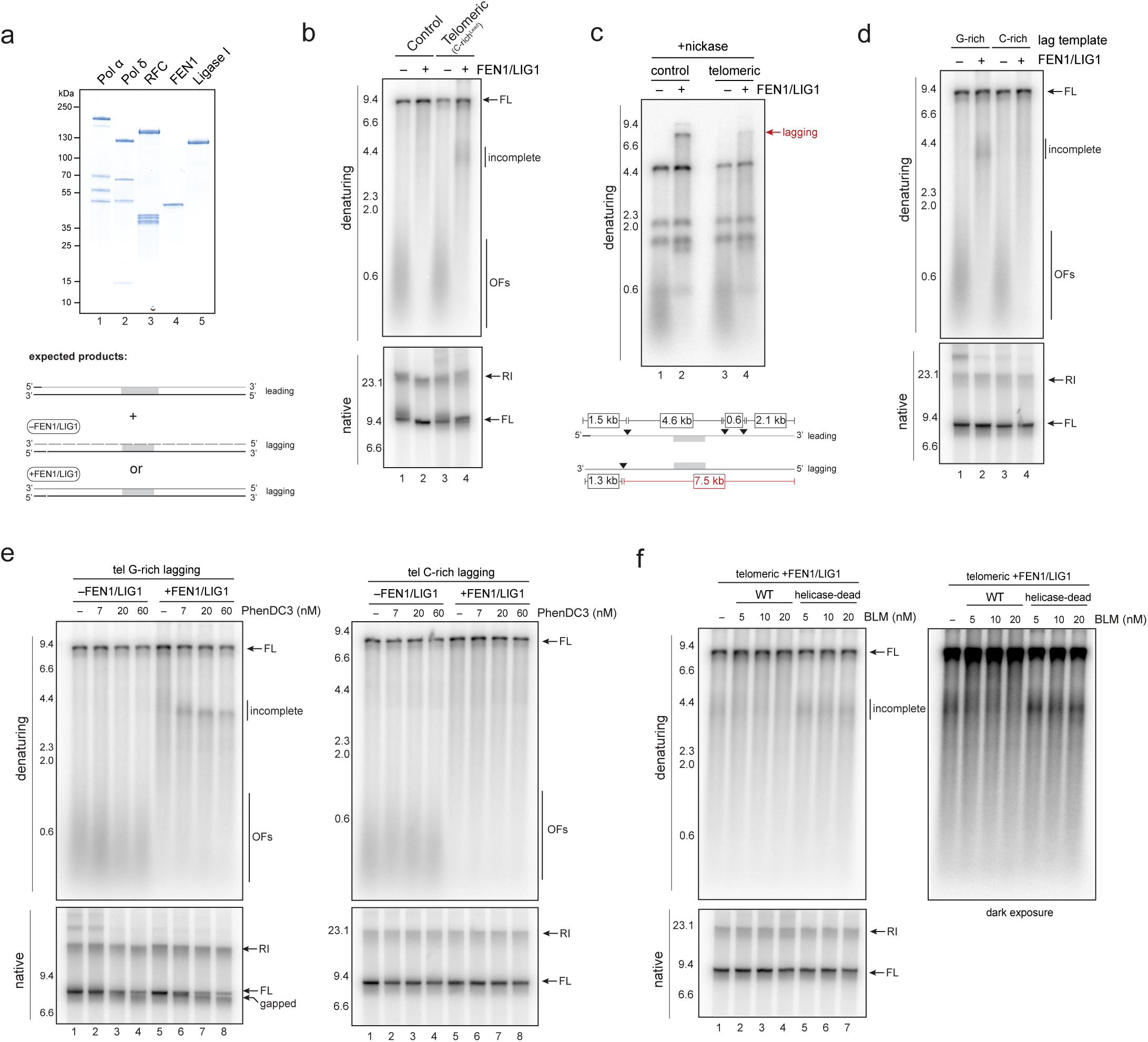
G-rich telomeric DNA directly inhibits lagging strand replication. **(a)** SDS-PAGE analysis of purified human lagging strand replication factors and schematic showing leading and lagging strand products expected with and without FEN1/LIG1. **(b)** Replication reactions performed for 30 min in the presence or absence of FEN1/LIG1 with the control and telomeric template as indicated were analysed by native or denaturing alkaline agarose electrophoresis as indicated. Okazaki fragments (OFs). **(c)** Products from replication products performed for 30 min on the templates indicated in the presence or absence of FEN1/LIG1 were treated with nicking enzyme Nb.BsrDI for 2 h were analysed by denaturing alkaline agarose electrophoresis. Schematic shows position of Nb.BsrDI sites and products expected from the leading and lagging strand. **(d)** Products from replication reactions performed for 30 min in the presence or absence of FEN1/LIG1 with G- or C-rich telomeric DNA as the lagging strand template as indicated were analysed as in b. **(e)** Products from replication reactions performed for 30 min in the presence or absence of FEN1/LIG1 and indicated concentrations of phenDC3 with G- or C-rich telomeric DNA as the lagging strand template were analysed as in b. **(f)** Products from replication reactions performed for 30 min with telomeric template in the presence of FEN1/LIG1 and indicated concentrations of BLM wt or helicase dead were analysed as in b.

As full replication of the lagging strand requires OF synthesis across the entire template, we used this system to determine whether replication can be completed on G-rich telomeric DNA. Figure 2b shows that with the telomeric template, FEN1/LIG1 addition resulted in a population of products around 4 kb in length, consistent with incomplete lagging strand replication across the telomere (‘incomplete’ products, compare lanes 2 and 4, see figure 1 for template organisation). In line with this idea, nicking assays revealed a reduction in the level of fully replicated lagging strands on the telomeric-compared with the control template (Fig. 2c) and flipping the orientation of the telomere revealed this defect was caused by G-rich telomeric DNA rather than the repetitive sequence per se (Fig. 2d). To determine whether this defect might reflect the ability of telomeres to form G-quadruplexes, phenDC3 was added. Even low concentrations of phenDC3 led to the appearance of a strong band of defective replication products when FEN1/LIG1 were added and G- but not C-rich telomeric DNA was the lagging strand template (Figs. 2e). The presence of putative ‘gapped’ replication products that run slightly smaller than full length after native agarose electrophoresis suggests a strong defect in lagging strand synthesis over the telomere under these conditions (Fig. 2e bottom panel ‘gapped’ product). Thus, G-rich telomeric DNA directly inhibits replication of the lagging strand.

The BLM helicase can unwind G4s in vitro^20,21,32^ and has been suggested to remove DNA structures on the lagging strand during telomere replication^23^. Yet whether BLM can directly prevent sequence-based replication defects remains to be determined. To test this idea, wild type BLM and a helicase inactive K695A mutant were purified (Fig. S2e) and titrated them into our reactions. Figure 2f shows that helicase active BLM, but not the inactive mutant, reduced the level of lagging strand defects over the telomere, demonstrating that BLM can directly overcome the inhibitory effect of G-rich telomeric DNA.

In summary, up to 1.5 kb of neither C- nor G-rich telomeric DNA has a strong impact on leading strand synthesis. In contrast, the G-rich strand can block lagging strand replication in a manner that is exacerbated by G-quadruplex stabilising agents and counteracted by BLM helicase activity, consistent with previous genetic work^23^.

### Shelterin can act as a replication roadblock

We next turned our attention to the main factor bound to telomeric DNA: the shelterin complex. There are estimated to be tens to hundreds of shelterin molecules at each human telomere^33,34^. A previous study with SV40 large T-antigen suggests that these complexes can block or slow a viral replisome^25^, but deletion of shelterin subunits such as TRF1 in mouse and human cells impedes rather than promotes the replication process ^9,35^. Whether this is also case for a human replication fork is therefore unclear.

To determine the impact of shelterin on the human replisome, we purified a six-subunit complex of TRF1, TRF2, TIN2, RAP1, TPP1 and POT1 that bound tightly to telomeric DNA in electrophoretic mobility shift assays (Fig. 3a and Figs. S3a and S3b). Titrating this complex into leading strand only replication reactions containing a template with a telomeric tract of only sixty TTAGGG repeats caused up to 60% of leading strands to accumulate as a tight band of telomere-arrested products (Fig. 3a). A similar effect was observed in reactions competent for lagging strand synthesis (Fig. S3b) and a distinct band of replication intermediates following native agarose electrophoresis suggested that progression of the replication fork along the template rather than just leading strand synthesis had failed (Fig. 3a, ‘stall’ product. Also discussed below). Thus, even at relatively modest concentrations, shelterin has the potential to act as a replication roadblock. To determine whether shelterin arrested forks are stalled or simply slowed, we performed a pulse chase analysis in which the chase was added 50 seconds after initiation factors. Figure 3b shows that a persistent band of telomere-arrested products was present across the full 60 minutes, suggesting the replisome alone is ineffective at bypassing shelterin once a stall has taken place (Fig. 3b).

**Figure 3.**
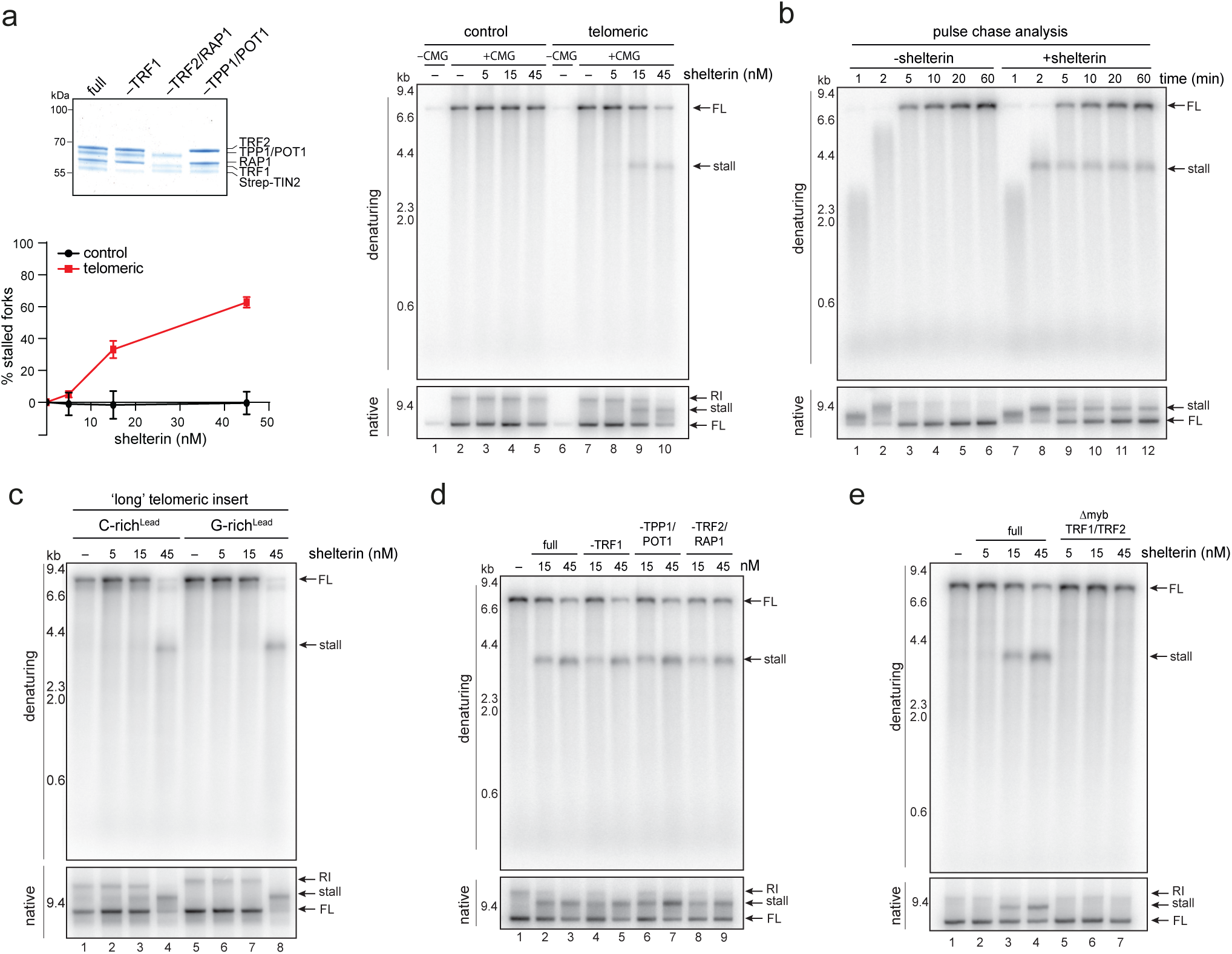
Shelterin can stall a human replisome on telomeric DNA. **(a)** SDS-PAGE analysis of purified shelterin variants. Replication reactions performed for 20 min on ‘short’ control or telomeric templates in the presence or absence of shelterin at the indicated concentrations were analysed by native and denaturing alkaline agarose electrophoresis as indicated. Position of the stall is indicated. Note the shorter telomeric length compared with figures 1 and 2. The percentage stalled forks after denaturing alkaline electrophoresis compared with reactions lacking shelterin is shown. Average of three independent experiments. Error bars show S.E.M. **(b)** Pulse chase analysis with ‘short’ control and telomeric templates in the presence or absence of shelterin. Chase added after 50 s and samples taken at the times indicated after initiation and analysed by native and denaturing alkaline agarose electrophoresis. **(c)** Products from replication reactions performed for 20 min with ‘long’ templates containing C- or G-rich telomeric DNA as the leading strand template with the shelterin concentrations indicated were analysed as in a. **(d)** Products from replication reactions performed for 20 min on ‘short’ telomeric templates in the presence or absence of shelterin or shelterin variants as indicated were analysed as in a. **(e)** Products from replication reactions performed for 20 min on ‘short’ telomeric template with shelterin variants indicated were analysed as in a. Δmyb TRF1/TRF2 - shelterin in which both TRF1 and TRF2 are lacking myb domains.

We probed the requirements for this effect. Shelterin was equally effective at stalling the replisome when either C- or G-rich telomeric DNA was the leading strand template (Fig. 3c). However, at the same shelterin concentration, a moderately weaker inhibitory effect was observed when more telomeric repeats were present (Fig. S3c, 15 nM shelterin), suggesting stalling may be sensitive to shelterin density on the template. To determine whether a particular subunit was responsible, we tested shelterin variants lacking TRF1, TRF2/RAP1 or TPP1/POT1. Figures 3d and e show that while no single shelterin subunit was essential, the myb domains of TRF1 and TRF2 were necessary for stalling to occur. Thus, binding of shelterin to telomeric DNA via either TRF1 or TRF2 can block progression of the replisome, and the density of shelterin on the template may influence the magnitude of this effect.

### Shelterin induces large lagging strand gaps during DNA replication

To probe the organisation of shelterin-stalled forks, replication reactions competent for leading and lagging strand synthesis but without maturation factors were performed in the presence or absence of shelterin, subjected to psoralen-UV crosslinking and imaged by electron microscopy (EM). To control for molecules on which CMG-independent strand displacement synthesis had taken place, we performed this analysis with and without Pol δ. Most visualised molecules were linear, likely because only a fraction was replicated. However, we observed an enrichment of Y-shaped replication forks when shelterin was added, the dimensions of which were consistent with most having stalled at the telomere (Fig. 4a. –Pol δ - 7 % (61 out of 826) Y-shaped with 15 nM shelterin compared with 2 % (11 out of 657) with no shelterin; + Pol δ - 7 % (65 out of 980) Y-shaped with 15 nM shelterin compared with 3 % (28 out of 981) with no shelterin).

**Figure 4.**
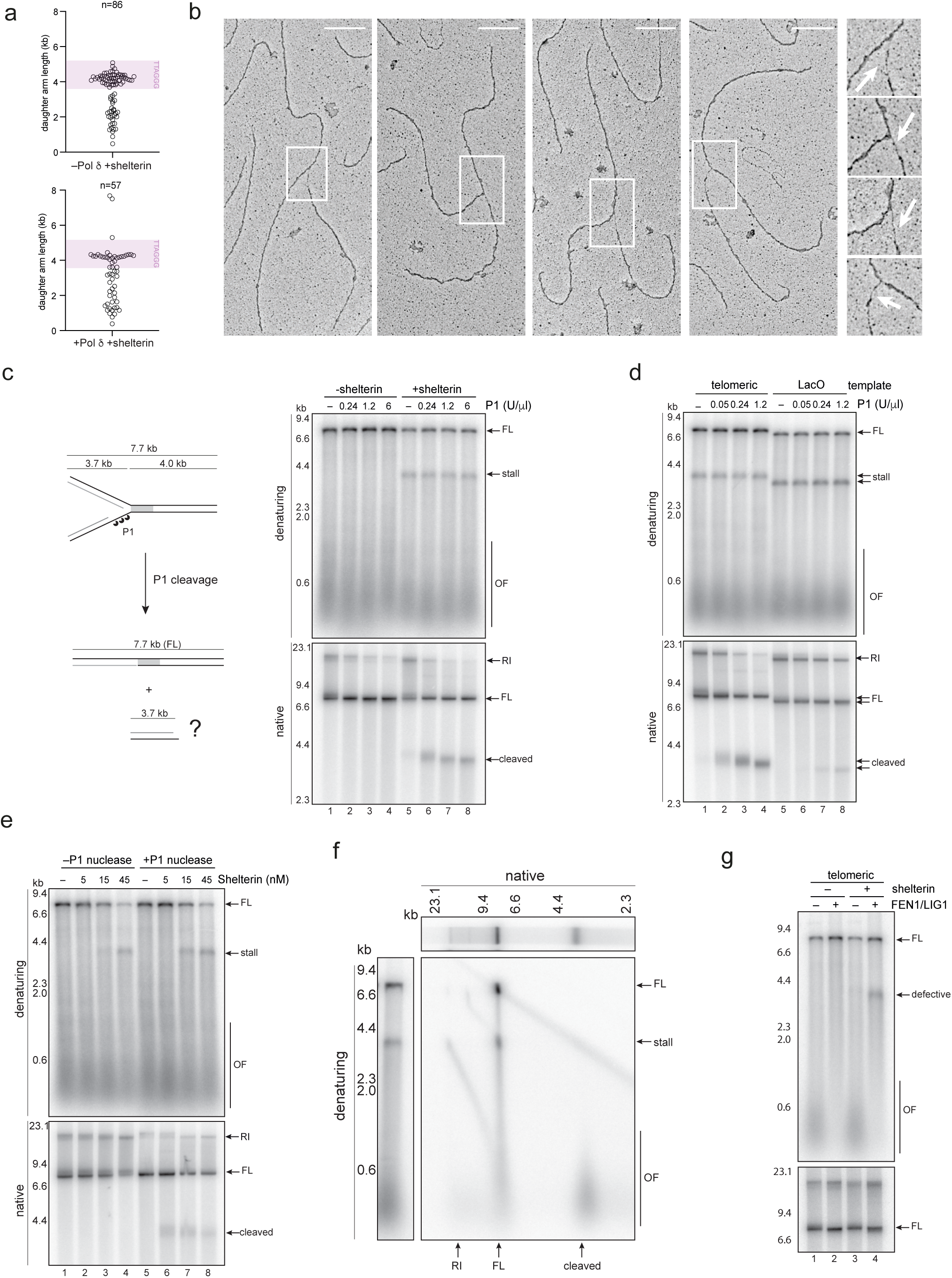
Shelterin induces large single-stranded gaps on the telomeric lagging strand. **(a)** Quantification of daughter arm lengths in the presence of shelterin and presence or absence of Pol δ. Replication reactions were performed for 15 min in the presence of 15 nM shelterin before crosslinking and EM analysis. **(b)** Examples of Y-shaped molecules identified in the presence of shelterin. Regions of single-stranded DNA indicated by the white rectangles, which are also enlarged. **(c)** Replication reactions performed for 15 min on ‘short’ telomeric template in the presence or absence of shelterin and the indicated amounts of P1 nuclease added 5 min before quenching. Products analysed by native and denaturing alkaline agarose electrophoresis as indicated. Schematic shows products expected after cleavage of gapped replication forks. **(d)** Products from replication reactions performed for 15 min on ‘short’ telomeric or LacO template with shelterin, LacI and the indicated amount of P1 nuclease were analysed as in c. P1 nuclease was added 5 min before quenching reactions. **(e)** Products from replication reactions performed for 15 min on ‘short’ telomeric template in the presence or absence of indicated shelterin concentrations were analysed as in c. 0.24 U/ml P1 nuclease was added 5 min before quenching reactions. **(f)** Replication products from a 15 min reaction containing ‘short’ telomeric template and 15 nM shelterin were analysed by native agarose electrophoresis (top panel). 1 U/ml P1 nuclease was added 5 min before quenching reactions. The lane was excised and analysed by denaturing alkaline agarose electrophoresis (left hand panel). **(g)** Replication reactions performed on the ‘short’ telomeric template with shelterin and lagging strand maturation factors as indicated were analysed by native and denaturing agarose electrophoresis as indicated.

Examining these structures revealed that ∼70 % (46 out of 70 –Pol δ, 27 out of 37 +Pol δ) contained hundreds of nucleotides of single stranded DNA (ssDNA) exposed on one of the daughter strands at the fork junction (Fig. 4b and Fig. S3d). This was unexpected given that a protein-blocked budding yeast replisome contained single-stranded gaps less than 30 nt in length^36^. To confirm this result, we treated our reactions with the single strand-specific P1 nuclease. Figure 4c shows that no specific P1 cleavage product was observed in the absence of shelterin. However, in its presence, a distinct 3.7 kb product was visible after native agarose electrophoresis that matches the size expected if one arm had been cleaved off the stalled replication fork (Fig. 4c, ‘cleaved’ product). Notably, at least 5-10 times more P1 was required to induce a similar level of cleavage with replication forks stalled by the Lac repressor, suggesting exposure of such large single-stranded tracts is specific to shelterin and telomeric DNA (Fig. 4d, compare lanes 1-4 and 5-8).

We examined whether formation of these single-stranded gaps required stalling of the replisome or was a general outcome of copying through shelterin-bound TTAGGG repeats. No persistent fork stalling was observed when the concentration of shelterin was decreased from 15 nM to 5 nM (Fig. 4e). However, replication products were nonetheless susceptible to P1 cleavage (Fig. 4e, compare lanes 6 and 7), indicating that lagging strand gaps can be induced without a persistent stalling event. Consistently, the number of full-length molecules on electron micrographs that were linear but contained ssDNA over the telomere increased in the presence of shelterin (Fig. S3e and f. –Pol δ: 4.6 % +shelterin (35 out of 765), 0.8 % –shelterin (5 out of 646); +Pol δ: 3.5 % +shelterin (32 out of 915), 0.5 % –shelterin (5 out of 953)).

To determine whether these gaps were specific to one or other replicated strand, P1-cleaved products from replication reactions containing shelterin were separated by native agarose electrophoresis and run in a second, denaturing dimension. Figure 4f shows that the 3.7 kb P1-cleaved product contained OFs but no corresponding leading strand and therefore derives from the lagging strand (Fig. 4f). Consistently, 360 bp of telomeric DNA was insufficient to induce a strong lagging strand defect in the presence of FEN1 and LIG1, but a prominent band of defective lagging strand products was observed when shelterin was also added (Fig. 4g).

### Lagging strand gaps are induced by POT1 and can be suppressed by CST

The data above reveals that in parallel to preventing the replisome from progressing along the template, shelterin can inhibit replication of the telomeric lagging strand. To determine the cause of this effect, we tested different shelterin subcomplexes in replication reactions treated with P1. Figure 5a shows that while P1 cleavage could occur with shelterin lacking TRF1 or TRF2/RAP1, this was not the case when TPP1/POT1 was omitted (compare lanes 4 and 8). Figure 5b shows that TPP1/POT1 was necessary for cleavage but not sufficient as no P1 sensitivity was observed without other shelterin subunits (Fig. 5b). Given that TPP1/POT1 is recruited via TRF1 or TRF2, this data suggests that TPP1/POT1 localised to telomeric DNA as part of shelterin caused the inhibitory effect on the lagging strand.

**Figure 5.**
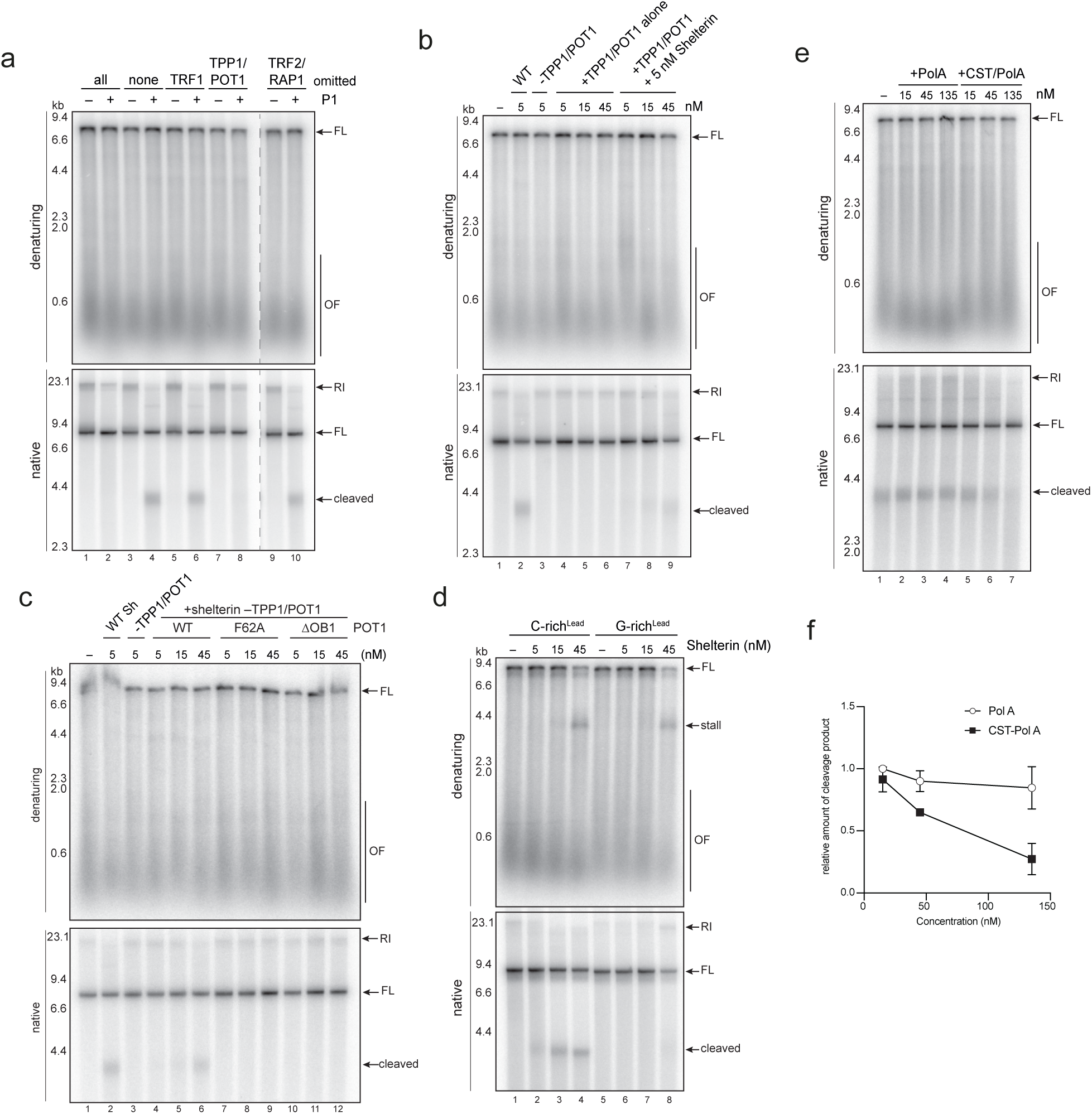
Lagging strand gaps are induced by POT1 binding to single-stranded G-rich DNA. **(a)** Replication reactions performed for 15 min on ‘short’ telomeric template with shelterin variants and 0.24 U/ml of P1 nuclease as indicated. Products were analysed by native and denaturing alkaline agarose electrophoresis as indicated. **(b)** Products from replication reactions performed for 15 min on ‘short’ telomeric template with shelterin, shelterin -TPP1/POT1, TPP1/POT1 alone or TPP1/POT1 with 5 nM shelterin -TPP1/POT1 as indicated were analysed as in a. 0.24 U/ml P1 nuclease was added 5 min before quenching reactions. **(c)** Products from replication reactions performed for 15 min on ‘short’ telomeric template with wt shelterin, or shelterin -TPP1/POT1 preincubated with the TPP1/POT1 variants indicated were analysed as in a. 0.24 U/ml P1 nuclease was added 5 min before quenching reactions. **(d)** Products from replication reactions performed for 15 min in the presence of the indicated shelterin concentrations with ‘long’ templates in which C- or G-rich telomeric DNA is the leading strand template were analysed as in a. 0.24 U/ml P1 nuclease was added 5 min before quenching reactions. **(e)** Products from replication reactions performed for 30 min on ‘short’ telomeric template in the presence of 5 nM shelterin and indicated concentrations of CST-pol a/prim or pol a/prim were analysed as in a. 0.035 U/ml P1 nuclease was added 5 min before quenching reactions. The concentration of RPA was reduced to 30 nM. **(f)** Quantification of the level of P1 cleavage product after native agarose electrophoresis from three independent repeats of the experiment in e relative to reactions containing 15 nM Pol a/prim. Error bars shows S.E.M.

POT1 prevents ATR activation at telomeres by binding tightly to G-rich single-stranded DNA through the first OB fold, which is frequently mutated in cancer^37^. As G-rich DNA was the template for lagging strand synthesis in these experiments, we hypothesised that POT1 may block lagging strand synthesis by simply binding to the unwound G-rich template. P1 cleavage was not observed with shelterin complexes containing POT1 mutants defective in ssDNA binding^38^ (Fig. 5c), or when the orientation of telomeric DNA was reversed so that the C-rich strand was the template for lagging strand synthesis (Fig. 5d), indicating that this was indeed the case.

Internal single stranded gaps on the telomeric lagging strand are not broadly observed in wild type cells. However, they are reported to occur in the absence of the Pol A/primase co-factor CST^39–41^. As CST is also known to bind TPP1/POT1^42–45^, we considered whether it might prevent the inhibitory effect of POT1. Figure 5e and f show that with a reduced concentration of RPA, addition of CST and Pol α/prim but not Pol α/prim alone reduced the sensitivity to P1. The relatively high CST concentrations required for this effect may indicate that P1 cleavage is not linear with respect to single-stranded DNA length, but may also be consistent with previous work showing phosphoregulation is required for POT1 to recruit CST to telomeres^44^.

## DISCUSSION

The cause of telomeric replication stress has been the focus of many studies that have revealed a range of properties that can perturb the replication process^13,14^. Yet, the pleiotropic effect of mutating many telomeric components, as well as the large number of factors bound to telomeric DNA has meant that the direct impact of different components on the replisome has been challenging to examine using genetic approaches alone. Here, we have overcome this hurdle by using a reconstituted system for human DNA replication to reveal how telomeric DNA and the shelterin complex directly impact on different aspects of the replication process. Our study provides a molecular description of the challenges encountered by the replication fork as it navigates a core telomeric template (Fig. 6).

**Figure 6.**
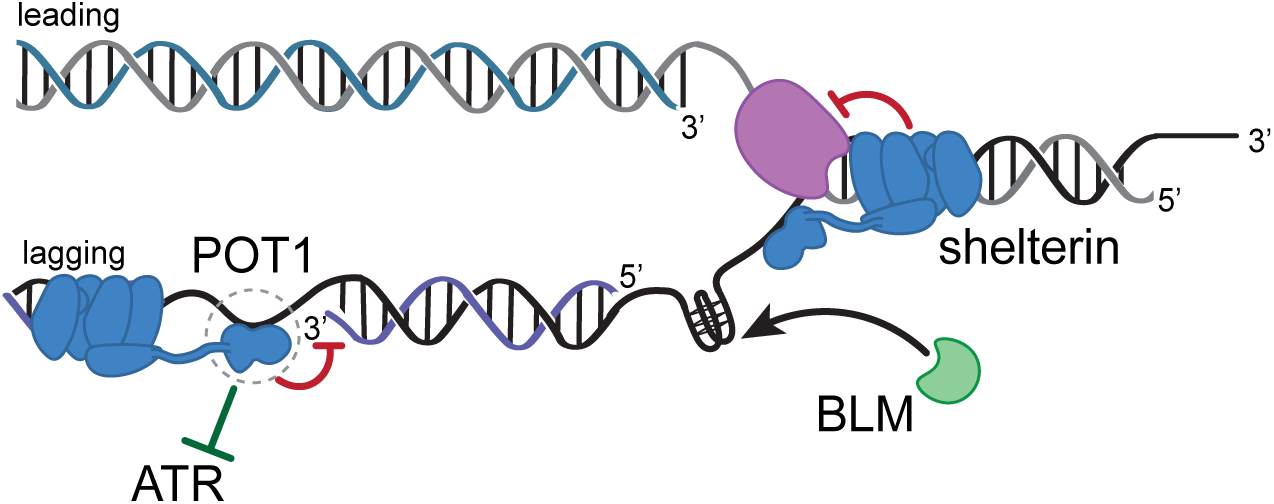
A model for the impact of shelterin and telomeric DNA on the telomere replication process. See main text for details.

We show that G-rich telomeric DNA directly inhibits lagging strand replication by the human replisome. As lagging strand telomere replication is exquisitely sensitive to G4 stabilising agents and promoted by the G4-unwinding helicase BLM in vitro, we favour that this effect is primarily caused by G-quadruplexes assembled on the unwound lagging strand template, in line with cell-based studies implicating G4s in telomeric replication stress^46,47^. However, as G-rich telomeric DNA is inhibitory for Pol δ independently of G4s in primer extension assays^48^, other properties may feed into this effect. Leading strand replication is apparently much less sensitive to the same G-rich sequence even in the presence of G4 stabilising agents, consistent with coupled DNA unwinding and DNA synthesis on the leading strand restricting interference from DNA secondary structures. Our data is consistent with a primary role for BLM at telomeres being to remove sequence-based structures on the lagging strand template, as has been previously suggested by cellular studies^9,23^. Notably, binding of BLM to TRF1 is required for this function in cells but is apparently dispensable in vitro, perhaps indicating that shelterin recruits BLM to telomeres and away from non-telomeric sites within the nucleus.

While telomeric sequence primarily inhibits the lagging strand, shelterin has the potential to stall synthesis entirely by acting as a replication roadblock. In a similar manner, TRF1 and TRF2 can stall the SV40 replicative helicase large T-antigen in cell extracts^25,26^ and may also do so in cells^10^ and Rap1, the primary telomere binding protein in budding yeast, is a potent replication barrier in vitro and in vivo^17,49,50^. EM analysis indicates that most shelterin-stalled forks exhibit a classic Y-shape structure in vitro. However, as protein-stalled forks can act as a substrate for fork reversal^51^, this may not be the case in vivo. Indeed, reversed forks are enriched at telomeres in mammalian cells^10^, and a recent study suggests that slowing or stalling of the replisome by TRF1 leads to reversed forks that prevent telomere fragility in human cells^52^. Whether shelterin-stalled forks are a direct substrate for reversal enzymes is an important question for the future. We consider it likely that mechanisms are in place to prevent stalling at shelterin: in eukaryotic cells, protein-based replication blocks are overcome by helicase proteins working alongside the replisome, including budding yeast Rrm3^53^ and RTEL1 in mammals^54^. Rrm3 and RTEL1 are required for telomere replication in yeasts and humans respectively^12,22^, and it is interesting to consider that in addition to its role unwinding G4s and t-loops^22,46^, RTEL1 may restrict the extent to which replication forks stall at DNA-bound shelterin molecules. Challenges purifying active RTEL1 have so far prevented us from directly testing this idea. The second feature that may attenuate the stalling effect of shelterin is chromatin. TRF1 makes direct contact with histones and can remodel nucleosomes^55–57^, while binding of TRF2 to telomeric DNA is inhibited by nucleosomes^58^. Whether these distinct effects influence the impact of shelterin by modulating its density along the template remains to be determined.

Crucially, shelterin is also a potent and specific inhibitor on the lagging strand, where binding of POT1 to single stranded G-rich DNA induces large unreplicated gaps. We consider this a secondary but penetrant consequence of the essential role of POT1 in binding to ssDNA to prevent ATR activation at telomeres^59–61^. Based on the concentration of components in our assays, only a few POT1 molecules are required on each template for this inhibition to take place. Which stage of lagging strand synthesis is blocked? With low shelterin concentrations, internal lagging strand gaps are left behind as the fork continues along the template, indicating that POT1 or a structure induced by POT1 must prevent Pol δ from extending the next Okazaki fragment into the gap. We favour that POT1-containing shelterin complexes are left behind on the defective lagging strand as a persistent barrier, perhaps explaining why POT1 is depleted from replication forks structures at telomeres^62^. We do not rule out that POT1 also interferes with the priming process: OF priming is undertaken by Pol α/prim, which accesses the newly unwound template via an interaction with CMG^6^. It is interesting to consider that POT1, and perhaps other sequence-specific ssDNA binding proteins at different regions of the genome, could compete with Pol α/prim to interfere with this process. How are such large, POT1-induced gaps prevented? The Pol α/prim cofactor and TPP1/POT1 binding partner CST is known to prevent internal telomeric gaps on the G-rich strand in human cells^39–41^ and reduces the sensitivity to P1 nuclease in vitro, consistent with it playing a role. Binding to CST could allow POT1 to fulfil its function preventing ATR activation while avoiding the inhibitory effect on the lagging strand; how regulation of this interaction by phosphorylation^44^ feeds into this effect is an important question for the future. It is also possible that POT1 is locally displaced by a helicase working in conjunction with pol δ on the lagging strand.

Beyond the findings above, the reconstituted system reported here provides a new tool with which to understand how further telomeric features including the non-coding RNA TERRA and the unique chromatin environment at telomeres influence and perturb the replication process.

## METHODS

### DNA construct assembly

For *in vitro* replication reactions, an 8.8 or 7.8 kb plasmid containing an insert of 1440 or 360 bp of telomeric DNA, respectively was constructed from a pUC19 vector backbone. A corresponding control construct was also generated containing a fragment of non-telomeric control sequence. To generate the LacO template, a fragment containing four lac operator sequences interspersed with 11 bp of random sequence was excised from a plasmid (a kind gift from John Diffley) by restriction endonuclease digestion with EcoRI-HF (New England Biolabs, NEB) and BamHI-HF (NEB). This LacO fragment was inserted into the 7.8 kb control template digested with EcoRI-HF and BamHI-HF. The sequences of telomeric and non-telomeric replication templates used in the study are provided in the ‘Supplementary Information’ file.

For expression plasmid constructs, cDNAs encoding all subunits of CMG (MCM2, MCM3, MCM4, MCM5, MCM6, MCM7, CDC45, PSF2, PSF3, PSF5, SLD5), Polymerase ε (POLE1, POLE2, POLE3, POLE4), Tipin-Timeless, Claspin, AND-1, RFC1, RFC2-3, RFC4-5, CTF18, CTF8/DSCC1, POLA2, PRIM1, and CST were codon-optimised for overexpression in insect cells and synthesised by Twist Biosciences. The details of the affinity tagging strategy and expression system used for each factor is detailed in Supplementary table S1. Plasmids for the following constructs were acquired from external sources: Polymerase δ (a kind gift from Dr Samir Hamdan)^63^, POLA1 and PRIM2 (a kind gift from Dr Joe Yeeles)^28^, RPA (a kind gift from Dr Marc Wold)^64^, PCNA (a kind gift from Dr Andrew Deans)^65^, BLM (a kind gift from Dr Petr Cejka)^66^, shelterin and variants (a kind gift from Prof. Sebastian Guettler) and Lac repressor (a kind gift from Dr John Diffley). The mutants indicated for POT1, TRF1, and TRF2, were generated using PCR-based mutagenesis using starting plasmids that were a kind gift from Dr Oviya Inian and Prof. Sebastian Guettler. A construct for the generation of biotinylated PCNA was generated as previously described^67^.

### Preparation of DNA templates for *in vitro* replication

Preparation of the 8.8 kb template containing 1.5 kb of telomeric DNA was based on a protocol developed in the laboratory of Lars Nordenskiöld^68^. Briefly plasmids were transformed into SURE2 *Escherichia coli* cells grown at 30°C. Colonies were inoculated onto LB-agar and an 8 ml small-scale culture supplemented with Ampicillin (50 µg/ml) and grown for 14-18 h. The presence of the telomeric DNA was verified by restriction endonuclease digestion with EcoRI-HF and PstI-HF (NEB) and new small-scale cultures were inoculated from the streaked colonies and grown for 6-8 h before inoculating 10 L of bacteria with a verified clone. The presence of telomeric DNA in the final large-scale cultures was verified. For all other DNA constructs used for *in vitro* replication reactions, plasmids were amplified using SURE2 *E.coli* cells grown at 30°C. Plasmids were isolated by using a QIAGEN Plasmid Maxi or Giga kit.

Forked replication templates were generated as previously described^69^, with minor modifications. Plasmid DNA (200 µg) was digested with 600 U of restriction enzyme (EcoRV-HF for ‘long’ control and telomeric constructs and NheI-HF for ‘short’ control, telomeric and LacO constructs) in a final volume of 800 µl for 2 h at 37°C. QuickCIP (NEB) was added to the sample (150 U) and incubated for 1 h at 37°C. Reactions were quenched by addition of 0.2% SDS, 25 mM EDTA pH 8.0, and 0.2 mg/ml Proteinase K and incubated for 20 min at 37°C. DNA was extracted by adding an equal volume of phenol:chloroform:isoamyl alcohol (25:24:1) (Invitrogen, 15593031) and centrifuged (20,000 *g*, 5 min). The aqueous phase was collected, and the DNA precipitated by addition of NaCl to a final concentration of 300 mM and 2.5 volumes of -20°C ethanol. Precipitated DNA was pelleted by centrifugation (20,000 *g*, 20 min, 4°C). The pellet was washed with room temperature 70% ethanol (v/v), centrifuged (20,000 *g*, 5 min) and air-dried prior to resuspension in 200 µl of TE buffer (10 mM Tris-HCl pH 8.0, 1 mM EDTA). The resulting linearised DNA was subsequently digested with 600 U of restriction enzyme (NEB, HindIII-HF for ‘long’ control and telomeric C-rich^Lead^, NotI-HF for ‘long’ control and telomeric G-rich^Lead^, and PstI-HF for ‘short’ control, telomeric, and LacO) in 1X rCutSmart buffer (NEB, B6004) in a final volume of 800 µl for 3 h at 37°C. The reaction was quenched, and the DNA extracted by phenol-chloroform followed by ethanol precipitation. The resulting pellet was resuspended in 100 µl TE buffer.

Oligonucleotides for generating the forked DNA end were adapted from a previously published system^70^ (see supplementary table S2 for sequences). Oligonucleotides were heated to 95°C for 5 min in TE buffer supplemented with 100 mM NaCl at a ratio of 3:1 and cooled to room temperature at a rate of -1°C/min to facilitate annealing. A ten-fold molar excess of the annealed fork was ligated to 60 µg of linearised template with 15,000 U T4 DNA Ligase (NEB) in 1X T4 DNA Ligase buffer (NEB) in a final volume of 600 µl and incubated for 2 h at 37°C. The reaction was quenched and the DNA purified by phenol-chloroform extraction and ethanol precipitation, as above. The resulting pellet was resuspended in 100 µl of TE buffer. Template was digested with 150 U of restriction enzyme (NEB, EcoRV-HF for ‘long’ control and telomeric constructs and NheI-HF for ‘short’ control, telomeric and LacO constructs) in a final volume of 200 µl for 3 h at 37°C. The reaction was quenched and the DNA purified by phenol-chloroform extraction and ethanol precipitation, as above. The resulting pellet was resuspended in 100 µl of TE buffer. Excess unligated fork was removed by size exclusion chromatography over a 10 ml bed volume of SP Sepharose Fast flow (GE Healthcare, 17-0729-10) equilibrated with TE + 100 mM NaCl and fractions of 200 µl were collected and analysed by agarose electrophoresis with SYBRGold. Peak fractions were pooled, supplemented with 500 mM NaCl and an equal volume of isopropanol and the DNA precipitated by centrifugation (20,000 *g*, 20 min), washed twice with 70% ethanol (v/v), air-dried and resuspended in TE to approximately 400-800 ng/µl.

### Protein expression

For expression in insect cells, plasmids were transposed into DH10MultiBac competent *E. coli* cells and grown in LB medium overnight at 37°C. Bacmid DNA was extracted and used to transfect *Spodoptera frugiperda* (Sf9) insect cells. The virus was amplified in Sf9 cells grown at 27°C, shaking at 130 rpm until passage 3 (P3), which was used to infect Hi5 suspension cultures at a density of 1-2 x 10^6^ cells/ml, grown at 27°C with shaking at 130 rpm. The following proteins were expressed by co-infection with multiple viruses: CMG (MCM5-6-7-Psf2, MCM2-4-Sld5-Psf1-3, MCM3), RFC1-5 (RFC1, RFC2-3, RFC4-5), CTF18-RFC (CTF18, RFC2-3, RFC4-5, CTF8-DSCC1), Pol α/prim (POLA1, POLA2, PRIM1, PRIM2), POT1/TPP1, CST-Pol α/prim co-complex (CST, POLA1, POLA2, PRIM1, PRIM2). For polymerase δ expression, Sf9 cells were used. Cells were harvested 72 h post-infection by centrifugation (760 *g*, 10 min). Cell pellets were washed in PBS, transferred to 50 ml Falcon tubes, and pelleted again by centrifugation (1,500 *g*, 10 min). Pellets were flash frozen in liquid nitrogen and stored at -80°C.

For bacterial expression, plasmids transformed into *E. coli* strain BL21 (DE3) were grown at 37°C with shaking at 180 rpm and induced with isopropyl β-D-thiogalactopyranoside (IPTG) as previously described for PCNA^65^, RPA^71^, Biotin-PCNA^67^. Cells were harvested by centrifugation (3500 *g*, 20 min). Cell pellets were washed with PBS, transferred to 50 ml Falcon tubes, pelleted again by centrifugation (3500 *g*, 20 min). Supernatants were discarded, the pellets flash frozen in liquid nitrogen to be stored at -80°C.

For expression in budding yeast, strains ySP53 or ySP54 were grown in YP + 2% raffinose at 30°C, to a density of 2–3 × 10^7^ cells / ml. Galactose (2%) was added for 3 h to induce protein expression. Cells were collected by centrifugation, washed once in that protein’s lysis buffer without protease inhibitors and resuspended in 0.3 - 0.4 volumes of lysis buffer + protease inhibitors (Roche, 11836170001, PMSF, AEBSF and pepstatin A). The cell suspension was frozen dropwise in liquid nitrogen and the resulting yeast popcorn was crushed in a SPEX CertiPrep 6850 Freezer/Mill (3x 2 min cycles at a crushing rate 15) and the powder stored at −80°C until required.

### Protein purification

#### CMG

CMG was purified as described^72^.

### Polymerase ε

Cells from 2 L of expression were resuspended in a 2X hypotonic lysis buffer (50 mM HEPES-KOH pH 7.6, 2mM EDTA, 2 mM DTT) supplemented with protease inhibitors (cOmplete, EDTA-free). Cold glycerol was added to a final concentration of 16.67% from a 50% stock followed by cold KCl to a final concentration of 300 mM. The lysate was incubated for 30 min, 4°C, stirring and insoluble material removed by centrifugation (35,000 *g*, 30 min, 4°C). Solid ammonium sulphate (0.16 g/ml) was added to the lysate and incubated for 30 min, at 4°C, stirring. Insoluble material was removed by centrifugation (35,000 *g*, 30 min, 4°C). Additional solid ammonium sulphate (0.3 g/ml final) was added to the supernatant and incubated for 30 min, at 4°C, stirring. Precipitated protein was collected by centrifugation (35,000 *g*, 30 min, 4°C) and resuspended in buffer A (25 mM HEPES-KOH pH 7.6, 10% glycerol, 1 mM EDTA, 1 mM DTT, 0.005% NP-40). FLAG M2 affinity gel (2 ml) was added to the lysate and incubated for 2 h, 4°C, rotating. Resin was collected in a 20 ml column and washed with 10 CV of buffer A + 300 mM KCl. Polymerase ε was eluted with 1 CV buffer A + 300 mM KCl + 0.2 mg/ml 3x FLAG peptide after incubating for 10 min, 2 CV buffer A + 300 mM KCl + 0.1 mg/ml 3x FLAG peptide after incubating for 5 min, and 3 CV of buffer A + 300 mM KCl. Eluates were pooled, diluted to 150 mM KCl, and applied to a 1 ml HiTrap Heparin HP column (GE Healthcare) equilibrated in buffer B (25 mM HEPES-KOH pH 7.6, 10% glycerol, 1 mM EDTA, 1 mM DTT, 0.01% NP-40) + 150 mM NaCl. The column was washed with 10 CV of buffer B + 150 mM NaCl and polymerase ε eluted over a 20 CV gradient from 150 mM to 1,000 mM NaCl. Peak fractions were pooled, diluted to 150 mM NaCl, and applied to a 1 ml MonoQ 5/50 GL column (GE Healthcare) equilibrated in buffer B + 150 mM NaCl. The column was washed with 10 CV of buffer B + 150 mM NaCl and polymerase ε eluted over a 20 CV gradient from 150 mM to 1,000 mM NaCl. Peak fractions were pooled and concentrated with an Amicon Ultra-15 30 kDa MWCO concentrator and applied onto a Superdex 200 Increase 10/300 GL column (GE healthcare) equilibrated with buffer C (25 mM HEPES-KOH pH 7.6, 400 mM potassium glutamate, 10% glycerol, 1 mM EDTA, 1 mM DTT, 0.01% NP-40). Peak fractions were pooled and concentrated with an Amicon Ultra-4 30 kDa MWCO concentrator, aliquoted, frozen in liquid nitrogen, and stored at -80°C.

### RPA

RPA was purified as described with some minor alterations^73^. Cells from 4 L of expression were resuspended in lysis buffer (25 mM Tris-HCl pH 7.5, 1 M NaCl, 10% glycerol, 1 mM EDTA, 1 mM DTT) at 5 ml per 1 g of pellet supplemented with protease inhibitors (Leupeptin 10 µg/ml (Discovery fine chemicals), AEBSF 0.5 mM (Melford labs), Pepstatin 10µg/ml (Discovery fine chemicals)). Lysozyme was added to 0.5 mg/ml and incubated on ice for 15 min. Cells were lysed via sonication (38%, 2 s on/4 s off, total 2 min) and insoluble material removed by centrifugation (14,000 *g*, 30 min, 4°C). The lysate was diluted to 500 mM NaCl and incubated with Affi-Gel Blue (BioRad, 1537301) resin (4 ml) for 1 h, 4°C, rotating. Resin was collected, washed with 10 CV of buffer A (25 mM Tris-HCl pH 7.5, 10% glycerol, 1 mM EDTA, 1 mM DTT) + 500 mM NaCl and 5 CV of buffer A + 800 mM NaCl. RPA was eluted with 8 CV of buffer A + 2.5 M NaCl + 30% ethylene glycol and diluted immediately to 500 mM NaCl. Pooled eluates were incubated with hydrated and cleared ssDNA-cellulose (Cell Systems, LS01132) resin (1.5 ml) for 1 h, 4°C, rotating. Resin was collected in a 20 ml column and washed with 10 CV of buffer A + 500 mM NaCl. RPA was eluted in 6 CV of buffer A + 1.5 M NaCl and 50% ethylene glycol and immediately diluted to 150 mM NaCl. Eluates were pooled and applied to a 1 ml MonoQ 5/50 GL column equilibrated in buffer A + 150 mM NaCl. The column was washed with 10 CV of buffer A + 150 mM NaCl and RPA eluted over a 30 CV gradient from 150 mM to 1,000 mM NaCl. Peak fractions were pooled and concentrated by dialysis for 2-4 h against 1 L of dialysis buffer (25 mM Tris-HCl pH 7.5, 150 mM NaCl, 40% glycerol, 1 mM EDTA, 1 mM DTT). RPA was aliquoted, frozen in liquid nitrogen, and stored at -80°C.

### Claspin

Claspin was purified as described with some minor alterations^72^. Cells from 1 L of expression were resuspended in a 2X hypotonic lysis buffer (100 mM Tris-HCl pH 8.0, 2 mM DTT) supplemented with protease inhibitors (cOmplete, EDTA-free). Cold glycerol was added to a final concentration of 16.67% from a 50% stock followed by cold NaCl to a final concentration of 400 mM. The lysate was incubated for 30 min, 4°C, stirring and insoluble material removed by centrifugation (35,000 *g*, 30 min, 4°C). The lysate was applied to 1 ml Streptactin XT superflow high-capacity resin and washed with 10 CV of buffer A (50 mM Tris-HCl pH 8.0, 10% glycerol, 1 mM DTT, 0.005% Tween-20) + 400 mM NaCl. The column was further washed with 10 CV of Buffer A + 400 mM NaCl and 5 mM Mg(OAc)_2_ + 0.5 mM ATP, and a further 10 CV wash with Buffer A + 400 mM NaCl. Claspin was eluted with 0.5 CV of buffer A + 400 mM NaCl and 30 mM biotin, followed by 1 CV after incubating for 10 min, and then 6 CV. Eluates were pooled, diluted to 100 mM NaCl and applied to a 1 ml MonoQ 5/50 GL column equilibrated in buffer A + 100 mM NaCl. The column was washed with 10 CV of buffer A + 100 mM NaCl and Claspin was eluted over a 20 CV gradient from 100 mM to 750 mM NaCl. Peak fractions were pooled and concentrated with an Amicon Ultra-15 30 kDa MWCO concentrator and applied onto a Superose 6 Increase 10/300 GL column (GE healthcare) equilibrated with Buffer B (40 mM Tris-HCl pH 8.0, 150 mM NaCl, 10% glycerol, 1 mM DTT, 0.005% Tween-20). Peak fractions were pooled, concentrated in an Amicon Ultra-4 30 kDa MWCO concentrator, aliquoted, frozen in liquid nitrogen, and stored at -80°C.

### Tipin-Timeless

Cells from 1 L of expression were resuspended in a 2X hypotonic lysis buffer (50 mM HEPES-KOH pH 7.6, 4 mM EDTA, 2 mM DTT) supplemented with protease inhibitors (cOmplete, EDTA-free). Cold glycerol was added to a final concentration of 16.67% from a 50% stock followed by cold NaCl to a final concentration of 300 mM. The lysate was incubated for 30 min, 4°C, stirring and insoluble material removed by centrifugation (35,000 *g*, 30 min, 4°C). FLAG M2 affinity gel (2 ml) was added to the lysate and incubated for 2 h, 4°C, rotating. Resin was collected in a 20 ml column and washed with 15 CV of buffer A (25 mM HEPES-KOH pH 7.6, 10% glycerol, 1 mM EDTA, 1 mM DTT, 0.01% NP-40) + 300 mM NaCl. Tipin-Timeless was eluted with 1 CV buffer A + 300 mM NaCl + 0.2 mg/ml 3x FLAG peptide after incubating for 10 min, 2 CV buffer A + 300 mM NaCl + 0.1 mg/ml 3x FLAG peptide after incubating for 5 min, and 3 CV of buffer A + 300 mM NaCl. Eluates were pooled, diluted to 100 mM NaCl and applied to a 1 ml MonoQ 5/50 GL column equilibrated in buffer A + 100 mM NaCl. The column was washed with 10 CV of buffer A + 100 mM NaCl and Tipin-Timeless was eluted over a 20 CV gradient from 100 mM to 1,000 mM NaCl. Peak fractions were pooled, diluted to 100 mM NaCl and applied to a 1 ml MonoS 5/50 GL column (GE Healthcare) equilibrated in buffer A + 100 mM NaCl. The column was washed with 10 CV of buffer A + 100 mM NaCl and Tipin-Timeless was eluted over a 20 CV gradient from 100 mM to 1,000 mM NaCl. Peak fractions were pooled, concentrated, and buffer exchanged in buffer A + 100 mM NaCl in an Amicon Ultra-4 30 kDa MWCO concentrator. The concentrated protein was aliquoted, frozen in liquid nitrogen, and stored at -80°C.

### AND-1

AND-1 was purified as described with some minor alterations^72^. Cells from 1 L of expression were resuspended in a 2X hypotonic lysis buffer (50 mM Tris-HCl pH 7.2, 2 mM DTT) supplemented with protease inhibitors (cOmplete, EDTA-free). Cold glycerol was added to a final concentration of 16.67% from a 50% stock followed by cold NaCl to a final concentration of 300 mM. The lysate was incubated for 30 min, 4°C, stirring and insoluble material removed by centrifugation (35,000 *g*, 30 min, 4°C). The lysate was applied to Streptactin XT superflow high-capacity resin (1 ml) in a 20 ml column and washed with 10 CV of buffer A (25 mM Tris-HCl pH 7.2, 10% glycerol, 1 mM DTT, 0.02% NP-40) + 300 mM NaCl. The column was further washed with 10 CV of Buffer A + 300 mM NaCl and 5 mM Mg(OAc)_2_ and 0.5 mM ATP, and a further 10 CV wash with Buffer A + 300 mM NaCl. AND-1 was eluted with 0.5 CV of buffer A + 300 mM NaCl + 30 mM biotin, followed by 1 CV after incubating for 10 min, and then 6 CV. Eluates were pooled, diluted to 150 mM NaCl and applied to a 1 ml MonoQ 5/50 GL column equilibrated in buffer A + 150 mM NaCl. The column was washed with 10 CV of buffer A + 150 mM NaCl and AND-1 was eluted over a 20 CV gradient from 150 mM to 1,000 mM NaCl. Peak fractions were pooled and concentrated with an Amicon Ultra-15 30 kDa MWCO concentrator and applied onto a Superose 6 Increase 10/300 GL column equilibrated with Buffer B (25 mM Tris-HCl pH 7.2, 150 mM NaCl, 10% glycerol, 1 mM DTT, 0.02% NP-40). Peak fractions were pooled, concentrated in an Amicon Ultra-4 30 kDa MWCO concentrator, aliquoted, frozen in liquid nitrogen, and stored at -80°C.

### CTF18-RFC

Cells from 1 L of expression were resuspended in a 2X hypotonic lysis buffer (50 mM HEPES-KOH pH 7.6, 2 mM DTT) supplemented with protease inhibitors (cOmplete, EDTA-free). Cold glycerol was added to a final concentration of 16.67% from a 50% stock followed by cold NaCl to a final concentration of 300 mM. The lysate was incubated for 30 min, 4°C, stirring and insoluble material removed by centrifugation (35,000 *g*, 30 min, 4°C). The lysate was applied to Streptactin XT superflow high-capacity resin (1 ml) in a 20 ml column and washed with 20 CV of buffer A (25 mM HEPES-KOH pH 7.6, 10% glycerol, 1 mM EDTA, 1 mM DTT, 0.02% NP-40) + 300 mM NaCl. CTF18-RFC was eluted with 0.5 CV of buffer A + 300 mM NaCl + 30 mM biotin, followed by 1 CV after incubating for 10 min, and then 6 CV. Pooled eluates were diluted to 100 mM NaCl and incubated with hydrated and cleared ssDNA-cellulose resin (0.5 ml) for 1 h, 4°C, rotating. Resin was collected in a 10 ml column and washed with 20 CV buffer A + 100 mM NaCl and CTF18-RFC was eluted with 10 CV buffer A + 300 mM NaCl. Eluates were pooled, diluted to 150 mM NaCl and applied to a 1 ml MonoQ 5/50 GL column equilibrated in buffer A + 150 mM NaCl. The column was washed with 10 CV of buffer A + 150 mM NaCl and CTF18-RFC was eluted over a 20 CV gradient from 150 mM to 1,000 mM NaCl. Peak fractions were pooled, concentrated, and buffer exchanged in buffer B (25 mM HEPES-KOH pH 7.6, 300 mM KOAc, 10% glycerol, 1 mM DTT, 0.005% Tween-20) in an Amicon Ultra-4 30 kDa MWCO concentrator. The concentrated protein was aliquoted, frozen in liquid nitrogen, and stored at -80°C.

### PCNA

PCNA was purified as described with some minor alterations^74^. Cells from 1 L of expression were resuspended in lysis buffer (50 mM Tris-HCl pH 8.0, 100 mM NaCl, 10% glycerol, 1 mM EDTA) at 5 ml per 1 g of pellet supplemented with protease inhibitors (Leupeptin 10 µg/ml, AEBSF 0.5 mM, Pepstatin 10µg/ml). Lysozyme was added to 0.5 mg/ml and incubated on ice for 15 min. Cells were lysed via sonication (38%, 2 s on/4 s off, total 2 min) and insoluble material removed by centrifugation (14,000 *g*, 30 min, 4°C). Filtered lysate was applied to a 5 ml HiTrap Q HP column (GE Healthcare) equilibrated with Buffer A (20 mM Tris-HCl pH 8.0, 10% glycerol) + 100 mM NaCl. The column was washed with 15 CV of buffer A + 100 mM NaCl. PCNA was eluted over a 5 CV gradient from 100 mM to 300 mM NaCl, followed by 3 CV at 300 mM NaCl, then 3 CV from 300 mM to 600 mM NaCl, then 5 CV from 600-700 mM NaCl, and 5 CV from 700 mM to 1,000 mM NaCl. Peak fractions were pooled and dialysed for 2 h against 2L of dialysis buffer (30 mM NaOAc pH 5.0, 5 mM NaCl, 10% glycerol). PCNA was applied to a 5 ml HiTrap Heparin HP column (GE Healthcare) equilibrated with Buffer B (30 mM NaOAc pH 5.0, 10% glycerol) + 5 mM NaCl. The column was washed with 5 CV of buffer B + 5 mM NaCl. PCNA was eluted over a 10 CV gradient from 5 mM to 1,000 mM NaCl. Peak fractions were pooled and concentrated with an Amicon Ultra-15 10 kDa MWCO concentrator and applied onto a HiPrep 26/600 Sephacryl S-200 HR column (GE healthcare) equilibrated with buffer C (25 mM Tris-HCl pH 7.5, 25 mM NaCl, 10% glycerol, 0.5 mM EDTA). Peak fractions were pooled and concentrated with an Amicon Ultra-4 10 kDa MWCO concentrator aliquoted into 500 µl fractions, frozen in liquid nitrogen, and stored at -80°C. A 500 µl aliquot was then applied to a 1 ml MonoQ 5/50 GL column (GE Healthcare) equilibrated in buffer A + 100 mM NaCl. The column was washed with 10 CV of buffer A + 100 mM NaCl and PCNA eluted over a 30 CV gradient from 100 mM to 1,000 mM NaCl. Peak fractions were pooled, concentrated, and buffer exchanged in buffer C in an Amicon Ultra-4 10 kDa MWCO concentrator. The concentrated protein was aliquoted, frozen in liquid nitrogen, and stored at -80°C.

### Pol α/prim

Pol α/prim was purified described with some minor adjustments^28^. Cells from 1 L of expression were resuspended in a 2X hypotonic lysis buffer (50 mM Tris-HCl pH 7.2, 2 mM DTT) supplemented with protease inhibitors (cOmplete, EDTA-free). Cold glycerol was added to a final concentration of 16.67% from a 50% stock followed by cold NaCl to a final concentration of 300 mM. The lysate was incubated for 30 min, 4°C, stirring and insoluble material removed by centrifugation (35,000 *g*, 30 min, 4°C). The lysate was applied to Streptactin XT superflow high-capacity resin (3 ml) in a 20 ml column and washed with 10 CV of buffer A (25 mM Tris-HCl pH 7.2, 10% glycerol, 1 mM DTT, 0.02% NP-40) + 300 mM NaCl and 10 CV of buffer A + 100 mM NaCl. Pol α/prim was eluted with 0.5 CV of buffer A + 100 mM NaCl + 30 mM biotin, followed by 1 CV after incubating for 10 min, and then 6 CV. Pooled eluates were applied to a 1 ml MonoQ 5/50 GL column equilibrated in buffer A + 100 mM NaCl. The column was washed with 10 CV of buffer A + 100 mM NaCl and polymerase α was eluted over a 20 CV gradient from 100 mM to 1,000 mM NaCl. Peak fractions were pooled and concentrated with an Amicon Ultra-15 30 kDa MWCO concentrator and applied onto a Superdex 200 Increase 10/300 GL column equilibrated with Buffer B (25 mM Tris-HCl pH 7.2, 150 mM NaCl, 10% glycerol, 1 mM DTT, 0.005% Tween-20). Peak fractions were pooled, concentrated in an Amicon Ultra-4 30 kDa MWCO concentrator, aliquoted, frozen in liquid nitrogen, and stored at -80°C.

### Biotin-PCNA Agarose

Biotin-PCNA was purified as described with minor alterations^75^. Cells from 1 L of expression were resuspended in lysis buffer (50 mM Tris-HCl pH 7.5, 80 mM NaCl, 5% glycerol, 1 mM EDTA, 10 mM 2-mercaptoethanol, 1% NP-40, 20 mM imidazole pH 7.5) at 5 ml per 1 g of pellet supplemented with protease inhibitors (Leupeptin 10 µg/ml, AEBSF 0.5 mM, Pepstatin 10µg/ml). Lysozyme was added to 2 mg/ml and incubated on ice for 15 min. Cells were lysed via sonication (38%, 2 s on/4 s off, total 2 min) and insoluble material removed by centrifugation (14,000 *g*, 30 min, 4°C). The lysate was applied onto a 5 ml HisTrap HP column (GE Healthcare) equilibrated with Buffer A (50 mM Tris-HCl pH 7.5, 500 mM NaCl, 5% glycerol, 10 mM β-mercaptoethanol) + 20 mM imidazole pH 7.5. The column was washed with 15CV of buffer A + 20 mM imidazole pH 7.5 and Biotin-PCNA was eluted over a 15 CV gradient from 20 mM to 500 mM imidazole pH 7.5. Peak fractions were pooled and aliquoted in 4 ml tubes, frozen in liquid nitrogen, and stored at -80°C. To make the biotin-PCNA Agarose column, 8 mg of protein was incubated with 2 ml of Neutravidin beads (Fisher Scientific, 11805845) for 2 h, 4°C, rotating. Resin was collected in a 20 ml column and washed with 15 CV of buffer A. Resin was resuspended in 1 CV of buffer A to generate the PCNA-agarose resin for use in polymerase δ purification.

### Polymerase δ

Polymerase δ was purified as described^67^ with minor adjustments. Cells from 2 L of expression were resuspended in lysis buffer (50 mM Tris-HCl pH 7.5, 500 mM NaCl, 10% glycerol, 5 mM β-mercaptoethanol, 0.2% NP-40) supplemented with protease inhibitors (cOmplete, EDTA-free) and 10 U/ml Benzonase (Millipore, E1014) at 3-5 ml per 1 g of pellet. Cells were lysed via sonication (30%, 2 s on/6 s off, total 2 x 3 min) and insoluble material removed by ultracentrifugation (150,000 *g*, 1 h, 4°C). The lysate was adjusted to 40 mM imidazole pH 7.5. The lysate was applied onto a 5 ml HisTrap HP column equilibrated with Buffer A (50 mM Tris-HCl pH 7.5, 80 mM NaCl, 10% glycerol, 5 mM β-mercaptoethanol) + 40 mM imidazole pH 7.5. The column was washed with 10CV of buffer A + 40 mM imidazole pH 7.5 and polymerase δ eluted over a 10 CV gradient from 40 mM to 500 mM imidazole pH 7.5. Peak fractions were pooled and incubated with PCNA-Agarose resin (1 ml of resin per 5 ml of eluate) pre-equilibrated with buffer B (50 mM Tris-HCl pH 7.5, 5% glycerol, 10 mM β-mercaptoethanol) + 100 mM NaCl overnight at 4°C, rotating. Resin was collected in a 20 ml column and washed with 10 CV of buffer B + 100 mM NaCl and 10 CV of buffer B + 200 mM NaCl. Proteins were eluted with 10 CV of buffer B + 1.2 M NaCl. Pooled eluates were concentrated with a Vivaspin6 10 kDa MWCO concentrator and applied onto a Superdex 200 Increase 10/300 GL column equilibrated with Buffer C (50 mM Tris-HCl pH 7.5, 150 mM NaCl, 5% glycerol, 1 mM DTT). Peak fractions were pooled, concentrated Vivaspin6 10 kDa MWCO concentrator aliquoted, frozen in liquid nitrogen, and stored at -80°C.

### RFC1-5

Cells from 1 L of expression were resuspended in a 2X hypotonic lysis buffer (50 mM HEPES-KOH pH 7.6, 2 mM DTT) supplemented with protease inhibitors (cOmplete, EDTA-free). Cold glycerol was added to a final concentration of 16.67% from a 50% stock followed by cold NaCl to a final concentration of 300 mM. The lysate was incubated for 30 min, 4°C, stirring and insoluble material removed by centrifugation (35,000 *g*, 30 min, 4°C). The lysate was applied to Streptactin XT superflow high-capacity resin (1 ml) in a 20 ml column and washed with 20 CV of buffer A (25 mM HEPES-KOH pH 7.6, 10% glycerol, 1 mM EDTA, 1 mM DTT, 0.02% NP-40) + 300 mM NaCl. RFC1-5 was eluted with 0.5 CV of buffer A + 300 mM NaCl + 30 mM biotin, followed by 1 CV after incubating for 10 min, and then 6 CV. Pooled eluates were incubated with hydrated and cleared ssDNA-cellulose resin (0.5 ml) for 1 h, 4°C, rotating. Resin was collected in a 10 ml column and washed with 20 CV buffer A + 300 mM NaCl and RFC1-5 was eluted with 10 CV buffer A + 500 mM NaCl. Eluates were pooled, diluted to 150 mM NaCl and applied to a 1 ml MonoQ 5/50 GL column equilibrated in buffer A + 150 mM NaCl. The column was washed with 10 CV of buffer A + 150 mM NaCl and RFC1-5 was eluted over a 20 CV gradient from 150 mM to 1,000 mM NaCl. Peak fractions were pooled, concentrated, and buffer exchanged in buffer B (25 mM HEPES-KOH pH 7.6, 300 mM KOAc, 10% glycerol, 1 mM DTT, 0.005% Tween-20) in an Amicon Ultra-4 30 kDa MWCO concentrator. The concentrated protein was aliquoted, frozen in liquid nitrogen, and stored at -80°C.

### FEN1 and LIG1

Frozen cell powder (from 12l ySP53 culture) was thawed and resuspended in FEN1 lysis buffer (25 mM Tris-Cl pH 7.2, 0.02% NP40, 10% glycerol, 0.5M NaCl, 1 mM DTT, protease inhibitors). Insoluble material was removed by centrifugation (235,000 x g, 4°C, 1 h) and the supernatant mixed with 2 mL anti-FLAG M2 affinity resin (Sigma-Aldrich) at 4°C for 90 min. The resin was collected in a disposable column and washed extensively in FEN1 lysis buffer without protease inhibitors, then FEN1 lysis buffer + 5 mM MgOAc + 1 mM ATP. FEN1 was eluted by incubating the resin with 2.5 ml of FEN1 lysis buffer + 0.5 mg/mL 3FLAG peptide for 30 min followed by 2.5 ml of FEN1 lysis buffer + 0.25 mg/mL 3FLAG peptide. The eluate was concentrated to 500 µl by centrifugation through a AMICON Ultra 15 10 KDa cutoff (Merck) and then loaded onto a 24 ml Superdex 200 gel filtration column equilibrated in 25 mM Tris-Cl pH 7.2, 0.02% NP40, 10% glycerol, 0.3M NaOAc, 1 mM DTT. Peak fractions containing FEN1 were pooled, aliquoted and snap froze. LIG1 was purified from strain ySP54 in the same way except that Hepes-KOH pH 7.6 was used in place of Tris-Cl pH 7.2 in the lysis and gel filtration buffers.

### Wild type and helicase inactive BLM

BLM was purified as described with some minor alterations^66^. Cells from 1 L of expression were resuspended in a 2X hypotonic lysis buffer (100 mM Tris-HCl pH 7.5, 2 mM EDTA, 2 mM DTT) protease inhibitors (cOmplete, EDTA-free). Cold glycerol was added to a final concentration of 16.67% from a 50% stock followed by cold NaCl to a final concentration of 300 mM. The lysate was incubated for 30 min, 4°C, stirring and insoluble material removed by centrifugation (35,000 *g*, 30 min, 4°C). Amylose resin (NEB, E8021) (4 ml) was added to the lysate and incubated for 1 h, 4°C, rotating. Resin was collected in a 20 ml column and washed with 15 CV of buffer A (50 mM Tris-HCl pH 7.5, 10% glycerol, 1 mM DTT) + 1 M NaCl. BLM was eluted with 10 CV of buffer A + 300 mM NaCl + 10 mM maltose. Eluates were pooled and incubated with 250 nM PreScission protease (GE Healthcare, 27084301) for 1 h, 4°C, rotating. Eluates were then supplemented with 10 mM imidazole pH 7.5 before incubating with Ni-NTA resin (QIAGEN, 30210) (1 ml) for 1 h, 4°C, rotating. Resin was collected in a 20 ml column and washed with 20 CV of buffer A + 1 M NaCl + 58 mM imidazole pH 7.5 followed by 5 CV of buffer A + 150 mM NaCl + 58 mM imidazole pH 7.5. BLM was eluted with 10 CV of buffer A + 100 mM NaCl + 300 mM imidazole pH 7.5. Eluates were pooled and concentrated by dialysis for 2 h against 1 L of buffer B (50 mM Tris-HCl pH 7.5, 100 mM NaCl 40% glycerol, 1 mM DTT). BLM was aliquoted, frozen in liquid nitrogen, and stored at -80°C.

### Shelterin wildtype and variants

Shelterin and shelterin variants were purified as described for the full complex, with some minor modifications^76^. Cells from 1 L of expression were resuspended in a 2X hypotonic lysis buffer (100 mM HEPES-KOH pH 7.6, 2 mM DTT) supplemented with protease inhibitors (cOmplete, EDTA-free). Cold glycerol was added to a final concentration of 16.67% from a 50% stock followed by cold NaCl to a final concentration of 300 mM. The lysate was incubated for 30 min, 4°C, stirring and insoluble material removed by centrifugation (35,000 *g*, 30 min, 4°C). The lysate was applied to Streptactin XT superflow high-capacity resin (1 ml) in a 20 ml column and washed with 30 CV of buffer A (50 mM HEPES-KOH pH 7.6, 300 mM NaCl, 10% glycerol, 1 mM DTT). Shelterin was eluted with 0.5 CV of buffer A + 30 mM biotin, followed by 1 CV after incubating for 10 min, and then 6 CV. The most concentrated eluate was applied onto a Superose 6 Increase 10/300 GL column equilibrated with Buffer A except for TPP1/POT1 complex, which was run over a Superdex 200 Increase 10/300 GL column equilibrated with Buffer A. Peak fractions were pooled, concentrated in a Vivaspin6 10 kDa MWCO concentrator, aliquoted, frozen in liquid nitrogen, and stored at -80°C.

### CST-Pol α/prim complex

Cells from 2 L of expression were resuspended in a 2X hypotonic lysis buffer (50 mM Tris-HCl pH 7.2, 2 mM DTT) supplemented with protease inhibitors (cOmplete, EDTA-free). Cold glycerol was added to a final concentration of 16.67% from a 50% stock followed by cold NaCl to a final concentration of 300 mM. The lysate was incubated for 30 min, 4°C, stirring and insoluble material removed by centrifugation (35,000 *g*, 30 min, 4°C). The lysate was applied to Streptactin XT superflow high-capacity resin (3 ml) in a 20 ml column and washed with 30 CV of buffer A (25 mM Tris-HCl pH 7.2, 300 mM NaCl, 10% glycerol, 1 mM DTT, 0.02% NP-40). CST-Pol α/prim was eluted with 0.5 CV of buffer A + 30 mM biotin, followed by 1 CV after incubating for 10 min, and then 6 CV. Eluates were pooled and incubated with FLAG M2 affinity gel (1 ml) and incubated for 2 h, 4°C, rotating. Resin was collected in a 20 ml column and washed with 15 CV of buffer A. CST-Pol α/prim was eluted with 1 CV buffer A + 0.2 mg/ml 3x FLAG peptide after incubating for 10 min, 2 CV buffer A + 0.1 mg/ml 3x FLAG peptide after incubating for 5 min, and 3 CV of buffer A. Eluates were pooled and dialysed against buffer B (25 mM Tris-HCl pH 7.2, 150 mM NaCl, 10% glycerol, 1 mM DTT, 0.02% NP-40). Peak fractions were pooled, concentrated in an Amicon Ultra-4 30 kDa MWCO concentrator, aliquoted, frozen in liquid nitrogen, and stored at -80°C.

### In vitro replication reactions

Replication assays were carried out at 37°C in three steps based on previously published work^28,69^. The final replication buffer contained 25 mM HEPES-KOH pH 7.6, 80 mM potassium glutamate, 10 mM Mg(OAc)_2_, 5% glycerol, 40 µg/ml BSA, 1 mM DTT, and 0.1 mM EDTA, with the concentration of potassium glutamate increased to 130 mM in the final initiation step for reactions containing Pol δ, Pol α/prim and RFC, except where LIG1 and FEN1 were added. Unless stated in the figure legend, the final concentrations of DNA, protein and nucleotides were: 1.5 nM DNA template, 1.2 or 0.45 nM initiating oligonucleotide (in the absence or presence of Pol δ, Pol α/prim and RFC1-5 respectively), 40 nM CMG, 20 nM Pol ε, 50 nM PCNA, 20-30 nM CTF18-RFC, 20 nM AND-1, 20 nM Claspin, 20 nM Tipin-Timeless, 50-100 nM RPA, 1.25-2.5 nM RFC, 2 nM Pol δ, 10-30 nM of Pol α/prim, 20 nM FEN1, 40 nM LIG1, 5 mM ATP, 60 µM dA/C/T/GTP, 200 µM CTP/UTP/GTP, and 66 nM α-^32^P-dCTP.

Reactions were set up as follows: the initiating oligonucleotide (see supplementary table S2 for sequence) was incubated with 3 nM template for 15 min in reaction buffer supplemented with 0.1 mM AMP-PNP. Subsequently, CMG was loaded onto the primed DNA substrate at a final concentration of 80 nM in a 5 µl reaction and incubated for 10 min. The reaction was subsequently diluted to 7.5 µl by adding reaction buffer supplemented with 240 µM dA/dCTP and 0.1 mm AMP-PNP, and a premixed solution of Pol ε, PCNA, CTF18-RFC, AND-1, Claspin, Tipin-Timeless, and Pol α/prim (in reactions competent for lagging strand synthesis) and the reaction was incubated for 2 min. For leading strand only reactions, replication was initiated by staggering the addition of 2.5 ul of reaction buffer supplemented with 20 mM ATP, 240 µM dT/GTP, 62 nM α-^32^P-dCTP, and RPA. For reaction competent for lagging strand synthesis, replication was initiated by adding 2.5 µl of reaction buffer supplemented with 20 mM ATP, 240 µM dT/GTP, 800 µM CTP/UTP/GTP, 62 nM α-^32^P-dCTP, RPA, RFC, Pol δ. For lagging strand maturation, FEN1 and LIG1 were added during the initiation step. Where indicated, BLM, shelterin, shelterin variants, LacI, CST-Pol α/prim, or phenDC3 were added in the last initiation mix or spiked in after initiation. For Figures 5b and 5c, shelterin lacking TPP1/POT1 was preincubated with wt TPP1/POT1 or TPP1/POT1 variants for 10 min. For P1 sensitivity assays, the amount of P1 nuclease (NEB, M0660) indicated in the figure legend was added 5 min before the reaction was quenched. Pulse-chase experiments were performed using the conditions for standard leading strand replication reactions, except that the concentration of dCTP was reduced to 20 µM during the pulse. 600 µM unlabelled dA/G/C/TTP was added after 50 s.

### Molecular weight markers

For molecular weight marker, 12.5 µg HindIII-digested phage lambda DNA (NEB, N3012) was dephosphorylated in 1X rCutSmart buffer (NEB, B6004) with 10 U of Quick-CIP (NEB, M0525) in a final volume of 30 µl for 1 h at 37°C. The phosphatase was inactivated at 80°C for 2 min and the DNA was purified through a Microspin G-50 column (Cytiva, 27533002) and stored at -20°C. The marker was end labelled in a 10 µl reaction with 2 µl of dephosphorylated marker, γ-^32^P-ATP (0.01 mCi), 1X PNK buffer and 6 U of PNK enzyme (NEB, M0201). The reaction was incubated for 30 min at 37°C, followed by heat inactivation of PNK at 65°C for 20 min. Unincorporated radiolabelled nucleotides were removed by passing the sample through a Microspin G-50 column and stored at -20°C.

### Processing and analysis of replication products

Reactions were quenched at the indicated time by addition of an equal volume of stop buffer containing 50 mM EDTA, 0.4% SDS, and 0.4 mg/ml Proteinase K. After 20 min at 37°C, an equal volume of phenol:chloroform:isoamyl alcohol (25:24:1) was added and the aqueous phase collected after centrifugation (14,000 *g*, 5 min). Unincorporated radiolabelled nucleotides were removed by passing the sample through a Microspin G-50 column. For nicking assays, reaction products were digested with 10 U of Nb.BsrDI (NEB) for 2 h at 37°C. For denaturing alkaline agarose electrophoresis, the samples and a molecular weight marker were supplemented with 20 mM EDTA, 50 mM NaOH, and 0.5% sucrose with xylene cyanol. Reactions were analysed by 0.7% denaturing alkaline agarose electrophoresis in a 20 x 17 cm cassette supplemented with 2 mM EDTA and 30 mM NaOH and run for 16 h at 24 V in 2 mM EDTA and 30 mM NaOH. Gels were fixed in 5% trichloroacetic acid for 30 min. For native agarose electrophoresis, the samples and a molecular weight marker were supplemented with 6X NEB Purple Loading Dye (NEB, B7024). Reactions were analysed by 0.7% agarose/TAE electrophoresis for 20 h at 30 V at 4°C in a 20 x 17 cm cassette. For 2D native-denaturing electrophoresis analysis, replication products were analysed by 0.7% native agarose electrophoresis as above, the lane excised and separated by 0.7% denaturing alkaline agarose electrophoresis. Gels were dried on filter paper before being exposed to a Storage Phosphor Screen (GE Healthcare, BAS-IP MS 2025, GE28-9564-75) and imaged on a Typhoon scanner (Cytiva). Images were analysed and quantified in ImageJ as described in the figure legends and graphs were made using Prism.

### BLM helicase assay

CLB353 (see table S2) was end labelled at a final concentration of 2 µM in a 10 µL reaction with γ-^32^P-ATP (0.02 mCi), 1X T4 Polynucleotide Kinase (PNK) buffer and 6 U of T4 PNK enzyme (NEB, M0201). The reaction was incubated for 30 min at 37°C, followed by heat inactivation of PNK for 20 min at 65°C for 20 min. Unincorporated radiolabelled nucleotides were removed by passing the sample through a G50 column. To make the forked DNA duplex, radiolabelled CLB353 and non-radiolabelled CLB354 (see table S2) were heated to 95°C for 5 min in TE buffer supplemented with 100 mM NaCl and cooled to room temperature at a rate of -1°C/min to facilitate annealing. The non-radiolabelled oligonucleotide was at a 2-fold molar excess to the radiolabelled oligo.

BLM at the indicated concentrations was incubated with 1.5 nM of labelled forked DNA duplex for 5 min at 37°C in a 10 µl reaction containing 20 mM Tris-HCl pH 7.5, 2 mM MgCl_2_, 1 mM TCEP, 0.1 mg/ml BSA. To stimulate unwinding, ATP was added to a final concentration of 5 mM along with 50 nM of unlabelled CLB353 to trap any unwound oligonucleotides and prevent substrate reannealing. After 10 min at 37°C reactions were quenched by addition of EDTA to 40 mM and SDS to 0.1% as well as xylene cyanol and bromophenol blue. Samples were analysed on 10% Novex TBE gels (ThermoFisher Scientific, LC6678) in 1X TBE (90 mM Tris-base, 90 mM Boric acid, 2.5 mM EDTA) at 90 V for 90 min. Gels were dried before being exposed to a Storage Phosphor Screen and imaged on a Typhoon scanner. Images were analysed and quantified in ImageJ.

### Immunoblotting

Recombinant shelterin and shelterin variants (1 µl) were resolved using SDS-PAGE and transferred to a nitrocellulose membrane. Western blot was performed with 5% milk in TBS containing 0.1% Tween-20. TRF1 was detected with a custom-made rabbit antibody (a kind gift from Prof. Sebastian Guettler) at 1:10,000 dilution. TRF2 was detected with antibody D1Y5D (Cell Signalling, 13136) at 1:5,000 dilution. POT1 was detected with antibody EPR6319 (Abcam, ab124784) at 1:2,000 dilution. Strep-TIN2 was detected with antibody StrepMAB-Classic (IBA lifesciences, 21507001) at 1:2,000 dilution. Primary antibodies were followed by secondary antibody goat anti-rabbit IgG (LI-COR, 926-68071), goat anti-mouse IgG (LI-COR, 926-68070), goat anti-rabbit IgG (LI-COR, 926-32211), or goat anti-mouse IgG (LI-COR, 926-32210) at 1:5,000 dilution. Immunoblots were visualised with the LI-COR Odyssey imaging system according to the manufacturer’s instructions. Images were analysed in Image Studio.

### Electrophoretic mobility shift assay

TBL297 (see table S2) was end labelled at a final concentration of 2 µM in a 10 µL reaction with γ-^32^P-ATP (0.02 mCi), 1X T4 Polynucleotide Kinase (PNK) buffer and 6 U of T4 PNK enzyme (NEB, M0201). The reaction was incubated for 30 min at 37°C, followed by heat inactivation of PNK for 20 min at 65°C for 20 min. Unincorporated radiolabelled nucleotides were removed by passing the sample through a G50 column. To make the forked DNA duplex, radiolabelled TBL297 and non-radiolabelled TBL296 (see table S2) were heated to 95°C for 5 min in TE buffer supplemented with 100 mM NaCl and cooled to room temperature at a rate of -1°C/min to facilitate annealing. The non-radiolabelled oligonucleotide was at a 1.2-fold molar excess to the radiolabelled oligo.

For the EMSA, labelled DNA duplex was incubated for 30 min on ice at a final concentration of 2 nM with the shelterin concentrations indicated in a buffer containing 20 mM HEPES-KOH pH 8 and 100 mM NaCl. Samples were supplemented with 6X NEB Purple Loading Dye, no SDS (NEB, B7025) and analysed by 1% agarose electrophoresis in 0.5X TBE at 100 V for 1.5 h in 0.5X TBE. Gels were dried on filter paper before being exposed to a Storage Phosphor Screen and imaged on a Typhoon scanner (Cytiva). Images were analysed and quantified in ImageJ.

### Electron Microscopy analysis

For EM analysis, replication assays were performed using the conditions for standard leading and lagging strand replication reactions, except radiolabelled nucleotides were omitted and each sample condition was 50 µl total. Reactions were quenched by addition of EDTA to 5 mM at the times indicated in the figure legend. For crosslinking, samples were pipetted onto parafilm, and trioxsalen (Sigma, T6137) added to a final concentration of 2 µg/ml. Samples were exposed to 365 nm UV light for 5 min using a UVP Blak Ray Lamp Model UVL-21, with 365 nm UV bulbs at 2-3 cm from the light source. Samples were returned to a 1.5 ml Eppendorf. The reaction was quenched by addition of stop buffer as described above, and the DNA obtained by phenol-chloroform extraction and ethanol precipitation. The resulting pellet was resuspended in 8 µl of TE buffer and the concentration measured by a Qubit 4 Fluorometer (ThermoFisher). For EM analysis, 5 µl of DNA in TE was mixed with 5 μl of formamide (Thermo Scientific, 17899) and 0.4 μl of 0.02% benzalkonium chloride (BAC, Sigma B6285) in TE. Immediately after mixing, the drop was allowed to slide through a freshly-cleaved mica sheet (Ted pellainc, product no: 52-6) used as a ramp inside a 15 cm dish containing 50 ml of distilled water. The monomolecular layer above the water surface was gently touched with a carbon-coated copper grid, previously placed in contact with an ethidium bromide droplet (33.3 μg/ml in H2O) for 30–45 min. Carbon grids with absorbed DNA molecules were immediately stained with a solution of uranyl acetate 0.2 μg/μL in ethanol and subjected to low-angle rotary shadowing with 8 nm of platinum using a MED020 evaporator, modified with the low-angle grid shadowing kit (Leica 16770525). TEM pictures were taken using a FEI Tecnai12 Bio twin microscope operated at 120 KV and equipped with a side-mounted GATAN Orius SC-1000 camera controlled by the Digital Micrograph software. For acquisition of large areas, overlapping fields were acquired and stitched using the Digital Micrograph software. Micrographs in DM3 format were analysed in FIJI/ImageJ software v2.0.0-rc-69/1.52p. In these conditions, in BAC spreads, a length of 0.36 μm corresponds to 1 kb of double-stranded DNA. Data annotation and storage during the analysis was performed with an Image J macro^77^

## Supporting information

Supplementary file detailing the DNA sequences and plasmid constructs used in this study

## ACKNOWLEDGEMENTS

The authors would like to thank P. Cejka, S. Hamdan, L. Nordenskiöld, A. Soman and J. Yeeles for sharing protocols and reagents. We are grateful to S. Guettler and O. Inian for sharing reagents and advice on shelterin purification. M.E.D is funded by Cancer Research UK Career Development Award (C68409/A28129).

## AUTHOR CONTRIBUTIONS

The study was conceived by M.E.D. In vitro replication reactions were performed by C.L.B, with assistance purifying replication factors from C.E.L.F and K.H. FEN1 and LIG1 were cloned and purified by T.D. EM imaging was performed by G.M. on a sample prepared by C.L.B. M.E.D and Y.D provided supervision and the manuscript was written by M.E.D with input from all authors.

## REFERENCES

1 Lim, C. J. & Cech, T. R. Shaping human telomeres: from shelterin and CST complexes to telomeric chromatin organization. Nat Rev Mol Cell Biol 22, 283–298 (2021). 10.1038/s41580-021-00328-y

2 de Lange, T. Shelterin-Mediated Telomere Protection. Annu Rev Genet 52, 223–247 (2018). 10.1146/annurev-genet-032918-021921

3 Muraki, K., Nyhan, K., Han, L. & Murnane, J. P. Mechanisms of telomere loss and their consequences for chromosome instability. Front Oncol 2, 135 (2012). 10.3389/fonc.2012.00135

4 Burgers, P. M. J. & Kunkel, T. A. Eukaryotic DNA Replication Fork. Annu Rev Biochem 86, 417–438 (2017). 10.1146/annurev-biochem-061516-044709

5 Goswami, P. et al. Structure of DNA-CMG-Pol epsilon elucidates the roles of the non-catalytic polymerase modules in the eukaryotic replisome. Nat Commun 9, 5061 (2018). 10.1038/s41467-018-07417-1

6 Jones, M. L., Aria, V., Baris, Y. & Yeeles, J. T. P. How Pol α-primase is targeted to replisomes to prime eukaryotic DNA replication. Mol Cell 83, 2911–2924.e2916 (2023). 10.1016/j.molcel.2023.06.035

7 Zeman, M. K. & Cimprich, K. A. Causes and consequences of replication stress. Nat Cell Biol 16, 2–9 (2014). 10.1038/ncb2897

8 Durkin, S. G. & Glover, T. W. Chromosome fragile sites. Annu Rev Genet 41, 169–192 (2007). 10.1146/annurev.genet.41.042007.165900

9 Sfeir, A. et al. Mammalian telomeres resemble fragile sites and require TRF1 for efficient replication. Cell 138, 90–103 (2009). 10.1016/j.cell.2009.06.021

10 Huda, A. et al. The telomerase reverse transcriptase elongates reversed replication forks at telomeric repeats. Sci Adv 9, eadf2011 (2023). 10.1126/sciadv.adf2011

11 Miller, K. M., Rog, O. & Cooper, J. P. Semi-conservative DNA replication through telomeres requires Taz1. Nature 440, 824–828 (2006). 10.1038/nature04638

12 Ivessa, A. S., Zhou, J. Q., Schulz, V. P., Monson, E. K. & Zakian, V. A. Saccharomyces Rrm3p, a 5’ to 3’ DNA helicase that promotes replication fork progression through telomeric and subtelomeric DNA. Genes Dev 16, 1383–1396 (2002).

13 Douglas, M. E. How to write an ending: Telomere replication as a multistep process. DNA Repair (Amst) 144, 103774 (2024). 10.1016/j.dnarep.2024.103774

14 Maestroni, L., Matmati, S. & Coulon, S. Solving the Telomere Replication Problem. Genes (Basel) 8 (2017). 10.3390/genes8020055

15 Tran, P. L., Mergny, J. L. & Alberti, P. Stability of telomeric G-quadruplexes. Nucleic Acids Res 39, 3282–3294 (2011). 10.1093/nar/gkq1292

16 Williams, S. L. et al. Replication-induced DNA secondary structures drive fork uncoupling and breakage. EMBO J 42, e114334 (2023). 10.15252/embj.2023114334

17 Douglas, M. E. & Diffley, J. F. X. Budding yeast Rap1, but not telomeric DNA, is inhibitory for multiple stages of DNA replication in vitro. Nucleic Acids Res 49, 5671–5683 (2021). 10.1093/nar/gkab416

18 Batra, S. et al. G-quadruplex-stalled eukaryotic replisome structure reveals helical inchworm DNA translocation. Science 387, eadt1978 (2025). 10.1126/science.adt1978

19 Drosopoulos, W. C., Kosiyatrakul, S. T. & Schildkraut, C. L. BLM helicase facilitates telomere replication during leading strand synthesis of telomeres. J Cell Biol 210, 191–208 (2015). 10.1083/jcb.201410061

20 Sun, H., Karow, J. K., Hickson, I. D. & Maizels, N. The Bloom’s syndrome helicase unwinds G4 DNA. J Biol Chem 273, 27587–27592 (1998). 10.1074/jbc.273.42.27587

21 Wu, W. Q., Hou, X. M., Li, M., Dou, S. X. & Xi, X. G. BLM unfolds G-quadruplexes in different structural environments through different mechanisms. Nucleic Acids Res 43, 4614–4626 (2015). 10.1093/nar/gkv361

22 Vannier, J. B. et al. RTEL1 is a replisome-associated helicase that promotes telomere and genome-wide replication. Science 342, 239–242 (2013). 10.1126/science.1241779

23 Zimmermann, M., Kibe, T., Kabir, S. & de Lange, T. TRF1 negotiates TTAGGG repeat-associated replication problems by recruiting the BLM helicase and the TPP1/POT1 repressor of ATR signaling. Genes Dev 28, 2477–2491 (2014). 10.1101/gad.251611.114

24 Crabbe, L., Verdun, R. E., Haggblom, C. I. & Karlseder, J. Defective telomere lagging strand synthesis in cells lacking WRN helicase activity. Science 306, 1951–1953 (2004). 10.1126/science.1103619

25 Ohki, R. & Ishikawa, F. Telomere-bound TRF1 and TRF2 stall the replication fork at telomeric repeats. Nucleic Acids Res 32, 1627–1637 (2004). 10.1093/nar/gkh309

26 Leman, A. R. et al. Timeless preserves telomere length by promoting efficient DNA replication through human telomeres. Cell Cycle 11, 2337–2347 (2012). 10.4161/cc.20810

27 Yang, Z., Sharma, K. & de Lange, T. TRF1 uses a noncanonical function of TFIIH to promote telomere replication. Genes Dev 36, 956–969 (2022). 10.1101/gad.349975.122

28 Baris, Y., Taylor, M. R. G., Aria, V. & Yeeles, J. T. P. Fast and efficient DNA replication with purified human proteins. Nature 606, 204–210 (2022). 10.1038/s41586-022-04759-1

29 Drosopoulos, W. C., Kosiyatrakul, S. T., Yan, Z., Calderano, S. G. & Schildkraut, C. L. Human telomeres replicate using chromosome-specific, rather than universal, replication programs. J Cell Biol 197, 253–266 (2012). 10.1083/jcb.201112083

30 Kumar, C., Batra, S., Griffith, J. D. & Remus, D. The interplay of RNA:DNA hybrid structure and G-quadruplexes determines the outcome of R-loop-replisome collisions. Elife 10 (2021). 10.7554/eLife.72286

31 Deegan, T. D., Baxter, J., Ortiz Bazan, M. A., Yeeles, J. T. P. & Labib, K. P. M. Pif1-Family Helicases Support Fork Convergence during DNA Replication Termination in Eukaryotes. Mol Cell 74, 231–244 e239 (2019). 10.1016/j.molcel.2019.01.040

32 Huber, M. D., Lee, D. C. & Maizels, N. G4 DNA unwinding by BLM and Sgs1p: substrate specificity and substrate-specific inhibition. Nucleic Acids Res 30, 3954–3961 (2002). 10.1093/nar/gkf530

33 Takai, K. K., Hooper, S., Blackwood, S., Gandhi, R. & de Lange, T. In vivo stoichiometry of shelterin components. J Biol Chem 285, 1457–1467 (2010). 10.1074/jbc.M109.038026

34 Janovič, T., Perez, G. I. & Schmidt, J. C. TRF1 and TRF2 form distinct shelterin subcomplexes at telomeres. bioRxiv, 2024.2012.2023.630076 (2024). 10.1101/2024.12.23.630076

35 Martínez, P. et al. Increased telomere fragility and fusions resulting from TRF1 deficiency lead to degenerative pathologies and increased cancer in mice. Genes Dev 23, 2060–2075 (2009). 10.1101/gad.543509

36 Westhorpe, R., Roske, J. J. & Yeeles, J. T. P. Mechanisms controlling replication fork stalling and collapse at topoisomerase 1 cleavage complexes. Mol Cell 84, 3469–3481.e3467 (2024). 10.1016/j.molcel.2024.08.004

37 Wu, Y., Poulos, R. C. & Reddel, R. R. Role of POT1 in Human Cancer. Cancers (Basel) 12 (2020). 10.3390/cancers12102739

38 Wu, L. et al. Pot1 deficiency initiates DNA damage checkpoint activation and aberrant homologous recombination at telomeres. Cell 126, 49–62 (2006). 10.1016/j.cell.2006.05.037

39 Huang, C., Dai, X. & Chai, W. Human Stn1 protects telomere integrity by promoting efficient lagging-strand synthesis at telomeres and mediating C-strand fill-in. Cell Res 22, 1681–1695 (2012). 10.1038/cr.2012.132

40 Kasbek, C., Wang, F. & Price, C. M. Human TEN1 maintains telomere integrity and functions in genome-wide replication restart. J Biol Chem 288, 30139–30150 (2013). 10.1074/jbc.M113.493478

41 Chen, L. Y., Majerská, J. & Lingner, J. Molecular basis of telomere syndrome caused by CTC1 mutations. Genes Dev 27, 2099–2108 (2013). 10.1101/gad.222893.113

42 Chen, L. Y., Redon, S. & Lingner, J. The human CST complex is a terminator of telomerase activity. Nature 488, 540–544 (2012). 10.1038/nature11269

43 Wan, M., Qin, J., Songyang, Z. & Liu, D. OB fold-containing protein 1 (OBFC1), a human homolog of yeast Stn1, associates with TPP1 and is implicated in telomere length regulation. J Biol Chem 284, 26725–26731 (2009). 10.1074/jbc.M109.021105

44 Cai, S. W. et al. POT1 recruits and regulates CST-Polα/primase at human telomeres. Cell 187, 3638–3651.e3618 (2024). 10.1016/j.cell.2024.05.002

45 Wu, P., Takai, H. & de Lange, T. Telomeric 3’ overhangs derive from resection by Exo1 and Apollo and fill-in by POT1b-associated CST. Cell 150, 39–52 (2012). 10.1016/j.cell.2012.05.026

46 Vannier, J. B., Pavicic-Kaltenbrunner, V., Petalcorin, M. I., Ding, H. & Boulton, S. J. RTEL1 dismantles T loops and counteracts telomeric G4-DNA to maintain telomere integrity. Cell 149, 795–806 (2012). 10.1016/j.cell.2012.03.030

47 Rizzo, A. et al. Stabilization of quadruplex DNA perturbs telomere replication leading to the activation of an ATR-dependent ATM signaling pathway. Nucleic Acids Res 37, 5353–5364 (2009). 10.1093/nar/gkp582

48 Lormand, J. D. et al. DNA polymerase δ stalls on telomeric lagging strand templates independently from G-quadruplex formation. Nucleic Acids Res 41, 10323–10333 (2013). 10.1093/nar/gkt813

49 Makovets, S., Herskowitz, I. & Blackburn, E. H. Anatomy and dynamics of DNA replication fork movement in yeast telomeric regions. Mol Cell Biol 24, 4019–4031 (2004).

50 Goto, G. H. et al. Binding of Multiple Rap1 Proteins Stimulates Chromosome Breakage Induction during DNA Replication. PLoS Genet 11, e1005283 (2015). 10.1371/journal.pgen.1005283

51 Kavlashvili, T., Liu, W., Mohamed, T. M., Cortez, D. & Dewar, J. M. Replication fork uncoupling causes nascent strand degradation and fork reversal. Nat Struct Mol Biol 30, 115–124 (2023). 10.1038/s41594-022-00871-y

52 Vaurs, M. et al. TRF1 relies on fork reversal to prevent fragility at human telomeres. Nat Commun 16, 6439 (2025). 10.1038/s41467-025-61828-5

53 Ivessa, A. S. et al. The Saccharomyces cerevisiae helicase Rrm3p facilitates replication past nonhistone protein-DNA complexes. Mol Cell 12, 1525–1536 (2003).

54 Sparks, J. L. et al. The CMG Helicase Bypasses DNA-Protein Cross-Links to Facilitate Their Repair. Cell 176, 167–181 e121 (2019). 10.1016/j.cell.2018.10.053

55 Pisano, S. et al. The human telomeric protein hTRF1 induces telomere-specific nucleosome mobility. Nucleic Acids Res 38, 2247–2255 (2010). 10.1093/nar/gkp1228

56 Galati, A. et al. The human telomeric protein TRF1 specifically recognizes nucleosomal binding sites and alters nucleosome structure. J Mol Biol 360, 377–385 (2006). 10.1016/j.jmb.2006.04.071

57 Hu, H. et al. Structural basis of telomeric nucleosome recognition by shelterin factor TRF1. Sci Adv 9, eadi4148 (2023). 10.1126/sciadv.adi4148

58 Galati, A. et al. TRF1 and TRF2 binding to telomeres is modulated by nucleosomal organization. Nucleic Acids Res 43, 5824–5837 (2015). 10.1093/nar/gkv507

59 Denchi, E. L. & de Lange, T. Protection of telomeres through independent control of ATM and ATR by TRF2 and POT1. Nature 448, 1068–1071 (2007). 10.1038/nature06065

60 Hockemeyer, D., Sfeir, A. J., Shay, J. W., Wright, W. E. & de Lange, T. POT1 protects telomeres from a transient DNA damage response and determines how human chromosomes end. EMBO J 24, 2667–2678 (2005). 10.1038/sj.emboj.7600733

61 Hockemeyer, D., Daniels, J. P., Takai, H. & de Lange, T. Recent expansion of the telomeric complex in rodents: Two distinct POT1 proteins protect mouse telomeres. Cell 126, 63–77 (2006). 10.1016/j.cell.2006.04.044

62 Lin, C. G. et al. The human telomeric proteome during telomere replication. Nucleic Acids Res 49, 12119–12135 (2021). 10.1093/nar/gkab1015

63 Lancey, C. et al. Structure of the processive human Pol δ holoenzyme. Nat Commun 11, 1109 (2020). 10.1038/s41467-020-14898-6

64 Kelly, T. J. et al. Replication of adenovirus and SV40 chromosomes in vitro. Philos Trans R Soc Lond Biol 317, 429–438 (1987).

65 Tan, W. et al. Preparation and purification of mono-ubiquitinated proteins using Avi-tagged ubiquitin. PLoS One 15, e0229000 (2020). 10.1371/journal.pone.0229000

66 Pinto, C., Kasaciunaite, K., Seidel, R. & Cejka, P. Human DNA2 possesses a cryptic DNA unwinding activity that functionally integrates with BLM or WRN helicases. Elife 5 (2016). 10.7554/eLife.18574

67 Tehseen, M. et al. Proliferating cell nuclear antigen-agarose column: A tag-free and tag-dependent tool for protein purification affinity chromatography. J Chromatogr A 1602, 341–349 (2019). 10.1016/j.chroma.2019.06.008

68 Soman, A. et al. Columnar structure of human telomeric chromatin. Nature 609, 1048–1055 (2022). 10.1038/s41586-022-05236-5

69 Georgescu, R. E. et al. Mechanism of asymmetric polymerase assembly at the eukaryotic replication fork. Nat Struct Mol Biol 21, 664–670 (2014). 10.1038/nsmb.2851

70 Davey, M. J., Fang, L., McInerney, P., Georgescu, R. E. & O’Donnell, M. The DnaC helicase loader is a dual ATP/ADP switch protein. Embo J 21, 3148–3159 (2002).

71 Binz, S. K., Dickson, A. M., Haring, S. J. & Wold, M. S. Functional assays for replication protein A (RPA). Methods Enzymol 409, 11–38 (2006). 10.1016/S0076-6879(05)09002-6

72 Jones, M. L., Baris, Y., Taylor, M. R. G. & Yeeles, J. T. P. Structure of a human replisome shows the organisation and interactions of a DNA replication machine. EMBO J 40, e108819 (2021). 10.15252/embj.2021108819

73 Devbhandari, S., Jiang, J., Kumar, C., Whitehouse, I. & Remus, D. Chromatin Constrains the Initiation and Elongation of DNA Replication. Mol Cell 65, 131–141 (2017). 10.1016/j.molcel.2016.10.035

74 Wilson, R. H. et al. PCNA dependent cellular activities tolerate dramatic perturbations in PCNA client interactions. DNA Repair (Amst) 50, 22–35 (2017). 10.1016/j.dnarep.2016.12.003

75 Raducanu, V. S., Tehseen, M., Shirbini, A., Raducanu, D. V. & Hamdan, S. M. Two chromatographic schemes for protein purification involving the biotin/avidin interaction under native conditions. J Chromatogr A 1621, 461051 (2020). 10.1016/j.chroma.2020.461051

76 Eickhoff, P. et al. Chromosome end protection by RAP1-mediated inhibition of DNA-PK. Nature (2025). 10.1038/s41586-025-08896-1

77 Mazzucco, G., Huda, A., Galli, M., Zanella, E. & Doksani, Y. Purification of mammalian telomeric DNA for single-molecule analysis. Nat Protoc 17, 1444–1467 (2022). 10.1038/s41596-022-00684-9

