## Supplementary figures and images for "Inhibition of lagging strand replication by G-rich telomeric DNA and the shelterin subunit POT1"

### Figure S1

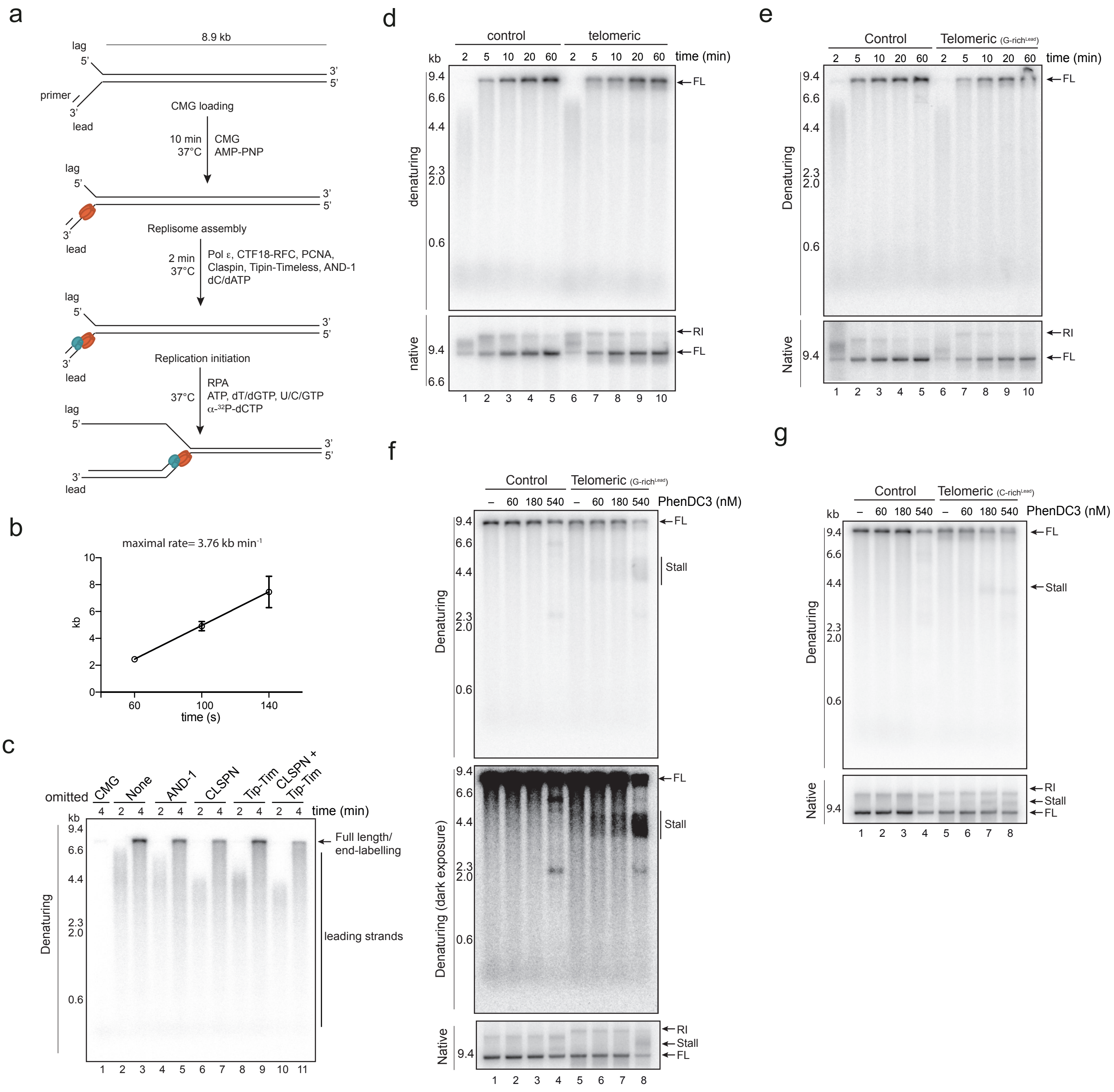

### Figure S2

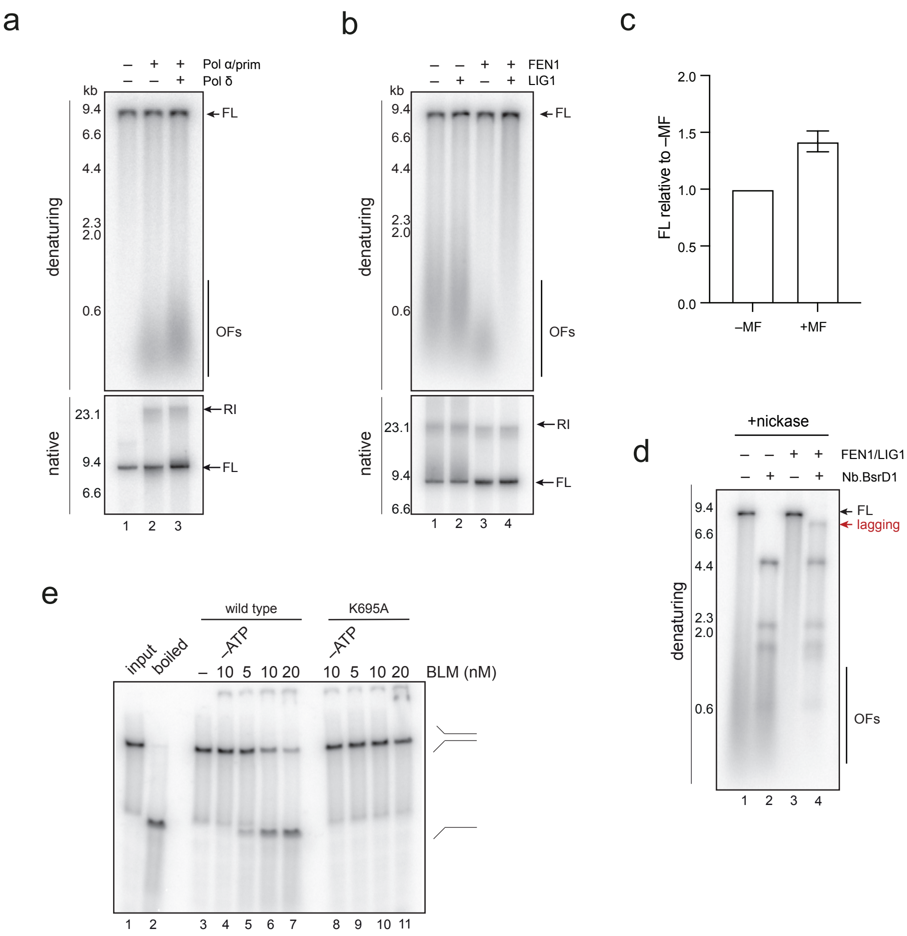

### Figure S3

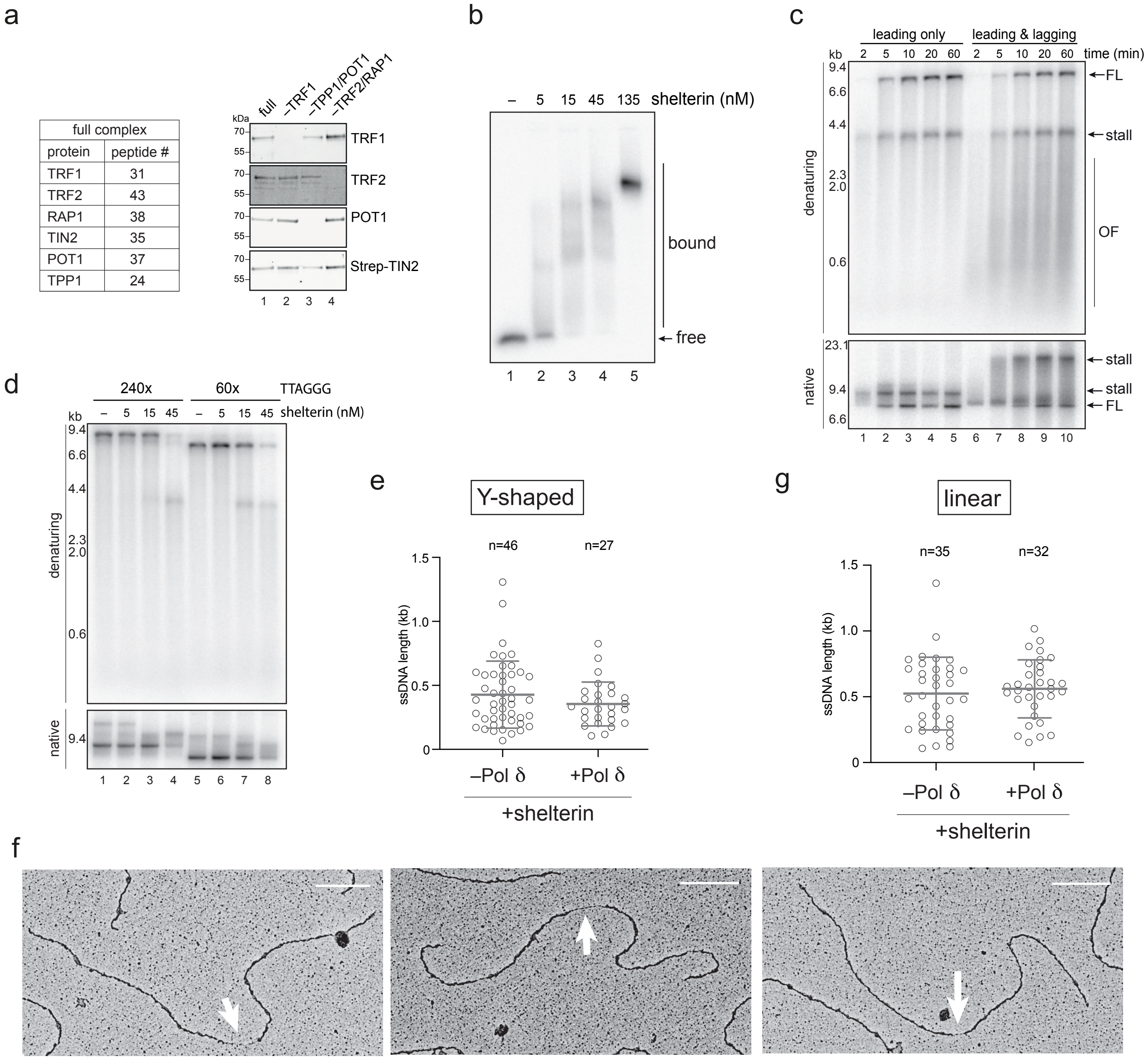
