## Supplementary file detailing the DNA sequences and plasmid constructs used in this study for "Inhibition of lagging strand replication by G-rich telomeric DNA and the shelterin subunit POT1"

**SUPPLEMENTARY INFORMATION**

**Sequence of DNA constructs used for *in vitro* replication:**

**‘Long’ control**

CCTCTGACACATGCAGCTCCCGGACCTCTAGCCCTGCCCAGCGGAGAAGAGGCAACGACCCTCTGACAAGCAGCCCCGGCAGAAGCAGCAGAAGAACCGACGCCCTGACCAGCTCCCCTGGCAGAGATCTGCCCCCATTCGAGGATGAGAGCGAGGGCCTGCTGGGCACAGAGGGACCTCTGGAAGAAGAAGAGGACGGCGAGGAACTGATCGGCGACGGCATGGAACGGGACTACAGAGCCATCCCCGAGCTGGATGCCTATGAGGCCGAAGGCCTGGCCCTGGACGACGAGGATGTGGAAGAACTGACCGCCAGCCAGAGAGAGGCCGCCGAGAGAGCTATGCGGCAGAGAGACAGAGAGGCTGGCAGAGGCCTGGGCAGAATGCGGAGAGGCCTGCTGTACGACAGCGACGAGGAAGATGAGGAAAGACCCGCCAGAAAGCGGAGACAGGTGGAAAGAGCCACCGAGGACGGGGAAGAGGATGAAGAGATGATCGAGAGCATCGAGAACCTGGAAGATCTGAAGGGCCACAGCGTGCGCGAGTGGGTGTCAATGGCTGGCCCCAGACTGGAAATCCACCACCGGTTCAAGAACTTTCTGCGGACCCACGTGGACAGCCACGGCCACAACGTGTTCAAAGAACGGATCAGCGACATGTGCAAAGAGAACCGCGAGAGCCTGGTCGTGAACTACGAGGATCTGGCCGCCAGAGAACACGTGCTGGCCTACTTTCTGCCTGAGGCCCCTGCCGAGCTGCTCCAGATCTTTGATGAGGCCGCCCTGGAAGTGGTGCTGGCCATGTACCCTAAGTACGACCGGATCACCAACCACATCCACGTGCGGATCAGCCATCTGCCCCTGGTGGAAGAACTGCGGAGCCTGAGACAGCTGCACCTGAACCAGCTGATCCGGACCTCCGGCGTCGTGACCTCTTGTACAGGCGTGCTGCCCCAGCTGTCTATGGTCAAGTACAACTGCAACAAGTGCAACTTCGTGCTGGGACCCTTCTGCCAGAGCCAGAACCAGGAAGTGAAGCCCGGCAGCTGTCCCGAGTGTCAGAGCGCCGGACCCTTCGAAGTGAACATGGAAGAGACAATCTACCAGAACTACCAGCGCATCCGCATAATACGACTCACTATAGGCATATGCGGTGTGAAATACCGCACAGATGCGTAAGGAGAAAATACCGCATCAGGCGCCATTCGCCATTCAGGCTGCGCAACTGTTGGGAAGGGCGATCGGTGCGGGCCTCTTCGCTATTACGCCAGCTGGCGAAAGGGGGATGTGCTGCAAGGCGATTAAGTTGGGTAACGCCAGGGTTTTCCCAGTCACGACGTTGTAAAACGACGGCCAGTGAATTCGTCCTGAAGGCTAAGCTGCAAGAGGCCATGAAGCTGCGCCGCTTCGAAGAAAGGCAGAAGCGTCAGGCTCTGTTCAAGCTGGACAACGAGGACGGTTTCGAAGAAGAAGAAGAGGAAGAGGAAGAAATGACCGACGAGTCCGAAGAGGACGGCGAGGAAAAGGTCGAAAAAGAGGAAAAAGAAGAAGAACTCGAGGAAGAAGAGGAAAAAGAAGAGGAAGAAGAAGAAGAGGGCAATCAAGAGACTGCCGAGTTCCTGCTGTCCTCCGAAGAAATCGAGACAAAGGACGAGAAAGAAATGGACAAAGAAAACAACGACGGCTCCTCCGAGATCGGCAAGGCTGTGGGTTTCCTGTCCGTGCCTAAGTCTCTGTCCTCTGACTCTACCCTGCTGCTGTTCAAGGACTCCTCCAGCAAGATGGGCTACTTCCCCACCGAAGAGAAGTCCGAAACCGACGAGAACTCCGGCAAGCAGCCCTCTAAGTTGGACGAGGACGACTCCTGCTCTCTGCTGACCAAAGAGTCCAGCCACAACTCCAGCTTCGAGCTGATCGGTTCTACTATCCCATCCTACCAGCCTTGCAACCGTCAGACCGGTCGTGGTACTTCTTTCTTCCCTACCGCTGGTGGTTTCAGGTCCCCTTCTCCTGGACTGTTCCGTGCTTCCCTGGTTTCCTCCGCTTCCAAGTCCTCCGGAAAGCTGTCCGAGCCTTCTCTGCCTATCGAGGACTCCCAGGACCTGTACAACGCTTCCCCTGAGCCTAAGACACTGTTCCTCGGTGCTGGCGACTTCCAGTTCTGCCTTGAGGACGACACTCAGTCCCAGCTGTTGGACGCTGACGGTTTCCTGAACGTGCGTAACCACCGTAACCAGTACCAGGCTTTGAAGCCTCGTCTGCCCCTGGCTTCCATGGACGAGAATGCTATGGACGCTAACATGGACGAACTGCTGGACCTGTGCACCGGCAAGTTCACTAGCCAGGCTGAGAAGCACTTGCCTCGCAAGTCCGACAAGAAAGAAAACATGGAAGAGTTGCTGAACCTCTGCTCGGGAAAGTTCACCAGTCAGGACGCTTCTACCCCTGCTTCCTCTGAGCTGAACAAGCAAGAGAAAGAATCCTCCATGGGCGACCCTATGGAAGAGGCTTTGGCTCTGTGCTCCGGTAGCTTCCCAACTGACAAAGAGGAAGAGGACGAAGAGGAAGAGTTCGGCGACTTCCGTCTGGTGTCCAACGACAACGAGTTCGACTCTGATGAGGACGAGCACAGCGACTCTGGCAACGACTTGGCTCTGGAAGATCACGAGGATGACGACGAAGAAGAATTGCTGAAGCGTTCCGAGAAGCTCAAGCGTCAGATGCGTCTGCGCAAGTACCTCGAGGATGAGGCTGAGGTGTCCGGTTCTGACGTCGGTTCAGAGGACGAGTACGATGGCGAAGAAATTGACGAATACGAGGAAGATGTGATCGACGAGGTGCTGCCCAATTGGAGGAACTGCAGAGTCATTCTGAGAATAGTGTATGCGGCGACCGAGTTGCTCTTGCCCGGCGTCAATACGGGATAGTACCGCGCCACATAGCAGCACTTTGAAAGTGCTCATCATTGGAAAACGTTCTTCGGGGCGAAAACTCTCAAGGATCTTACCGCTGTTGAGATCCAGTTCGATGTAACCCACTCGTGCACCCAACTGATCTTCAGCATCTTTTACTTTCACCAGCGTTTCTGGGTGAGCAAGAACAGGAAGGCACAATGCCGCAGACAAGGGAATAAGGGCGACACGGAAATGTTGAATACTCATACTCTTCCTGCTTCAATAGTATTGAAGCATCTATCAGGGTTAGTGTCTCATGAGCGGATACATATCTGAATGTATGTAGAACACTAGACACATAGGGGTTCCGCGCACATTTCCCCGAGAAGTGCCACCTGACGTCTAAGACACCAGTATCATCATGACATTGACCTCTAGACgGATCCGGGTCGACCGCCACCATGAGCGGCTTTGACGACCCCGGCATCTTCTACAGCGACAGCTTCGGCGGAGATGCCCAGGCCGATGAAGGCCAGGCCAGAAAGTCTCAGCTCCAGCGGCGGTTCAAAGAGTTCCTGCGGCAGTACAGAGTGGGCACCGACAGAACCGGCTTCACCTTCAAGTACCGGGACGAGCTGAAGCGGCACTACAACCTGGGCGAGTACTGGATCGAGGTGGAAATGGAAGATCTGGCCAGCTTCGACGAGGACCTGGCCGACTACCTGTACAAGCAGCCCGCCGAGCATCTCCAGCTGCTGGAAGAGGCCGCTAAAGAGGTGGCCGACGAAGTGACCAGACCCAGGCCTTCTGGCGAGGAAGTGCTCCAGGACATCCAAGTGATGCTGAAGTCCGACGCCAGCCCCAGCAGCATCAGATCCCTGAAGTCTGACATGATGAGCCACCTCGTGAAGATCCCCGGAATCATCATTGCCGCCTCTGCCGTGCGGGCCAAGGCCACCAGAATCAGCATCCAGTGCAGAAGCTGCCGGAACACCCTGACCAATATCGCCATGCGCCCTGGCCTGGAAGGCTACGCCCTGCCCAGAAAGTGCAACACCGACCAGGCTGGCAGACCCAAGTGCCCCCTGGACCCCTACTTCATCATGCCCGACAAGTGCAAATGCGTGGACTTCCAGACCCTGAAACTCCAGGAACTGCCCGACGCCGTGCCTCACGGCGAAATGCCTAGACACATGCAGCTGTACTGCGACAGATACCTGTGCGACAAGGTGGTGCCCGGCAACAGAGTGACCATCATGGGCATCTACAGCATCAAGAAGTTCGGCCTGACCACCAGCCGGGGCAGAGATAGAGTGGGCGTGGGCATCCGGTCCAGCTACATCAGAGTGCTGGGCATCCAGGTGGACACCGATGGCAGCGGCAGATCCTTTGCTGGGGCCGTGAGCCCCCAGGAAGAGGAAGAGTTTAGACGGCTGGCCGCCCTGCCTAACGTGTACGAAGTGATCAGCAAGTCTATCGCCCCCAGCATCTTCGGCGGCACCGATATGAAGAAGGCCATTGCCTGCCTGCTGTTCGGCGGCTCCAGAAAGAGACTGCCTGACGGCCTGACAAGACGGGGCGACATCAACCTGCTGATGCTGGGCGATCCTGGCACCGCCAAATCACAGCTGCTGAAGTTCGTGGAAAAGTGCAGCCCCATCGGCGTGTACACCAGCGGCAAGGGATCTTCTGCCGCCGGACTGACAGCCAGCGTGATGAGAGATCCCAGCAGCCGGAATTTCATCATGGAAGGCGGCGCTATGGTGCTGGCCGATGGCGGAGTCGTGTGCATCGACGAGTTCGACAAGATGCGCGAGGACGACCGGGTGGCCATTCACGAGGCTATGGAACAGCAGACCATCTCTATCGCCAAGGCCGGAATCACCACCACCCTGAACAGCCGGTGTAGCGTGCTGGCTGCCGCCAATAGCGTGTTCGGCAGATGGGACGAGACAAAGGGCGAGGACAACATCGACTTCATGCCCACCATCCTGAGCCGCTTCGACATGATCTTCATCGTGAAGGACGAGCACAACGAGGAACGGGACGTGATGCTGGCCAAGCACGTGATCACCCTGCACGTGTCCGCCCTGACACAGACACAGGCCGTGGAAGGCGAGATCGACCTGGCCAAACTGAAGAAGTTTATCGCCTACTGTCGCGTGAAGTGCGGCCCCAGACTGTCTGCCGAGGCCGCCGAGAAGCTGAAGAACCGGTACATCATCATGCGGAGCGGAGCCCGGCAGCACGAGAGAGACAGCGATCGGAGAAGCAGCATCCCCATCACCGTGCGGCAGCTGGAAGCCATCGTGCGGATTGCCGAAGCCCTGAGCAAGATGAAGCTCCAGCCCTTCGCCACCGAGGCCGACGTGGAAGAAGCTCTGAGACTGTTCCAGGTGTCAACCCTGGACGCCGCCCTGAGCGGAACACTGTCTGGCGTGGAAGGATTCACCAGCCAGGAAGATCAGGAAATGCTGAGCAGAATCGAGAAACAGCTGAAGAGAAGATTCGCCATCGGAAGCCAGGTGTCCGAGCACAGCATCATCAAGGACTTCACCAAGCAGAAGTACCCCGAGCACGCCATCCACAAGGTGCTCCAGCTGATGCTGAGAAGAGGCGAGATCCAGCACCGGATGCAGCGGAAGGTGCTGTACCGGCTGAAATGAGGCGCCGGAATCTTTGCAGGTGCTTACGCTTACTACCTAAACTACAATGGTGTTGTCGCTACTAGTGCCGCTTCTTCATCCACTGCATCTGGTGCTTCCGCTTCCGTTACCGGTTCTAAGAAACTTCAGAAACTTCAGAAACCAGTGCTGCCCCTAAGACATACACTACTGCCACTGTTACTCAATGTGATGACAATGGTTGTAACGTCAAGATAATCACCTCTCAAATACCTGAAGCTACTTCAACCGTCACCGCAACTAGTGCTTCTCCAAAGTCATACACTACTGTCACTTCTGAGGGTTCTAAAGCAACCTCATTGAGTTCTCACTGCCCGCTTTCCAGTCGGGAAACCTGTCGTGCCAGCTGCATTAATGAATCGGCCAACGCGCGGGGAGAGGCGGTTTGCGTATTGGGCGCTCTTCCGCTTCCTCGCTCACTGACTCGCTGCGCTCGGTCGTTCGGCTGCGGCGAGCGGTATCAGCTCACTCAAAGGCGGTAATACGGTTATCCACAGAATCAGGGGATAACGCAGGACAGAACATGTGAGCAGAAGGCCAGCACGAGGCCAGGAGCCGTAAGAAGGCCGCGTTGCTGGCGTTTTTCCATAGGCTCCGCCCCCTCGCGAGCTGAGGACTTAAGTGCTGAGGTCGCGAGCATTCAGAAGAGCCATTCAAGGGATCGATGTGGTCGCACGACTACGCATCCCTCTGGGCCTTACATAGCCGGATACAGTGACTTATCGATACGCGTGTTAACCTCGAGGCGGCCGCCATGGTGACAGGACTTGCATAGCTGCGTGCGGGGGAAGGAACTCTTGCGTGTGAGTATGTAGACCCCTGTACTACGGATGCGGGCAGAAGATGTGGGCAGAGACACCCGCGTCAAGTTCTCGACCTTCCCGTGGGAGGTGTTCCAGTCCGCCATACGACCATACCGTTCGGGCATGGCACTATGTACGCGATATCGCTAGCGTTTGCGGGGCACAGCAATATGCAGGCATGCAAGCTTGGCGTAATCATGGTCATAGCTGTTTCCTGTGTGAAATTGTTATCCGCTCACAATTCCACACAACATACGAGCCGGAAGCATAAAGTGTAAAGCCTGGGGTGCCTAATGAGTGAGCTAACTCACATTAATTGCGTTGCGCTCACTGCCCGCTTTCCAGTCGGGAAACCTGTCGTGCCAGCTGCATTAATGAATCGGCCAACGCGCGGGGAGAGGCGGTTTGCGTATTGGGCGCTCTTCCGCTTCCTCGCTCACTGACTCGCTGCGCTCGGTCGTTCGGCTGCGGCGAGCGGTATCAGCTCACTCAAAGGCGGTAATACGGTTATCCACAGAATCAGGGGATAACGCAGGAAAGAACATGTGAGCAAAAGGCCAGCAAAAGGCCAGGAACCGTAAAAAGGCCGCGTTGCTGGCGTTTTTCCATAGGCTCCGCCCCCCTGACGAGCATCACAAAAATCGACGCTCAAGTCAGAGGTGGCGAAACCCGACAGGACTATAAAGATACCAGGCGTTTCCCCCTGGAAGCTCCCTCGTGCGCTCTCCTGTTCCGACCCTGCCGCTTACCGGATACCTGTCCGCCTTTCTCCCTTCGGGAAGCGTGGCGCTTTCTCATAGCTCACGCTGTAGGTATCTCAGTTCGGTGTAGGTCGTTCGCTCCAAGCTGGGCTGTGTGCACGAACCCCCCGTTCAGCCCGACCGCTGCGCCTTATCCGGTAACTATCGTCTTGAGTCCAACCCGGTAAGACACGACTTATCGCCACTGGCAGCAGCCACTGGTAACAGGATTAGCAGAGCGAGGTATGTAGGCGGTGCTACAGAGTTCTTGAAGTGGTGGCCTAACTACGGCTACACTAGAAGAACAGTATTTGGTATCTGCGCTCTGCTGAAGCCAGTTACCTTCGGAAAAAGAGTTGGTAGCTCTTGATCCGGCAAACAAACCACCGCTGGTAGCGGTGGTTTTTTTGTTTGCAAGCAGCAGATTACGCGCAGAAAAAAAGGATCTCAAGAAGATCCTTTGATCTTTTCTACGGGGTCTGACGCTCAGTGGAACGAAAACTCACGTTAAGGGATTTTGGTCATGAGATTATCAAAAAGGATCTTCACCTAGATCCTTTTAAATTAAAAATGAAGTTTTAAATCAATCTAAAGTATATATGAGTAAACTTGGTCTGACAGTTACCAATGCTTAATCAGTGAGGCACCTATCTCAGCGATCTGTCTATTTCGTTCATCCATAGTTGCCTGACTCCCCGTCGTGTAGATAACTACGATACGGGAGGGCTTACCATCTGGCCCCAGTGCTGCAATGATACCGCGAGACCCACGCTCACCGGCTCCAGATTTATCAGCAATAAACCAGCCAGCCGGAAGGGCCGAGCGCAGAAGTGGTCCTGCAACTTTATCCGCCTCCATCCAGTCTATTAATTGTTGCCGGGAAGCTAGAGTAAGTAGTTCGCCAGTTAATAGTTTGCGCAACGTTGTTGCCATTGCTACAGGCATCGTGGTGTCACGCTCGTCGTTTGGTATGGCTTCATTCAGCTCCGGTTCCCAACGATCAAGGCGAGTTACATGATCCCCCATGTTGTGCAAAAAAGCGGTTAGCTCCTTCGGTCCTCCGATCGTTGTCAGAAGTAAGTTGGCCGCAGTGTTATCACTCATGGTTATGGCAGCACTGCATAATTCTCTTACTGTCATGCCATCCGTAAGATGCTTTTCTGTGACTGGTGAGTACTCAACCAAGTCATTCTGAGAATAGTGTATGCGGCGACCGAGTTGCTCTTGCCCGGCGTCAATACGGGATAATACCGCGCCACATAGCAGAACTTTAAAAGTGCTCATCATTGGAAAACGTTCTTCGGGGCGAAAACTCTCAAGGATCTTACCGCTGTTGAGATCCAGTTCGATGTAACCCACTCGTGCACCCAACTGATCTTCAGCATCTTTTACTTTCACCAGCGTTTCTGGGTGAGCAAAAACAGGAAGGCAAAATGCCGCAAAAAAGGGAATAAGGGCGACACGGAAATGTTGAATACTCATACTCTTCCTTTTTCAATATTATTGAAGCATTTATCAGGGTTATTGTCTCATGAGCGGATACATATTTGAATGTATTTAGAAAAATAAACAAATAGGGGTTCCGCGCACATTTCCCCGAAAAGTGCCACCTGACGTCTAAGAAACCATTATTATCATGACATTAACCTATAAAAATAGGCGTATCACGAGGCCCTTTCG

**‘Long’ telomeric**

CCTCTGACACATGCAGCTCCCGGACCTCTAGCCCTGCCCAGCGGAGAAGAGGCAACGACCCTCTGACAAGCAGCCCCGGCAGAAGCAGCAGAAGAACCGACGCCCTGACCAGCTCCCCTGGCAGAGATCTGCCCCCATTCGAGGATGAGAGCGAGGGCCTGCTGGGCACAGAGGGACCTCTGGAAGAAGAAGAGGACGGCGAGGAACTGATCGGCGACGGCATGGAACGGGACTACAGAGCCATCCCCGAGCTGGATGCCTATGAGGCCGAAGGCCTGGCCCTGGACGACGAGGATGTGGAAGAACTGACCGCCAGCCAGAGAGAGGCCGCCGAGAGAGCTATGCGGCAGAGAGACAGAGAGGCTGGCAGAGGCCTGGGCAGAATGCGGAGAGGCCTGCTGTACGACAGCGACGAGGAAGATGAGGAAAGACCCGCCAGAAAGCGGAGACAGGTGGAAAGAGCCACCGAGGACGGGGAAGAGGATGAAGAGATGATCGAGAGCATCGAGAACCTGGAAGATCTGAAGGGCCACAGCGTGCGCGAGTGGGTGTCAATGGCTGGCCCCAGACTGGAAATCCACCACCGGTTCAAGAACTTTCTGCGGACCCACGTGGACAGCCACGGCCACAACGTGTTCAAAGAACGGATCAGCGACATGTGCAAAGAGAACCGCGAGAGCCTGGTCGTGAACTACGAGGATCTGGCCGCCAGAGAACACGTGCTGGCCTACTTTCTGCCTGAGGCCCCTGCCGAGCTGCTCCAGATCTTTGATGAGGCCGCCCTGGAAGTGGTGCTGGCCATGTACCCTAAGTACGACCGGATCACCAACCACATCCACGTGCGGATCAGCCATCTGCCCCTGGTGGAAGAACTGCGGAGCCTGAGACAGCTGCACCTGAACCAGCTGATCCGGACCTCCGGCGTCGTGACCTCTTGTACAGGCGTGCTGCCCCAGCTGTCTATGGTCAAGTACAACTGCAACAAGTGCAACTTCGTGCTGGGACCCTTCTGCCAGAGCCAGAACCAGGAAGTGAAGCCCGGCAGCTGTCCCGAGTGTCAGAGCGCCGGACCCTTCGAAGTGAACATGGAAGAGACAATCTACCAGAACTACCAGCGCATCCGCATAATACGACTCACTATAGGCATATGCGGTGTGAAATACCGCACAGATGCGTAAGGAGAAAATACCGCATCAGGCGCCATTCGCCATTCAGGCTGCGCAACTGTTGGGAAGGGCGATCGGTGCGGGCCTCTTCGCTATTACGCCAGCTGGCGAAAGGGGGATGTGCTGCAAGGCGATTAAGTTGGGTAACGCCAGGGTTTTCCCAGTCACGACGTTGTAAAACGACGGCCAGTGAATTCGAGCTCGGTACCCGGGGATCCTCTAGAGTCGACGAAGACTTAGGGTTAGGGTTAGGGTTAGGGTTAGGGTTAGGGTTAGGGTTAGGGTTAGGGTTAGGGTTAGGGTTAGGGTTAGGGTTAGGGTTAGGGTTAGGGTTAGGGTTAGGGTTAGGGTTAGGGTTAGGGTTAGGGTTAGGGTTAGGGTTAGGGTTAGGGTTAGGGTTAGGGTTAGGGTTAGGGTTAGGGTTAGGGTTAGGGTTAGGGTTAGGGTTAGGGTTAGGGTTAGGGTTAGGGTTAGGGTTAGGGTTAGGGTTAGGGTTAGGGTTAGGGTTAGGGTTAGGGTTAGGGTTAGGGTTAGGGTTAGGGTTAGGGTTAGGGTTAGGGTTAGGGTTAGGGTTAGGGTTAGGGTTAGGGTTAGGGTTAGGGTTAGGGTTAGGGTTAGGGTTAGGGTTAGGGTTAGGGTTAGGGTTAGGGTTAGGGTTAGGGTTAGGGTTAGGGTTAGGGTTAGGGTTAGGGTTAGGGTTAGGGTTAGGGTTAGGGTTAGGGTTAGGGTTAGGGTTAGGGTTAGGGTTAGGGTTAGGGTTAGGGTTAGGGTTAGGGTTAGGGTTAGGGTTAGGGTTAGGGTTAGGGTTAGGGTTAGGGTTAGGGTTAGGGTTAGGGTTAGGGTTAGGGTTAGGGTTAGGGTTAGGGTTAGGGTTAGGGTTAGGGTTAGGGTTAGGGTTAGGGTTAGGGTTAGGGTTAGGGTTAGGGTTAGGGTTAGGGTTAGGGTTAGGGTTAGGGTTAGGGTTAGGGTTAGGGTTAGGGTTAGGGTTAGGGTTAGGGTTAGGGTTAGGGTTAGGGTTAGGGTTAGGGTTAGGGTTAGGGTTAGGGTTAGGGTTAGGGTTAGGGTTAGGGTTAGGGTTAGGGTTAGGGTTAGGGTTAGGGTTAGGGTTAGGGTTAGGGTTAGGGTTAGGGTTAGGGTTAGGGTTAGGGTTAGGGTTAGGGTTAGGGTTAGGGTTAGGGTTAGGGTTAGGGTTAGGGTTAGGGTTAGGGTTAGGGTTAGGGTTAGGGTTAGGGTTAGGGTTAGGGTTAGGGTTAGGGTTAGGGTTAGGGTTAGGGTTAGGGTTAGGGTTAGGGTTAGGGTTAGGGTTAGGGTTAGGGTTAGGGTTAGGGTTAGGGTTAGGGTTAGGGTTAGGGTTAGGGTTAGGGTTAGGGTTAGGGTTAGGGTTAGGGTTAGGGTTAGGGTTAGGGTTAGGGTTAGGGTTAGGGTTAGGGTTAGGGTTAGGGTTAGGGTTAGGGTTAGGGTTAGGGTTAGGGTTAGGGTTAGGGTTAGGGTTAGGGTTAGGGTTAGGGTTAGGGTTAGGGTTAGGGTTAGGGTTAGGGTTAGGGTTAGGGTTAGGGTTAGGGTTAGGGTTAGGGTTAGGGTTAGGGTTAGGGTTAGGGTTAGGGTTAGGGTTAGGGTTAGGGTTAGGGTTAGGGTTAGGGTTAGGGTTAGGGTTAGGGTTAGGGTTAGGGTTAGGGCGAGACGTGTCCCGTCGACCTGCATAGCGCTCAATTGCAACCACTGCAGAGTCATTCTGAGAATAGTGTATGCGGCGACCGAGTTGCTCTTGCCCGGCGTCAATACGGGATAGTACCGCGCCACATAGCAGCACTTTGAAAGTGCTCATCATTGGAAAACGTTCTTCGGGGCGAAAACTCTCAAGGATCTTACCGCTGTTGAGATCCAGTTCGATGTAACCCACTCGTGCACCCAACTGATCTTCAGCATCTTTTACTTTCACCAGCGTTTCTGGGTGAGCAAGAACAGGAAGGCACAATGCCGCAGACAAGGGAATAAGGGCGACACGGAAATGTTGAATACTCATACTCTTCCTGCTTCAATAGTATTGAAGCATCTATCAGGGTTAGTGTCTCATGAGCGGATACATATCTGAATGTATGTAGAACACTAGACACATAGGGGTTCCGCGCACATTTCCCCGAGAAGTGCCACCTGACGTCTAAGACACCAGTATCATCATGACATTGACCTCTAGACgGATCCGGGTCGACCGCCACCATGAGCGGCTTTGACGACCCCGGCATCTTCTACAGCGACAGCTTCGGCGGAGATGCCCAGGCCGATGAAGGCCAGGCCAGAAAGTCTCAGCTCCAGCGGCGGTTCAAAGAGTTCCTGCGGCAGTACAGAGTGGGCACCGACAGAACCGGCTTCACCTTCAAGTACCGGGACGAGCTGAAGCGGCACTACAACCTGGGCGAGTACTGGATCGAGGTGGAAATGGAAGATCTGGCCAGCTTCGACGAGGACCTGGCCGACTACCTGTACAAGCAGCCCGCCGAGCATCTCCAGCTGCTGGAAGAGGCCGCTAAAGAGGTGGCCGACGAAGTGACCAGACCCAGGCCTTCTGGCGAGGAAGTGCTCCAGGACATCCAAGTGATGCTGAAGTCCGACGCCAGCCCCAGCAGCATCAGATCCCTGAAGTCTGACATGATGAGCCACCTCGTGAAGATCCCCGGAATCATCATTGCCGCCTCTGCCGTGCGGGCCAAGGCCACCAGAATCAGCATCCAGTGCAGAAGCTGCCGGAACACCCTGACCAATATCGCCATGCGCCCTGGCCTGGAAGGCTACGCCCTGCCCAGAAAGTGCAACACCGACCAGGCTGGCAGACCCAAGTGCCCCCTGGACCCCTACTTCATCATGCCCGACAAGTGCAAATGCGTGGACTTCCAGACCCTGAAACTCCAGGAACTGCCCGACGCCGTGCCTCACGGCGAAATGCCTAGACACATGCAGCTGTACTGCGACAGATACCTGTGCGACAAGGTGGTGCCCGGCAACAGAGTGACCATCATGGGCATCTACAGCATCAAGAAGTTCGGCCTGACCACCAGCCGGGGCAGAGATAGAGTGGGCGTGGGCATCCGGTCCAGCTACATCAGAGTGCTGGGCATCCAGGTGGACACCGATGGCAGCGGCAGATCCTTTGCTGGGGCCGTGAGCCCCCAGGAAGAGGAAGAGTTTAGACGGCTGGCCGCCCTGCCTAACGTGTACGAAGTGATCAGCAAGTCTATCGCCCCCAGCATCTTCGGCGGCACCGATATGAAGAAGGCCATTGCCTGCCTGCTGTTCGGCGGCTCCAGAAAGAGACTGCCTGACGGCCTGACAAGACGGGGCGACATCAACCTGCTGATGCTGGGCGATCCTGGCACCGCCAAATCACAGCTGCTGAAGTTCGTGGAAAAGTGCAGCCCCATCGGCGTGTACACCAGCGGCAAGGGATCTTCTGCCGCCGGACTGACAGCCAGCGTGATGAGAGATCCCAGCAGCCGGAATTTCATCATGGAAGGCGGCGCTATGGTGCTGGCCGATGGCGGAGTCGTGTGCATCGACGAGTTCGACAAGATGCGCGAGGACGACCGGGTGGCCATTCACGAGGCTATGGAACAGCAGACCATCTCTATCGCCAAGGCCGGAATCACCACCACCCTGAACAGCCGGTGTAGCGTGCTGGCTGCCGCCAATAGCGTGTTCGGCAGATGGGACGAGACAAAGGGCGAGGACAACATCGACTTCATGCCCACCATCCTGAGCCGCTTCGACATGATCTTCATCGTGAAGGACGAGCACAACGAGGAACGGGACGTGATGCTGGCCAAGCACGTGATCACCCTGCACGTGTCCGCCCTGACACAGACACAGGCCGTGGAAGGCGAGATCGACCTGGCCAAACTGAAGAAGTTTATCGCCTACTGTCGCGTGAAGTGCGGCCCCAGACTGTCTGCCGAGGCCGCCGAGAAGCTGAAGAACCGGTACATCATCATGCGGAGCGGAGCCCGGCAGCACGAGAGAGACAGCGATCGGAGAAGCAGCATCCCCATCACCGTGCGGCAGCTGGAAGCCATCGTGCGGATTGCCGAAGCCCTGAGCAAGATGAAGCTCCAGCCCTTCGCCACCGAGGCCGACGTGGAAGAAGCTCTGAGACTGTTCCAGGTGTCAACCCTGGACGCCGCCCTGAGCGGAACACTGTCTGGCGTGGAAGGATTCACCAGCCAGGAAGATCAGGAAATGCTGAGCAGAATCGAGAAACAGCTGAAGAGAAGATTCGCCATCGGAAGCCAGGTGTCCGAGCACAGCATCATCAAGGACTTCACCAAGCAGAAGTACCCCGAGCACGCCATCCACAAGGTGCTCCAGCTGATGCTGAGAAGAGGCGAGATCCAGCACCGGATGCAGCGGAAGGTGCTGTACCGGCTGAAATGAGGCGCCGGAATCTTTGCAGGTGCTTACGCTTACTACCTAAACTACAATGGTGTTGTCGCTACTAGTGCCGCTTCTTCATCCACTGCATCTGGTGCTTCCGCTTCCGTTACCGGTTCTAAGAAACTTCAGAAACTTCAGAAACCAGTGCTGCCCCTAAGACATACACTACTGCCACTGTTACTCAATGTGATGACAATGGTTGTAACGTCAAGATAATCACCTCTCAAATACCTGAAGCTACTTCAACCGTCACCGCAACTAGTGCTTCTCCAAAGTCATACACTACTGTCACTTCTGAGGGTTCTAAAGCAACCTCATTGAGTTCTCACTGCCCGCTTTCCAGTCGGGAAACCTGTCGTGCCAGCTGCATTAATGAATCGGCCAACGCGCGGGGAGAGGCGGTTTGCGTATTGGGCGCTCTTCCGCTTCCTCGCTCACTGACTCGCTGCGCTCGGTCGTTCGGCTGCGGCGAGCGGTATCAGCTCACTCAAAGGCGGTAATACGGTTATCCACAGAATCAGGGGATAACGCAGGACAGAACATGTGAGCAGAAGGCCAGCACGAGGCCAGGAGCCGTAAGAAGGCCGCGTTGCTGGCGTTTTTCCATAGGCTCCGCCCCCTCGCGAGCTGAGGACTTAAGTGCTGAGGTCGCGAGCATTCAGAAGAGCCATTCAAGGGATCGATGTGGTCGCACGACTACGCATCCCTCTGGGCCTTACATAGCCGGATACAGTGACTTATCGATACGCGTGTTAACCTCGAGGCGGCCGCCATGGTGACAGGACTTGCATAGCTGCGTGCGGGGGAAGGAACTCTTGCGTGTGAGTATGTAGACCCCTGTACTACGGATGCGGGCAGAAGATGTGGGCAGAGACACCCGCGTCAAGTTCTCGACCTTCCCGTGGGAGGTGTTCCAGTCCGCCATACGACCATACCGTTCGGGCATGGCACTATGTACGCGATATCGCTAGCGTTTGCGGGGCACAGCAATATGCAGGCATGCAAGCTTGGCGTAATCATGGTCATAGCTGTTTCCTGTGTGAAATTGTTATCCGCTCACAATTCCACACAACATACGAGCCGGAAGCATAAAGTGTAAAGCCTGGGGTGCCTAATGAGTGAGCTAACTCACATTAATTGCGTTGCGCTCACTGCCCGCTTTCCAGTCGGGAAACCTGTCGTGCCAGCTGCATTAATGAATCGGCCAACGCGCGGGGAGAGGCGGTTTGCGTATTGGGCGCTCTTCCGCTTCCTCGCTCACTGACTCGCTGCGCTCGGTCGTTCGGCTGCGGCGAGCGGTATCAGCTCACTCAAAGGCGGTAATACGGTTATCCACAGAATCAGGGGATAACGCAGGAAAGAACATGTGAGCAAAAGGCCAGCAAAAGGCCAGGAACCGTAAAAAGGCCGCGTTGCTGGCGTTTTTCCATAGGCTCCGCCCCCCTGACGAGCATCACAAAAATCGACGCTCAAGTCAGAGGTGGCGAAACCCGACAGGACTATAAAGATACCAGGCGTTTCCCCCTGGAAGCTCCCTCGTGCGCTCTCCTGTTCCGACCCTGCCGCTTACCGGATACCTGTCCGCCTTTCTCCCTTCGGGAAGCGTGGCGCTTTCTCATAGCTCACGCTGTAGGTATCTCAGTTCGGTGTAGGTCGTTCGCTCCAAGCTGGGCTGTGTGCACGAACCCCCCGTTCAGCCCGACCGCTGCGCCTTATCCGGTAACTATCGTCTTGAGTCCAACCCGGTAAGACACGACTTATCGCCACTGGCAGCAGCCACTGGTAACAGGATTAGCAGAGCGAGGTATGTAGGCGGTGCTACAGAGTTCTTGAAGTGGTGGCCTAACTACGGCTACACTAGAAGAACAGTATTTGGTATCTGCGCTCTGCTGAAGCCAGTTACCTTCGGAAAAAGAGTTGGTAGCTCTTGATCCGGCAAACAAACCACCGCTGGTAGCGGTGGTTTTTTTGTTTGCAAGCAGCAGATTACGCGCAGAAAAAAAGGATCTCAAGAAGATCCTTTGATCTTTTCTACGGGGTCTGACGCTCAGTGGAACGAAAACTCACGTTAAGGGATTTTGGTCATGAGATTATCAAAAAGGATCTTCACCTAGATCCTTTTAAATTAAAAATGAAGTTTTAAATCAATCTAAAGTATATATGAGTAAACTTGGTCTGACAGTTACCAATGCTTAATCAGTGAGGCACCTATCTCAGCGATCTGTCTATTTCGTTCATCCATAGTTGCCTGACTCCCCGTCGTGTAGATAACTACGATACGGGAGGGCTTACCATCTGGCCCCAGTGCTGCAATGATACCGCGAGACCCACGCTCACCGGCTCCAGATTTATCAGCAATAAACCAGCCAGCCGGAAGGGCCGAGCGCAGAAGTGGTCCTGCAACTTTATCCGCCTCCATCCAGTCTATTAATTGTTGCCGGGAAGCTAGAGTAAGTAGTTCGCCAGTTAATAGTTTGCGCAACGTTGTTGCCATTGCTACAGGCATCGTGGTGTCACGCTCGTCGTTTGGTATGGCTTCATTCAGCTCCGGTTCCCAACGATCAAGGCGAGTTACATGATCCCCCATGTTGTGCAAAAAAGCGGTTAGCTCCTTCGGTCCTCCGATCGTTGTCAGAAGTAAGTTGGCCGCAGTGTTATCACTCATGGTTATGGCAGCACTGCATAATTCTCTTACTGTCATGCCATCCGTAAGATGCTTTTCTGTGACTGGTGAGTACTCAACCAAGTCATTCTGAGAATAGTGTATGCGGCGACCGAGTTGCTCTTGCCCGGCGTCAATACGGGATAATACCGCGCCACATAGCAGAACTTTAAAAGTGCTCATCATTGGAAAACGTTCTTCGGGGCGAAAACTCTCAAGGATCTTACCGCTGTTGAGATCCAGTTCGATGTAACCCACTCGTGCACCCAACTGATCTTCAGCATCTTTTACTTTCACCAGCGTTTCTGGGTGAGCAAAAACAGGAAGGCAAAATGCCGCAAAAAAGGGAATAAGGGCGACACGGAAATGTTGAATACTCATACTCTTCCTTTTTCAATATTATTGAAGCATTTATCAGGGTTATTGTCTCATGAGCGGATACATATTTGAATGTATTTAGAAAAATAAACAAATAGGGGTTCCGCGCACATTTCCCCGAAAAGTGCCACCTGACGTCTAAGAAACCATTATTATCATGACATTAACCTATAAAAATAGGCGTATCACGAGGCCCTTTCG

**‘Short’ control**

GTTTGCAAGCAGCAGATTACGCGCAGAAAAAAAGGATCTCAAGAAGATCCTTTGATCTTTTCTACGGGGTCTGACGCTCAGTGGAACGAAAACTCACGTTAAGGGATTTTGGTCATGAGATTATCAAAAAGGATCTTCACCTAGATCCTTTTAAATTAAAAATGAAGTTTTAAATCAATCTAAAGTATATATGAGTAAACTTGGTCTGACAGTTACCAATGCTTAATCAGTGAGGCACCTATCTCAGCGATCTGTCTATTTCGTTCATCCATAGTTGCCTGACTCCCCGTCGTGTAGATAACTACGATACGGGAGGGCTTACCATCTGGCCCCAGTGCTGCAATGATACCGCGAGACCCACGCTCACCGGCTCCAGATTTATCAGCAATAAACCAGCCAGCCGGAAGGGCCGAGCGCAGAAGTGGTCCTGCAACTTTATCCGCCTCCATCCAGTCTATTAATTGTTGCCGGGAAGCTAGAGTAAGTAGTTCGCCAGTTAATAGTTTGCGCAACGTTGTTGCCATTGCTACAGGCATCGTGGTGTCACGCTCGTCGTTTGGTATGGCTTCATTCAGCTCCGGTTCCCAACGATCAAGGCGAGTTACATGATCCCCCATGTTGTGCAAAAAAGCGGTTAGCTCCTTCGGTCCTCCGATCGTTGTCAGAAGTAAGTTGGCCGCAGTGTTATCACTCATGGTTATGGCAGCACTGCATAATTCTCTTACTGTCATGCCATCCGTAAGATGCTTTTCTGTGACTGGTGAGTACTCAACCAAGTCATTCTGAGAATAGTGTATGCGGCGACCGAGTTGCTCTTGCCCGGCGTCAATACGGGATAATACCGCGCCACATAGCAGAACTTTAAAAGTGCTCATCATTGGAAAACGTTCTTCGGGGCGAAAACTCTCAAGGATCTTACCGCTGTTGAGATCCAGTTCGATGTAACCCACTCGTGCACCCAACTGATCTTCAGCATCTTTTACTTTCACCAGCGTTTCTGGGTGAGCAAAAACAGGAAGGCAAAATGCCGCAAAAAAGGGAATAAGGGCGACACGGAAATGTTGAATACTCATACTCTTCCTTTTTCAATATTATTGAAGCATTTATCAGGGTTATTGTCTCATGAGCGGATACATATTTGAATGTATTTAGAAAAATAAACAAATAGGGGTTCCGCGCACATTTCCCCGAAAAGTGCCACCTGACGTCTAAGAAACCATTATTATCATGACATTAACCTATAAAAATAGGCGTATCACGAGGCCCTTTCCCTCTGACACATGCAGCTCCCGGACCTCTAGCCCTGCCCAGCGGAGAAGAGGCAACGACCCTCTGACAAGCAGCCCCGGCAGAAGCAGCAGAAGAACCGACGCCCTGACCAGCTCCCCTGGCAGAGATCTGCCCCCATTCGAGGATGAGAGCGAGGGCCTGCTGGGCACAGAGGGACCTCTGGAAGAAGAAGAGGACGGCGAGGAACTGATCGGCGACGGCATGGAACGGGACTACAGAGCCATCCCCGAGCTGGATGCCTATGAGGCCGAAGGCCTGGCCCTGGACGACGAGGATGTGGAAGAACTGACCGCCAGCCAGAGAGAGGCCGCCGAGAGAGCTATGCGGCAGAGAGACAGAGAGGCTGGCAGAGGCCTGGGCAGAATGCGGAGAGGCCTGCTGTACGACAGCGACGAGGAAGATGAGGAAAGACCCGCCAGAAAGCGGAGACAGGTGGAAAGAGCCACCGAGGACGGGGAAGAGGATGAAGAGATGATCGAGAGCATCGAGAACCTGGAAGATCTGAAGGGCCACAGCGTGCGCGAGTGGGTGTCAATGGCTGGCCCCAGACTGGAAATCCACCACCGGTTCAAGAACTTTCTGCGGACCCACGTGGACAGCCACGGCCACAACGTGTTCAAAGAACGGATCAGCGACATGTGCAAAGAGAACCGCGAGAGCCTGGTCGTGAACTACGAGGATCTGGCCGCCAGAGAACACGTGCTGGCCTACTTTCTGCCTGAGGCCCCTGCCGAGCTGCTCCAGATCTTTGATGAGGCCGCCCTGGAAGTGGTGCTGGCCATGTACCCTAAGTACGACCGGATCACCAACCACATCCACGTGCGGATCAGCCATCTGCCCCTGGTGGAAGAACTGCGGAGCCTGAGACAGCTGCACCTGAACCAGCTGATCCGGACCTCCGGCGTCGTGACCTCTTGTACAGGCGTGCTGCCCCAGCTGTCTATGGTCAAGTACAACTGCAACAAGTGCAACTTCGTGCTGGGACCCTTCTGCCAGAGCCAGAACCAGGAAGTGAAGCCCGGCAGCTGTCCCGAGTGTCAGAGCGCCGGACCCTTCGAAGTGAACATGGAAGAGACAATCTACCAGAACTACCAGCGCATCCGCATAATACGACTCACTATAGGCATATGCGGTGTGAAATACCGCACAGATGCGTAAGGAGAAAATACCGCATCAGGCGCCATTCGCCATTCAGGCTGCGCAACTGTTGGGAAGGGCGATCGGTGCGGGCCTCTTCGCTATTACGCCAGCTGGCGAAAGGGGGATGTGCTGCAAGGCGATTAAGTTGGGTAACGCCAGGGTTTTCCCAGTCACGACGTTGTAAAACGACGGCCAGTGAATTCGTCCTGAAGGCTAAGCTGCAAGAGGCCATGAAGCTGCGCCGCTTCGAAGAAAGGCAGAAGCGTCAGGCTCTGTTCAAGCTGGACAACGAGGACGGTTTCGAAGAAGAAGAAGAGGAAGAGGAAGAAATGACCGACGAGTCCGAAGAGGACGGCGAGGTACTTGCGCAGACGCATCTGACGCTTGAGCTTCTCGGAACGCTTCAGCAATTCTTCTTCGTCGTCATCCTCGTGATCTTCCAGAGCCAAGTCGTTGCCAGAGTCGCTGTGCTCGTCCTCATCAGAGTCGAACTCGTTGTCGTTGGACACCAGACGGAAGTCGGAGGACAGCAGGAACTCGGCAGTCTCTTGATTGCGATGAGGCTGAGGTGTCCGGTTCTGACGTCGGTTCAGAGGACGAGTACGATGGCGAAGAAATTGACGAATACGAGGAAGATGTGATCGACGAGGTGCTGCCCAATTGGAGGAACGAGTCATTCTGAGAATAGTGTATGCGGCGACCGAGTTGCTCTTGCCCGGCGTCAATACGGGATAGTACCGCGCCACATAGCAGCACTTTGAAAGTGCTCATCATTGGAAAACGTTCTTCGGGGCGAAAACTCTCAAGGATCTTACCGCTGTTGAGATCCAGTTCGATGTAACCCACTCGTGCACCCAACTGATCTTCAGCATCTTTTACTTTCACCAGCGTTTCTGGGTGAGCAAGAACAGGAAGGCACAATGCCGCAGACAAGGGAATAAGGGCGACACGGAAATGTTGAATACTCATACTCTTCCTGCTTCAATAGTATTGAAGCATCTATCAGGGTTAGTGTCTCATGAGCGGATACATATCTGAATGTATGTAGAACACTAGACACATAGGGGTTCCGCGCACATTTCCCCGAGAAGTGCCACCTGACGTCTAAGACACCAGTATCATCATGACATTGACCTCTAGACGGATCCGGGTCGACCGCCACCATGAGCGGCTTTGACGACCCCGGCATCTTCTACAGCGACAGCTTCGGCGGAGATGCCCAGGCCGATGAAGGCCAGGCCAGAAAGTCTCAGCTCCAGCGGCGGTTCAAAGAGTTCCTGCGGCAGTACAGAGTGGGCACCGACAGAACCGGCTTCACCTTCAAGTACCGGGACGAGCTGAAGCGGCACTACAACCTGGGCGAGTACTGGATCGAGGTGGAAATGGAAGATCTGGCCAGCTTCGACGAGGACCTGGCCGACTACCTGTACAAGCAGCCCGCCGAGCATCTCCAGCTGCTGGAAGAGGCCGCTAAAGAGGTGGCCGACGAAGTGACCAGACCCAGGCCTTCTGGCGAGGAAGTGCTCCAGGACATCCAAGTGATGCTGAAGTCCGACGCCAGCCCCAGCAGCATCAGATCCCTGAAGTCTGACATGATGAGCCACCTCGTGAAGATCCCCGGAATCATCATTGCCGCCTCTGCCGTGCGGGCCAAGGCCACCAGAATCAGCATCCAGTGCAGAAGCTGCCGGAACACCCTGACCAATATCGCCATGCGCCCTGGCCTGGAAGGCTACGCCCTGCCCAGAAAGTGCAACACCGACCAGGCTGGCAGACCCAAGTGCCCCCTGGACCCCTACTTCATCATGCCCGACAAGTGCAAATGCGTGGACTTCCAGACCCTGAAACTCCAGGAACTGCCCGACGCCGTGCCTCACGGCGAAATGCCTAGACACATGCAGCTGTACTGCGACAGATACCTGTGCGACAAGGTGGTGCCCGGCAACAGAGTGACCATCATGGGCATCTACAGCATCAAGAAGTTCGGCCTGACCACCAGCCGGGGCAGAGATAGAGTGGGCGTGGGCATCCGGTCCAGCTACATCAGAGTGCTGGGCATCCAGGTGGACACCGATGGCAGCGGCAGATCCTTTGCTGGGGCCGTGAGCCCCCAGGAAGAGGAAGAGTTTAGACGGCTGGCCGCCCTGCCTAACGTGTACGAAGTGATCAGCAAGTCTATCGCCCCCAGCATCTTCGGCGGCACCGATATGAAGAAGGCCATTGCCTGCCTGCTGTTCGGCGGCTCCAGAAAGAGACTGCCTGACGGCCTGACAAGACGGGGCGACATCAACCTGCTGATGCTGGGCGATCCTGGCACCGCCAAATCACAGCTGCTGAAGTTCGTGGAAAAGTGCAGCCCCATCGGCGTGTACACCAGCGGCAAGGGATCTTCTGCCGCCGGACTGACAGCCAGCGTGATGAGAGATCCCAGCAGCCGGAATTTCATCATGGAAGGCGGCGCTATGGTGCTGGCCGATGGCGGAGTCGTGTGCATCGACGAGTTCGACAAGATGCGCGAGGACGACCGGGTGGCCATTCACGAGGCTATGGAACAGCAGACCATCTCTATCGCCAAGGCCGGAATCACCACCACCCTGAACAGCCGGTGTAGCGTGCTGGCTGCCGCCAATAGCGTGTTCGGCAGATGGGACGAGACAAAGGGCGAGGACAACATCGACTTCATGCCCACCATCCTGAGCCGCTTCGACATGATCTTCATCGTGAAGGACGAGCACAACGAGGAACGGGACGTGATGCTGGCCAAGCACGTGATCACCCTGCACGTGTCCGCCCTGACACAGACACAGGCCGTGGAAGGCGAGATCGACCTGGCCAAACTGAAGAAGTTTATCGCCTACTGTCGCGTGAAGTGCGGCCCCAGACTGTCTGCCGAGGCCGCCGAGAAGCTGAAGAACCGGTACATCATCATGCGGAGCGGAGCCCGGCAGCACGAGAGAGACAGCGATCGGAGAAGCAGCATCCCCATCACCGTGCGGCAGCTGGAAGCCATCGTGCGGATTGCCGAAGCCCTGAGCAAGATGAAGCTCCAGCCCTTCGCCACCGAGGCCGACGTGGAAGAAGCTCTGAGACTGTTCCAGGTGTCAACCCTGGACGCCGCCCTGAGCGGAACACTGTCTGGCGTGGAAGGATTCACCAGCCAGGAAGATCAGGAAATGCTGAGCAGAATCGAGAAACAGCTGAAGAGAAGATTCGCCATCGGAAGCCAGGTGTCCGAGCACAGCATCATCAAGGACTTCACCAAGCAGAAGTACCCCGAGCACGCCATCCACAAGGTGCTCCAGCTGATGCTGAGAAGAGGCGAGATCCAGCACCGGATGCAGCGGAAGGTGCTGTACCGGCTGAAATGAGGCGCCGGAATCTTTGCAGGTGCTTACGCTTACTACCTAAACTACAATGGTGTTGTCGCTACTAGTGCCGCTTCTTCATCCACTGCATCTGGTGCTTCCGCTTCCGTTACCGGTTCTAAGAAACTTCAGAAACTTCAGAAACCAGTGCTGCCCCTAAGACATACACTACTGCCACTGTTACTCAATGTGATGACAATGGTTGTAACGTCAAGATAATCACCTCTCAAATACCTGAAGCTACTTCAACCGTCACCGCAACTAGTGCTTCTCCAAAGTCATACACTACTGTCACTTCTGAGGGTTCTAAAGCAACCTCATTGAGTTCTCACTGCCCGCTTTCCAGTCGGGAAACCTGTCGTGCCAGCTGCATTAATTAATCGGCCAACGCGCGGGGAGAGGCGGTTTGCGTATTGGGCGCTCTTCCGCTTCCTCGCTCACTGACTCGCTGCGCTCGGTCGTTCGGCTGCGGCGAGCGGTATCAGCTCACTCAAAGGCGGTAATACGGTTATCCACAGAATCAGGGGATAACGCAGGACAGAACATGTGAGCAGAAGGCCAGCACGAGGCCAGGAGCCGTAAGAAGGCCGCGTTGCTGGCGTTTTTCCATAGGCTCCGCCCCCTCGCGAGCTGAGGACTTAAGTGCTGAGGTCGCGAGCATTCAGAAGAGCCATTCAAGGGATCGATGTGGTCGCACGACTACGCATCCCTCTGGGCCTTACATAGCCGGATACAGTGACTTATCGATACGCGTGATGCCTTGCCTGCAGAGCCGCGTGTTAACCTCGAGGCGGCCGCCATGGTGACAGGACTTGCATAGCTGCGTGCGGGGGAAGGAACTCTTGCGTGTGAGTATGTAGACCCCTGTACTACGGATGCGGGCAGAAGATGTGGGCAGAGACACCCGCGTCAAGTTCTCGACCTTCCCGTGGGAGGTGTTCCAGTCCGCCATACGACCATACCGTTCGGGCATGGCACTATGTACGCGATATCGCTAGCGTTTGCGGGGCACAGCAATATGCAGGCATGCAAGCTTGGCGTAATCATGGTCATAGCTGTTTCCTGTGTGAAATTGTTATCCGCTCACAATTCCACACAACATACGAGCCGGAAGCATAAAGTGTAAAGCCTGGGGTGCCTAATGAGTGAGCTAACTCACATTAATTGCGTTGCGCTCACTGCCCGCTTTCCAGTCGGGAAACCTGTCGTGCCAGCTGCATTAATGAATCGGCCAACGCGCGGGGAGAGGCGGTTTGCGTATTGGGCGCTCTTCCGCTTCCTCGCTCACTGACTCGCTGCGCTCGGTCGTTCGGCTGCGGCGAGCGGTATCAGCTCACTCAAAGGCGGTAATACGGTTATCCACAGAATCAGGGGATAACGCAGGAAAGAACATGTGAGCAAAAGGCCAGCAAAAGGCCAGGAACCGTAAAAAGGCCGCGTTGCTGGCGTTTTTCCATAGGCTCCGCCCCCCTGACGAGCATCACAAAAATCGACGCTCAAGTCAGAGGTGGCGAAACCCGACAGGACTATAAAGATACCAGGCGTTTCCCCCTGGAAGCTCCCTCGTGCGCTCTCCTGTTCCGACCCTGCCGCTTACCGGATACCTGTCCGCCTTTCTCCCTTCGGGAAGCGTGGCGCTTTCTCATAGCTCACGCTGTAGGTATCTCAGTTCGGTGTAGGTCGTTCGCTCCAAGCTGGGCTGTGTGCACGAACCCCCCGTTCAGCCCGACCGCTGCGCCTTATCCGGTAACTATCGTCTTGAGTCCAACCCGGTAAGACACGACTTATCGCCACTGGCAGCAGCCACTGGTAACAGGATTAGCAGAGCGAGGTATGTAGGCGGTGCTACAGAGTTCTTGAAGTGGTGGCCTAACTACGGCTACACTAGAAGAACAGTATTTGGTATCTGCGCTCTGCTGAAGCCAGTTACCTTCGGAAAAAGAGTTGGTAGCTCTTGATCCGGCAAACAAACCACCGCTGGTAGCGGTGGTTTTTTT

**‘Short’ telomeric**

CGAAAGGGCCTCGTGATACGCCTATTTTTATAGGTTAATGTCATGATAATAATGGTTTCTTAGACGTCAGGTGGCACTTTTCGGGGAAATGTGCGCGGAACCCCTATTTGTTTATTTTTCTAAATACATTCAAATATGTATCCGCTCATGAGACAATAACCCTGATAAATGCTTCAATAATATTGAAAAAGGAAGAGTATGAGTATTCAACATTTCCGTGTCGCCCTTATTCCCTTTTTTGCGGCATTTTGCCTTCCTGTTTTTGCTCACCCAGAAACGCTGGTGAAAGTAAAAGATGCTGAAGATCAGTTGGGTGCACGAGTGGGTTACATCGAACTGGATCTCAACAGCGGTAAGATCCTTGAGAGTTTTCGCCCCGAAGAACGTTTTCCAATGATGAGCACTTTTAAAGTTCTGCTATGTGGCGCGGTATTATCCCGTATTGACGCCGGGCAAGAGCAACTCGGTCGCCGCATACACTATTCTCAGAATGACTTGGTTGAGTACTCACCAGTCACAGAAAAGCATCTTACGGATGGCATGACAGTAAGAGAATTATGCAGTGCTGCCATAACCATGAGTGATAACACTGCGGCCAACTTACTTCTGACAACGATCGGAGGACCGAAGGAGCTAACCGCTTTTTTGCACAACATGGGGGATCATGTAACTCGCCTTGATCGTTGGGAACCGGAGCTGAATGAAGCCATACCAAACGACGAGCGTGACACCACGATGCCTGTAGCAATGGCAACAACGTTGCGCAAACTATTAACTGGCGAACTACTTACTCTAGCTTCCCGGCAACAATTAATAGACTGGATGGAGGCGGATAAAGTTGCAGGACCACTTCTGCGCTCGGCCCTTCCGGCTGGCTGGTTTATTGCTGATAAATCTGGAGCCGGTGAGCGTGGGTCTCGCGGTATCATTGCAGCACTGGGGCCAGATGGTAAGCCCTCCCGTATCGTAGTTATCTACACGACGGGGAGTCAGGCAACTATGGATGAACGAAATAGACAGATCGCTGAGATAGGTGCCTCACTGATTAAGCATTGGTAACTGTCAGACCAAGTTTACTCATATATACTTTAGATTGATTTAAAACTTCATTTTTAATTTAAAAGGATCTAGGTGAAGATCCTTTTTGATAATCTCATGACCAAAATCCCTTAACGTGAGTTTTCGTTCCACTGAGCGTCAGACCCCGTAGAAAAGATCAAAGGATCTTCTTGAGATCCTTTTTTTCTGCGCGTAATCTGCTGCTTGCAAACAAAAAAACCACCGCTACCAGCGGTGGTTTGTTTGCCGGATCAAGAGCTACCAACTCTTTTTCCGAAGGTAACTGGCTTCAGCAGAGCGCAGATACCAAATACTGTTCTTCTAGTGTAGCCGTAGTTAGGCCACCACTTCAAGAACTCTGTAGCACCGCCTACATACCTCGCTCTGCTAATCCTGTTACCAGTGGCTGCTGCCAGTGGCGATAAGTCGTGTCTTACCGGGTTGGACTCAAGACGATAGTTACCGGATAAGGCGCAGCGGTCGGGCTGAACGGGGGGTTCGTGCACACAGCCCAGCTTGGAGCGAACGACCTACACCGAACTGAGATACCTACAGCGTGAGCTATGAGAAAGCGCCACGCTTCCCGAAGGGAGAAAGGCGGACAGGTATCCGGTAAGCGGCAGGGTCGGAACAGGAGAGCGCACGAGGGAGCTTCCAGGGGGAAACGCCTGGTATCTTTATAGTCCTGTCGGGTTTCGCCACCTCTGACTTGAGCGTCGATTTTTGTGATGCTCGTCAGGGGGGCGGAGCCTATGGAAAAACGCCAGCAACGCGGCCTTTTTACGGTTCCTGGCCTTTTGCTGGCCTTTTGCTCACATGTTCTTTCCTGCGTTATCCCCTGATTCTGTGGATAACCGTATTACCGCCTTTGAGTGAGCTGATACCGCTCGCCGCAGCCGAACGACCGAGCGCAGCGAGTCAGTGAGCGAGGAAGCGGAAGAGCGCCCAATACGCAAACCGCCTCTCCCCGCGCGTTGGCCGATTCATTAATGCAGCTGGCACGACAGGTTTCCCGACTGGAAAGCGGGCAGTGAGCGCAACGCAATTAATGTGAGTTAGCTCACTCATTAGGCACCCCAGGCTTTACACTTTATGCTTCCGGCTCGTATGTTGTGTGGAATTGTGAGCGGATAACAATTTCACACAGGAAACAGCTATGACCATGATTACGCCAAGCTTGCATGCCTGCATATTGCTGTGCCCCGCAAACGCTAGCGATATCGCGTACATAGTGCCATGCCCGAACGGTATGGTCGTATGGCGGACTGGAACACCTCCCACGGGAAGGTCGAGAACTTGACGCGGGTGTCTCTGCCCACATCTTCTGCCCGCATCCGTAGTACAGGGGTCTACATACTCACACGCAAGAGTTCCTTCCCCCGCACGCAGCTATGCAAGTCCTGTCACCATGGCGGCCGCCTCGAGGTTAACACGCGGCTCTGCAGGCAAGGCATCACGCGTATCGATAAGTCACTGTATCCGGCTATGTAAGGCCCAGAGGGATGCGTAGTCGTGCGACCACATCGATCCCTTGAATGGCTCTTCTGAATGCTCGCGACCTCAGCACTTAAGTCCTCAGCTCGCGAGGGGGCGGAGCCTATGGAAAAACGCCAGCAACGCGGCCTTCTTACGGCTCCTGGCCTCGTGCTGGCCTTCTGCTCACATGTTCTGTCCTGCGTTATCCCCTGATTCTGTGGATAACCGTATTACCGCCTTTGAGTGAGCTGATACCGCTCGCCGCAGCCGAACGACCGAGCGCAGCGAGTCAGTGAGCGAGGAAGCGGAAGAGCGCCCAATACGCAAACCGCCTCTCCCCGCGCGTTGGCCGATTCATTAATGCAGCTGGCACGACAGGTTTCCCGACTGGAAAGCGGGCAGTGAGAACTCAATGAGGTTGCTTTAGAACCCTCAGAAGTGACAGTAGTGTATGACTTTGGAGAAGCACTAGTTGCGGTGACGGTTGAAGTAGCTTCAGGTATTTGAGAGGTGATTATCTTGACGTTACAACCATTGTCATCACATTGAGTAACAGTGGCAGTAGTGTATGTCTTAGGGGCAGCACTGGTTTCTGAAGTTTCTGAAGTTTCTTAGAACCGGTAACGGAAGCGGAAGCACCAGATGCAGTGGATGAAGAAGCGGCACTAGTAGCGACAACACCATTGTAGTTTAGGTAGTAAGCGTAAGCACCTGCAAAGATTCCGGCGCCTCATTTCAGCCGGTACAGCACCTTCCGCTGCATCCGGTGCTGGATCTCGCCTCTTCTCAGCATCAGCTGGAGCACCTTGTGGATGGCGTGCTCGGGGTACTTCTGCTTGGTGAAGTCCTTGATGATGCTGTGCTCGGACACCTGGCTTCCGATGGCGAATCTTCTCTTCAGCTGTTTCTCGATTCTGCTCAGCATTTCCTGATCTTCCTGGCTGGTGAATCCTTCCACGCCAGACAGTGTTCCGCTCAGGGCGGCGTCCAGGGTTGACACCTGGAACAGTCTCAGAGCTTCTTCCACGTCGGCCTCGGTGGCGAAGGGCTGGAGCTTCATCTTGCTCAGGGCTTCGGCAATCCGCACGATGGCTTCCAGCTGCCGCACGGTGATGGGGATGCTGCTTCTCCGATCGCTGTCTCTCTCGTGCTGCCGGGCTCCGCTCCGCATGATGATGTACCGGTTCTTCAGCTTCTCGGCGGCCTCGGCAGACAGTCTGGGGCCGCACTTCACGCGACAGTAGGCGATAAACTTCTTCAGTTTGGCCAGGTCGATCTCGCCTTCCACGGCCTGTGTCTGTGTCAGGGCGGACACGTGCAGGGTGATCACGTGCTTGGCCAGCATCACGTCCCGTTCCTCGTTGTGCTCGTCCTTCACGATGAAGATCATGTCGAAGCGGCTCAGGATGGTGGGCATGAAGTCGATGTTGTCCTCGCCCTTTGTCTCGTCCCATCTGCCGAACACGCTATTGGCGGCAGCCAGCACGCTACACCGGCTGTTCAGGGTGGTGGTGATTCCGGCCTTGGCGATAGAGATGGTCTGCTGTTCCATAGCCTCGTGAATGGCCACCCGGTCGTCCTCGCGCATCTTGTCGAACTCGTCGATGCACACGACTCCGCCATCGGCCAGCACCATAGCGCCGCCTTCCATGATGAAATTCCGGCTGCTGGGATCTCTCATCACGCTGGCTGTCAGTCCGGCGGCAGAAGATCCCTTGCCGCTGGTGTACACGCCGATGGGGCTGCACTTTTCCACGAACTTCAGCAGCTGTGATTTGGCGGTGCCAGGATCGCCCAGCATCAGCAGGTTGATGTCGCCCCGTCTTGTCAGGCCGTCAGGCAGTCTCTTTCTGGAGCCGCCGAACAGCAGGCAGGCAATGGCCTTCTTCATATCGGTGCCGCCGAAGATGCTGGGGGCGATAGACTTGCTGATCACTTCGTACACGTTAGGCAGGGCGGCCAGCCGTCTAAACTCTTCCTCTTCCTGGGGGCTCACGGCCCCAGCAAAGGATCTGCCGCTGCCATCGGTGTCCACCTGGATGCCCAGCACTCTGATGTAGCTGGACCGGATGCCCACGCCCACTCTATCTCTGCCCCGGCTGGTGGTCAGGCCGAACTTCTTGATGCTGTAGATGCCCATGATGGTCACTCTGTTGCCGGGCACCACCTTGTCGCACAGGTATCTGTCGCAGTACAGCTGCATGTGTCTAGGCATTTCGCCGTGAGGCACGGCGTCGGGCAGTTCCTGGAGTTTCAGGGTCTGGAAGTCCACGCATTTGCACTTGTCGGGCATGATGAAGTAGGGGTCCAGGGGGCACTTGGGTCTGCCAGCCTGGTCGGTGTTGCACTTTCTGGGCAGGGCGTAGCCTTCCAGGCCAGGGCGCATGGCGATATTGGTCAGGGTGTTCCGGCAGCTTCTGCACTGGATGCTGATTCTGGTGGCCTTGGCCCGCACGGCAGAGGCGGCAATGATGATTCCGGGGATCTTCACGAGGTGGCTCATCATGTCAGACTTCAGGGATCTGATGCTGCTGGGGCTGGCGTCGGACTTCAGCATCACTTGGATGTCCTGGAGCACTTCCTCGCCAGAAGGCCTGGGTCTGGTCACTTCGTCGGCCACCTCTTTAGCGGCCTCTTCCAGCAGCTGGAGATGCTCGGCGGGCTGCTTGTACAGGTAGTCGGCCAGGTCCTCGTCGAAGCTGGCCAGATCTTCCATTTCCACCTCGATCCAGTACTCGCCCAGGTTGTAGTGCCGCTTCAGCTCGTCCCGGTACTTGAAGGTGAAGCCGGTTCTGTCGGTGCCCACTCTGTACTGCCGCAGGAACTCTTTGAACCGCCGCTGGAGCTGAGACTTTCTGGCCTGGCCTTCATCGGCCTGGGCATCTCCGCCGAAGCTGTCGCTGTAGAAGATGCCGGGGTCGTCAAAGCCGCTCATGGTGGCGGTCGACCCGGATCcGTCTAGAGGTCAATGTCATGATGATACTGGTGTCTTAGACGTCAGGTGGCACTTCTCGGGGAAATGTGCGCGGAACCCCTATGTGTCTAGTGTTCTACATACATTCAGATATGTATCCGCTCATGAGACACTAACCCTGATAGATGCTTCAATACTATTGAAGCAGGAAGAGTATGAGTATTCAACATTTCCGTGTCGCCCTTATTCCCTTGTCTGCGGCATTGTGCCTTCCTGTTCTTGCTCACCCAGAAACGCTGGTGAAAGTAAAAGATGCTGAAGATCAGTTGGGTGCACGAGTGGGTTACATCGAACTGGATCTCAACAGCGGTAAGATCCTTGAGAGTTTTCGCCCCGAAGAACGTTTTCCAATGATGAGCACTTTCAAAGTGCTGCTATGTGGCGCGGTACTATCCCGTATTGACGCCGGGCAAGAGCAACTCGGTCGCCGCATACACTATTCTCAGAATGACTCGTGGTTGCAATTGAGCGGGGATCCTCTAGAGTCGACGAAGACTTAGGGTTAGGGTTAGGGTTAGGGTTAGGGTTAGGGTTAGGGTTAGGGTTAGGGTTAGGGTTAGGGTTAGGGTTAGGGTTAGGGTTAGGGTTAGGGTTAGGGTTAGGGTTAGGGTTAGGGTTAGGGTTAGGGTTAGGGTTAGGGTTAGGGTTAGGGTTAGGGTTAGGGTTAGGGTTAGGGTTAGGGTTAGGGTTAGGGTTAGGGTTAGGGTTAGGGTTAGGGTTAGGGTTAGGGTTAGGGTTAGGGTTAGGGTTAGGGTTAGGGTTAGGGTTAGGGTTAGGGTTAGGGTTAGGGTTAGGGTTAGGGTTAGGGTTAGGGTTAGGGTTAGGGTTAGGGTTAGGGTTAGGGTTAGGGTTAGGGCGAGACGTGTCCCGTCGACCTGCATAGCGGGTACCGAGCTCGAATTCACTGGCCGTCGTTTTACAACGTCGTGACTGGGAAAACCCTGGCGTTACCCAACTTAATCGCCTTGCAGCACATCCCCCTTTCGCCAGCTGGCGTAATAGCGAAGAGGCCCGCACCGATCGCCCTTCCCAACAGTTGCGCAGCCTGAATGGCGAATGGCGCCTGATGCGGTATTTTCTCCTTACGCATCTGTGCGGTATTTCACACCGCATATGCCTATAGTGAGTCGTATTATGCGGATGCGCTGGTAGTTCTGGTAGATTGTCTCTTCCATGTTCACTTCGAAGGGTCCGGCGCTCTGACACTCGGGACAGCTGCCGGGCTTCACTTCCTGGTTCTGGCTCTGGCAGAAGGGTCCCAGCACGAAGTTGCACTTGTTGCAGTTGTACTTGACCATAGACAGCTGGGGCAGCACGCCTGTACAAGAGGTCACGACGCCGGAGGTCCGGATCAGCTGGTTCAGGTGCAGCTGTCTCAGGCTCCGCAGTTCTTCCACCAGGGGCAGATGGCTGATCCGCACGTGGATGTGGTTGGTGATCCGGTCGTACTTAGGGTACATGGCCAGCACCACTTCCAGGGCGGCCTCATCAAAGATCTGGAGCAGCTCGGCAGGGGCCTCAGGCAGAAAGTAGGCCAGCACGTGTTCTCTGGCGGCCAGATCCTCGTAGTTCACGACCAGGCTCTCGCGGTTCTCTTTGCACATGTCGCTGATCCGTTCTTTGAACACGTTGTGGCCGTGGCTGTCCACGTGGGTCCGCAGAAAGTTCTTGAACCGGTGGTGGATTTCCAGTCTGGGGCCAGCCATTGACACCCACTCGCGCACGCTGTGGCCCTTCAGATCTTCCAGGTTCTCGATGCTCTCGATCATCTCTTCATCCTCTTCCCCGTCCTCGGTGGCTCTTTCCACCTGTCTCCGCTTTCTGGCGGGTCTTTCCTCATCTTCCTCGTCGCTGTCGTACAGCAGGCCTCTCCGCATTCTGCCCAGGCCTCTGCCAGCCTCTCTGTCTCTCTGCCGCATAGCTCTCTCGGCGGCCTCTCTCTGGCTGGCGGTCAGTTCTTCCACATCCTCGTCGTCCAGGGCCAGGCCTTCGGCCTCATAGGCATCCAGCTCGGGGATGGCTCTGTAGTCCCGTTCCATGCCGTCGCCGATCAGTTCCTCGCCGTCCTCTTCTTCTTCCAGAGGTCCCTCTGTGCCCAGCAGGCCCTCGCTCTCATCCTCGAATGGGGGCAGATCTCTGCCAGGGGAGCTGGTCAGGGCGTCGGTTCTTCTGCTGCTTCTGCCGGGGCTGCTTGTCAGAGGGTCGTTGCCTCTTCTCCGCTGGGCAGGGCTAGAGGTCCGGGAGCTGCATGTGTCAGAGG

**LacO**

CCTCTGACACATGCAGCTCCCGGACCTCTAGCCCTGCCCAGCGGAGAAGAGGCAACGACCCTCTGACAAGCAGCCCCGGCAGAAGCAGCAGAAGAACCGACGCCCTGACCAGCTCCCCTGGCAGAGATCTGCCCCCATTCGAGGATGAGAGCGAGGGCCTGCTGGGCACAGAGGGACCTCTGGAAGAAGAAGAGGACGGCGAGGAACTGATCGGCGACGGCATGGAACGGGACTACAGAGCCATCCCCGAGCTGGATGCCTATGAGGCCGAAGGCCTGGCCCTGGACGACGAGGATGTGGAAGAACTGACCGCCAGCCAGAGAGAGGCCGCCGAGAGAGCTATGCGGCAGAGAGACAGAGAGGCTGGCAGAGGCCTGGGCAGAATGCGGAGAGGCCTGCTGTACGACAGCGACGAGGAAGATGAGGAAAGACCCGCCAGAAAGCGGAGACAGGTGGAAAGAGCCACCGAGGACGGGGAAGAGGATGAAGAGATGATCGAGAGCATCGAGAACCTGGAAGATCTGAAGGGCCACAGCGTGCGCGAGTGGGTGTCAATGGCTGGCCCCAGACTGGAAATCCACCACCGGTTCAAGAACTTTCTGCGGACCCACGTGGACAGCCACGGCCACAACGTGTTCAAAGAACGGATCAGCGACATGTGCAAAGAGAACCGCGAGAGCCTGGTCGTGAACTACGAGGATCTGGCCGCCAGAGAACACGTGCTGGCCTACTTTCTGCnCTGAGGCCCCTGCCGAGCTGCTCCAGATCTTTGATGAGGCCGCCCTGGAAGTGGTGCTGGCCATGTACCCTAAGTACGACCGGATCACCAACCACATCCACGTGCGGATCAGCCATCTGCCCCTGGTGGAAGAACTGCGGAGCCTGAGACAGCTGCACCTGAACCAGCTGATCCGGACCTCCGGCGTCGTGACCTCTTGTACAGGCGTGCTGCCCCAGCTGTCTATGGTCAAGTACAACTGCAACAAGTGCAACTTCGTGCTGGGACCCTTCTGCCAGAGCCAGAACCAGGAAGTGAAGCCCGGCAGCTGTCCCGAGTGTCAGAGCGCCGGACCCTTCGAAGTGAACATGGAAGAGACAATCTACCAGAACTACCAGCGCATCCGCATAATACGACTCACTATAGGCATATGCGGTGTGAAATACCGCACAGATGCGTAAGGAGAAAATACCGCATCAGGCGCCATTCGCCATTCAGGCTGCGCAACTGTTGGGAAGGGCGATCGGTGCGGGCCTCTTCGCTATTACGCCAGCTGGCGAAAGGGGGATGTGCTGCAAGGCGATTAAGTTGGGTAACGCCAGGGTTTTCCCAGTCACGACGTTGTAAAACGACGGCCAGTGaattcCTCGATTTTTTTATGTTTAGTTTCGCGGACGACGGTTTCGAGGTGGCGGTCTGGACCACGCCGGAGAGCGTCGAAGCGGAGGCGGTGTTCGCCGAGATCGGGAGCTCTCACACCTACAAGGGATGTACATCAATTGTGAGCGGATAACAATTGTTAGGGAGGAATTGTGAGCGGATAACAATTTGGAGTTGATAATTGTGAGCGGATAACAATTGGCTTCAACGTAATTGTGAGCGGATAACAATTTCCGTACGAATGTGCCGAACTTATGGTACCGCCGAGCGCGACGGTGTGCCCGCCTTCCTGGAGACCTCCGCGCTCCGCAACCTCCACTTCTAAAAGCGGCTCGGCTTCAACTAAACATAAAAAtACGAGGTGCCCGAAGGACCGCGCACCTGGTGCATGACCCGCAAGCCCGGTGCCTGACGCTCGCCACACGACCCGCAGCGCCCGACCGAAAGGAGCGCACGAcccgggGATCCGGGTCGACCGCCACCATGAGCGGCTTTGACGACCCCGGCATCTTCTACAGCGACAGCTTCGGCGGAGATGCCCAGGCCGATGAAGGCCAGGCCAGAAAGTCTCAGCTCCAGCGGCGGTTCAAAGAGTTCCTGCGGCAGTACAGAGTGGGCACCGACAGAACCGGCTTCACCTTCAAGTACCGGGACGAGCTGAAGCGGCACTACAACCTGGGCGAGTACTGGATCGAGGTGGAAATGGAAGATCTGGCCAGCTTCGACGAGGACCTGGCCGACTACCTGTACAAGCAGCCCGCCGAGCATCTCCAGCTGCTGGAAGAGGCCGCTAAAGAGGTGGCCGACGAAGTGACCAGACCCAGGCCTTCTGGCGAGGAAGTGCTCCAGGACATCCAAGTGATGCTGAAGTCCGACGCCAGCCCCAGCAGCATCAGATCCCTGAAGTCTGACATGATGAGCCACCTCGTGAAGATCCCCGGAATCATCATTGCCGCCTCTGCCGTGCGGGCCAAGGCCACCAGAATCAGCATCCAGTGCAGAAGCTGCCGGAACACCCTGACCAATATCGCCATGCGCCCTGGCCTGGAAGGCTACGCCCTGCCCAGAAAGTGCAACACCGACCAGGCTGGCAGACCCAAGTGCCCCCTGGACCCCTACTTCATCATGCCCGACAAGTGCAAATGCGTGGACTTCCAGACCCTGAAACTCCAGGAACTGCCCGACGCCGTGCCTCACGGCGAAATGCCTAGACACATGCAGCTGTACTGCGACAGATACCTGTGCGACAAGGTGGTGCCCGGCAACAGAGTGACCATCATGGGCATCTACAGCATCAAGAAGTTCGGCCTGACCACCAGCCGGGGCAGAGATAGAGTGGGCGTGGGCATCCGGTCCAGCTACATCAGAGTGCTGGGCATCCAGGTGGACACCGATGGCAGCGGCAGATCCTTTGCTGGGGCCGTGAGCCCCCAGGAAGAGGAAGAGTTTAGACGGCTGGCCGCCCTGCCTAACGTGTACGAAGTGATCAGCAAGTCTATCGCCCCCAGCATCTTCGGCGGCACCGATATGAAGAAGGCCATTGCCTGCCTGCTGTTCGGCGGCTCCAGAAAGAGACTGCCTGACGGCCTGACAAGACGGGGCGACATCAACCTGCTGATGCTGGGCGATCCTGGCACCGCCAAATCACAGCTGCTGAAGTTCGTGGAAAAGTGCAGCCCCATCGGCGTGTACACCAGCGGCAAGGGATCTTCTGCCGCCGGACTGACAGCCAGCGTGATGAGAGATCCCAGCAGCCGGAATTTCATCATGGAAGGCGGCGCTATGGTGCTGGCCGATGGCGGAGTCGTGTGCATCGACGAGTTCGACAAGATGCGCGAGGACGACCGGGTGGCCATTCACGAGGCTATGGAACAGCAGACCATCTCTATCGCCAAGGCCGGAATCACCACCACCCTGAACAGCCGGTGTAGCGTGCTGGCTGCCGCCAATAGCGTGTTCGGCAGATGGGACGAGACAAAGGGCGAGGACAACATCGACTTCATGCCCACCATCCTGAGCCGCTTCGACATGATCTTCATCGTGAAGGACGAGCACAACGAGGAACGGGACGTGATGCTGGCCAAGCACGTGATCACCCTGCACGTGTCCGCCCTGACACAGACACAGGCCGTGGAAGGCGAGATCGACCTGGCCAAACTGAAGAAGTTTATCGCCTACTGTCGCGTGAAGTGCGGCCCCAGACTGTCTGCCGAGGCCGCCGAGAAGCTGAAGAACCGGTACATCATCATGCGGAGCGGAGCCCGGCAGCACGAGAGAGACAGCGATCGGAGAAGCAGCATCCCCATCACCGTGCGGCAGCTGGAAGCCATCGTGCGGATTGCCGAAGCCCTGAGCAAGATGAAGCTCCAGCCCTTCGCCACCGAGGCCGACGTGGAAGAAGCTCTGAGACTGTTCCAGGTGTCAACCCTGGACGCCGCCCTGAGCGGAACACTGTCTGGCGTGGAAGGATTCACCAGCCAGGAAGATCAGGAAATGCTGAGCAGAATCGAGAAACAGCTGAAGAGAAGATTCGCCATCGGAAGCCAGGTGTCCGAGCACAGCATCATCAAGGACTTCACCAAGCAGAAGTACCCCGAGCACGCCATCCACAAGGTGCTCCAGCTGATGCTGAGAAGAGGCGAGATCCAGCACCGGATGCAGCGGAAGGTGCTGTACCGGCTGAAATGAGGCGCCGGAATCTTTGCAGGTGCTTACGCTTACTACCTAAACTACAATGGTGTTGTCGCTACTAGTGCCGCTTCTTCATCCACTGCATCTGGTGCTTCCGCTTCCGTTACCGGTTCTAAGAAACTTCAGAAACTTCAGAAACCAGTGCTGCCCCTAAGACATACACTACTGCCACTGTTACTCAATGTGATGACAATGGTTGTAACGTCAAGATAATCACCTCTCAAATACCTGAAGCTACTTCAACCGTCACCGCAACTAGTGCTTCTCCAAAGTCATACACTACTGTCACTTCTGAGGGTTCTAAAGCAACCTCATTGAGTTCTCACTGCCCGCTTTCCAGTCGGGAAACCTGTCGTGCCAGCTGCATTAATGAATCGGCCAACGCGCGGGGAGAGGCGGTTTGCGTATTGGGCGCTCTTCCGCTTCCTCGCTCACTGACTCGCTGCGCTCGGTCGTTCGGCTGCGGCGAGCGGTATCAGCTCACTCAAAGGCGGTAATACGGTTATCCACAGAATCAGGGGATAACGCAGGACAGAACATGTGAGCAGAAGGCCAGCACGAGGCCAGGAGCCGTAAGAAGGCCGCGTTGCTGGCGTTTTTCCATAGGCTCCGCCCCCTCGCGAGCTGAGGACTTAAGTGCTGAGGTCGCGAGCATTCAGAAGAGCCATTCAAGGGATCGATGTGGTCGCACGACTACGCATCCCTCTGGGCCTTACATAGCCGGATACAGTGACTTATCGATACGCGTGATGCCTTGCCTGCAGAGCCGCGTGTTAACCTCGAGGCGGCCGCCATGGTGACAGGACTTGCATAGCTGCGTGCGGGGGAAGGAACTCTTGCGTGTGAGTATGTAGACCCCTGTACTACGGATGCGGGCAGAAGATGTGGGCAGAGACACCCGCGTCAAGTTCTCGACCTTCCCGTGGGAGGTGTTCCAGTCCGCCATACGACCATACCGTTCGGGCATGGCACTATGTACGCGATATCGCTAGCGTTTGCGGGGCACAGCAATATGCAGGCATGCAAGCTTGGCGTAATCATGGTCATAGCTGTTTCCTGTGTGAAATTGTTATCCGCTCACAATTCCACACAACATACGAGCCGGAAGCATAAAGTGTAAAGCCTGGGGTGCCTAATGAGTGAGCTAACTCACATTAATTGCGTTGCGCTCACTGCCCGCTTTCCAGTCGGGAAACCTGTCGTGCCAGCTGCATTAATGAATCGGCCAACGCGCGGGGAGAGGCGGTTTGCGTATTGGGCGCTCTTCCGCTTCCTCGCTCACTGACTCGCTGCGCTCGGTCGTTCGGCTGCGGCGAGCGGTATCAGCTCACTCAAAGGCGGTAATACGGTTATCCACAGAATCAGGGGATAACGCAGGAAAGAACATGTGAGCAAAAGGCCAGCAAAAGGCCAGGAACCGTAAAAAGGCCGCGTTGCTGGCGTTTTTCCATAGGCTCCGCCCCCCTGACGAGCATCACAAAAATCGACGCTCAAGTCAGAGGTGGCGAAACCCGACAGGACTATAAAGATACCAGGCGTTTCCCCCTGGAAGCTCCCTCGTGCGCTCTCCTGTTCCGACCCTGCCGCTTACCGGATACCTGTCCGCCTTTCTCCCTTCGGGAAGCGTGGCGCTTTCTCATAGCTCACGCTGTAGGTATCTCAGTTCGGTGTAGGTCGTTCGCTCCAAGCTGGGCTGTGTGCACGAACCCCCCGTTCAGCCCGACCGCTGCGCCTTATCCGGTAACTATCGTCTTGAGTCCAACCCGGTAAGACACGACTTATCGCCACTGGCAGCAGCCACTGGTAACAGGATTAGCAGAGCGAGGTATGTAGGCGGTGCTACAGAGTTCTTGAAGTGGTGGCCTAACTACGGCTACACTAGAAGAACAGTATTTGGTATCTGCGCTCTGCTGAAGCCAGTTACCTTCGGAAAAAGAGTTGGTAGCTCTTGATCCGGCAAACAAACCACCGCTGGTAGCGGTGGTTTTTTTGTTTGCAAGCAGCAGATTACGCGCAGAAAAAAAGGATCTCAAGAAGATCCTTTGATCTTTTCTACGGGGTCTGACGCTCAGTGGAACGAAAACTCACGTTAAGGGATTTTGGTCATGAGATTATCAAAAAGGATCTTCACCTAGATCCTTTTAAATTAAAAATGAAGTTTTAAATCAATCTAAAGTATATATGAGTAAACTTGGTCTGACAGTTACCAATGCTTAATCAGTGAGGCACCTATCTCAGCGATCTGTCTATTTCGTTCATCCATAGTTGCCTGACTCCCCGTCGTGTAGATAACTACGATACGGGAGGGCTTACCATCTGGCCCCAGTGCTGCAATGATACCGCGAGACCCACGCTCACCGGCTCCAGATTTATCAGCAATAAACCAGCCAGCCGGAAGGGCCGAGCGCAGAAGTGGTCCTGCAACTTTATCCGCCTCCATCCAGTCTATTAATTGTTGCCGGGAAGCTAGAGTAAGTAGTTCGCCAGTTAATAGTTTGCGCAACGTTGTTGCCATTGCTACAGGCATCGTGGTGTCACGCTCGTCGTTTGGTATGGCTTCATTCAGCTCCGGTTCCCAACGATCAAGGCGAGTTACATGATCCCCCATGTTGTGCAAAAAAGCGGTTAGCTCCTTCGGTCCTCCGATCGTTGTCAGAAGTAAGTTGGCCGCAGTGTTATCACTCATGGTTATGGCAGCACTGCATAATTCTCTTACTGTCATGCCATCCGTAAGATGCTTTTCTGTGACTGGTGAGTACTCAACCAAGTCATTCTGAGAATAGTGTATGCGGCGACCGAGTTGCTCTTGCCCGGCGTCAATACGGGATAATACCGCGCCACATAGCAGAACTTTAAAAGTGCTCATCATTGGAAAACGTTCTTCGGGGCGAAAACTCTCAAGGATCTTACCGCTGTTGAGATCCAGTTCGATGTAACCCACTCGTGCACCCAACTGATCTTCAGCATCTTTTACTTTCACCAGCGTTTCTGGGTGAGCAAAAACAGGAAGGCAAAATGCCGCAAAAAAGGGAATAAGGGCGACACGGAAATGTTGAATACTCATACTCTTCCTTTTTCAATATTATTGAAGCATTTATCAGGGTTATTGTCTCATGAGCGGATACATATTTGAATGTATTTAGAAAAATAAACAAATAGGGGTTCCGCGCACATTTCCCCGAAAAGTGCCACCTGACGTCTAAGAAACCATTATTATCATGACATTAACCTATAAAAATAGGCGTATCACGAGGCCCTTTCG

**‘Long’ control (Figure S1a and b)**

CCTCTGACACATGCAGCTCCCGGACCTCTAGCCCTGCCCAGCGGAGAAGAGGCAACGACCCTCTGACAAGCAGCCCCGGCAGAAGCAGCAGAAGAACCGACGCCCTGACCAGCTCCCCTGGCAGAGATCTGCCCCCATTCGAGGATGAGAGCGAGGGCCTGCTGGGCACAGAGGGACCTCTGGAAGAAGAAGAGGACGGCGAGGAACTGATCGGCGACGGCATGGAACGGGACTACAGAGCCATCCCCGAGCTGGATGCCTATGAGGCCGAAGGCCTGGCCCTGGACGACGAGGATGTGGAAGAACTGACCGCCAGCCAGAGAGAGGCCGCCGAGAGAGCTATGCGGCAGAGAGACAGAGAGGCTGGCAGAGGCCTGGGCAGAATGCGGAGAGGCCTGCTGTACGACAGCGACGAGGAAGATGAGGAAAGACCCGCCAGAAAGCGGAGACAGGTGGAAAGAGCCACCGAGGACGGGGAAGAGGATGAAGAGATGATCGAGAGCATCGAGAACCTGGAAGATCTGAAGGGCCACAGCGTGCGCGAGTGGGTGTCAATGGCTGGCCCCAGACTGGAAATCCACCACCGGTTCAAGAACTTTCTGCGGACCCACGTGGACAGCCACGGCCACAACGTGTTCAAAGAACGGATCAGCGACATGTGCAAAGAGAACCGCGAGAGCCTGGTCGTGAACTACGAGGATCTGGCCGCCAGAGAACACGTGCTGGCCTACTTTCTGCCTGAGGCCCCTGCCGAGCTGCTCCAGATCTTTGATGAGGCCGCCCTGGAAGTGGTGCTGGCCATGTACCCTAAGTACGACCGGATCACCAACCACATCCACGTGCGGATCAGCCATCTGCCCCTGGTGGAAGAACTGCGGAGCCTGAGACAGCTGCACCTGAACCAGCTGATCCGGACCTCCGGCGTCGTGACCTCTTGTACAGGCGTGCTGCCCCAGCTGTCTATGGTCAAGTACAACTGCAACAAGTGCAACTTCGTGCTGGGACCCTTCTGCCAGAGCCAGAACCAGGAAGTGAAGCCCGGCAGCTGTCCCGAGTGTCAGAGCGCCGGACCCTTCGAAGTGAACATGGAAGAGACAATCTACCAGAACTACCAGCGCATCCGCATAATACGACTCACTATAGGCATATGCGGTGTGAAATACCGCACAGATGCGTAAGGAGAAAATACCGCATCAGGCGCCATTCGCCATTCAGGCTGCGCAACTGTTGGGAAGGGCGATCGGTGCGGGCCTCTTCGCTATTACGCCAGCTGGCGAAAGGGGGATGTGCTGCAAGGCGATTAAGTTGGGTAACGCCAGGGTTTTCCCAGTCACGACGTTGTAAAACGACGGCCAGTGAATTCGAGCTCGGTACCCGGGGATCCTCTAGAGTCGACGAAGACCACCAGCTAGTACAACCACATACTTTATGGAGAAATTTCAAAACGCAAAGAAGAACGAAGATAAACAAATTGCCAAGTTTGAAAGCATGATGAATGCAAGAGTACATACGTTCAGTACCGATGAGAAGAAATATGTGCCGATAATCACAAACGAATTAGAAAGCTTTTCAAATCTTTGGGTTAAAAAGAGGTACATACCTGAAGATGACTTAAAACGGGCTTTGCATGAGATCAAAATCCTTCGGTTGGGCAAACTTTTTGCTAAAATTCGCCCACCTAAATTTCAAGAGCCTGAATACGCCAACTGGGCCACCGTAGGCCTCATTAGCCACAAATCGGACATCAAATTTACATCATCTGAGACGTGTCCCGTCGACCTGCATAGCGCTCAATTGCAACCACTGCAGAGTCATTCTGAGAATAGTGTATGCGGCGACCGAGTTGCTCTTGCCCGGCGTCAATACGGGATAGTACCGCGCCACATAGCAGCACTTTGAAAGTGCTCATCATTGGAAAACGTTCTTCGGGGCGAAAACTCTCAAGGATCTTACCGCTGTTGAGATCCAGTTCGATGTAACCCACTCGTGCACCCAACTGATCTTCAGCATCTTTTACTTTCACCAGCGTTTCTGGGTGAGCAAGAACAGGAAGGCACAATGCCGCAGACAAGGGAATAAGGGCGACACGGAAATGTTGAATACTCATACTCTTCCTGCTTCAATAGTATTGAAGCATCTATCAGGGTTAGTGTCTCATGAGCGGATACATATCTGAATGTATGTAGAACACTAGACACATAGGGGTTCCGCGCACATTTCCCCGAGAAGTGCCACCTGACGTCTAAGACACCAGTATCATCATGACATTGACCTCTAGACgGATCCGGGTCGACCGCCACCATGAGCGGCTTTGACGACCCCGGCATCTTCTACAGCGACAGCTTCGGCGGAGATGCCCAGGCCGATGAAGGCCAGGCCAGAAAGTCTCAGCTCCAGCGGCGGTTCAAAGAGTTCCTGCGGCAGTACAGAGTGGGCACCGACAGAACCGGCTTCACCTTCAAGTACCGGGACGAGCTGAAGCGGCACTACAACCTGGGCGAGTACTGGATCGAGGTGGAAATGGAAGATCTGGCCAGCTTCGACGAGGACCTGGCCGACTACCTGTACAAGCAGCCCGCCGAGCATCTCCAGCTGCTGGAAGAGGCCGCTAAAGAGGTGGCCGACGAAGTGACCAGACCCAGGCCTTCTGGCGAGGAAGTGCTCCAGGACATCCAAGTGATGCTGAAGTCCGACGCCAGCCCCAGCAGCATCAGATCCCTGAAGTCTGACATGATGAGCCACCTCGTGAAGATCCCCGGAATCATCATTGCCGCCTCTGCCGTGCGGGCCAAGGCCACCAGAATCAGCATCCAGTGCAGAAGCTGCCGGAACACCCTGACCAATATCGCCATGCGCCCTGGCCTGGAAGGCTACGCCCTGCCCAGAAAGTGCAACACCGACCAGGCTGGCAGACCCAAGTGCCCCCTGGACCCCTACTTCATCATGCCCGACAAGTGCAAATGCGTGGACTTCCAGACCCTGAAACTCCAGGAACTGCCCGACGCCGTGCCTCACGGCGAAATGCCTAGACACATGCAGCTGTACTGCGACAGATACCTGTGCGACAAGGTGGTGCCCGGCAACAGAGTGACCATCATGGGCATCTACAGCATCAAGAAGTTCGGCCTGACCACCAGCCGGGGCAGAGATAGAGTGGGCGTGGGCATCCGGTCCAGCTACATCAGAGTGCTGGGCATCCAGGTGGACACCGATGGCAGCGGCAGATCCTTTGCTGGGGCCGTGAGCCCCCAGGAAGAGGAAGAGTTTAGACGGCTGGCCGCCCTGCCTAACGTGTACGAAGTGATCAGCAAGTCTATCGCCCCCAGCATCTTCGGCGGCACCGATATGAAGAAGGCCATTGCCTGCCTGCTGTTCGGCGGCTCCAGAAAGAGACTGCCTGACGGCCTGACAAGACGGGGCGACATCAACCTGCTGATGCTGGGCGATCCTGGCACCGCCAAATCACAGCTGCTGAAGTTCGTGGAAAAGTGCAGCCCCATCGGCGTGTACACCAGCGGCAAGGGATCTTCTGCCGCCGGACTGACAGCCAGCGTGATGAGAGATCCCAGCAGCCGGAATTTCATCATGGAAGGCGGCGCTATGGTGCTGGCCGATGGCGGAGTCGTGTGCATCGACGAGTTCGACAAGATGCGCGAGGACGACCGGGTGGCCATTCACGAGGCTATGGAACAGCAGACCATCTCTATCGCCAAGGCCGGAATCACCACCACCCTGAACAGCCGGTGTAGCGTGCTGGCTGCCGCCAATAGCGTGTTCGGCAGATGGGACGAGACAAAGGGCGAGGACAACATCGACTTCATGCCCACCATCCTGAGCCGCTTCGACATGATCTTCATCGTGAAGGACGAGCACAACGAGGAACGGGACGTGATGCTGGCCAAGCACGTGATCACCCTGCACGTGTCCGCCCTGACACAGACACAGGCCGTGGAAGGCGAGATCGACCTGGCCAAACTGAAGAAGTTTATCGCCTACTGTCGCGTGAAGTGCGGCCCCAGACTGTCTGCCGAGGCCGCCGAGAAGCTGAAGAACCGGTACATCATCATGCGGAGCGGAGCCCGGCAGCACGAGAGAGACAGCGATCGGAGAAGCAGCATCCCCATCACCGTGCGGCAGCTGGAAGCCATCGTGCGGATTGCCGAAGCCCTGAGCAAGATGAAGCTCCAGCCCTTCGCCACCGAGGCCGACGTGGAAGAAGCTCTGAGACTGTTCCAGGTGTCAACCCTGGACGCCGCCCTGAGCGGAACACTGTCTGGCGTGGAAGGATTCACCAGCCAGGAAGATCAGGAAATGCTGAGCAGAATCGAGAAACAGCTGAAGAGAAGATTCGCCATCGGAAGCCAGGTGTCCGAGCACAGCATCATCAAGGACTTCACCAAGCAGAAGTACCCCGAGCACGCCATCCACAAGGTGCTCCAGCTGATGCTGAGAAGAGGCGAGATCCAGCACCGGATGCAGCGGAAGGTGCTGTACCGGCTGAAATGAGGCGCCGGAATCTTTGCAGGTGCTTACGCTTACTACCTAAACTACAATGGTGTTGTCGCTACTAGTGCCGCTTCTTCATCCACTGCATCTGGTGCTTCCGCTTCCGTTACCGGTTCTAAGAAACTTCAGAAACTTCAGAAACCAGTGCTGCCCCTAAGACATACACTACTGCCACTGTTACTCAATGTGATGACAATGGTTGTAACGTCAAGATAATCACCTCTCAAATACCTGAAGCTACTTCAACCGTCACCGCAACTAGTGCTTCTCCAAAGTCATACACTACTGTCACTTCTGAGGGTTCTAAAGCAACCTCATTGAGTTCTCACTGCCCGCTTTCCAGTCGGGAAACCTGTCGTGCCAGCTGCATTAATGAATCGGCCAACGCGCGGGGAGAGGCGGTTTGCGTATTGGGCGCTCTTCCGCTTCCTCGCTCACTGACTCGCTGCGCTCGGTCGTTCGGCTGCGGCGAGCGGTATCAGCTCACTCAAAGGCGGTAATACGGTTATCCACAGAATCAGGGGATAACGCAGGACAGAACATGTGAGCAGAAGGCCAGCACGAGGCCAGGAGCCGTAAGAAGGCCGCGTTGCTGGCGTTTTTCCATAGGCTCCGCCCCCTCGCGAGCTGAGGACTTAAGTGCTGAGGTCGCGAGCATTCAGAAGAGCCATTCAAGGGATCGATGTGGTCGCACGACTACGCATCCCTCTGGGCCTTACATAGCCGGATACAGTGACTTATCGATACGCGTGTTAACCTCGAGGCGGCCGCCATGGTGACAGGACTTGCATAGCTGCGTGCGGGGGAAGGAACTCTTGCGTGTGAGTATGTAGACCCCTGTACTACGGATGCGGGCAGAAGATGTGGGCAGAGACACCCGCGTCAAGTTCTCGACCTTCCCGTGGGAGGTGTTCCAGTCCGCCATACGACCATACCGTTCGGGCATGGCACTATGTACGCGATATCGCTAGCGTTTGCGGGGCACAGCAATATGCAGGCATGCAAGCTTGGCGTAATCATGGTCATAGCTGTTTCCTGTGTGAAATTGTTATCCGCTCACAATTCCACACAACATACGAGCCGGAAGCATAAAGTGTAAAGCCTGGGGTGCCTAATGAGTGAGCTAACTCACATTAATTGCGTTGCGCTCACTGCCCGCTTTCCAGTCGGGAAACCTGTCGTGCCAGCTGCATTAATGAATCGGCCAACGCGCGGGGAGAGGCGGTTTGCGTATTGGGCGCTCTTCCGCTTCCTCGCTCACTGACTCGCTGCGCTCGGTCGTTCGGCTGCGGCGAGCGGTATCAGCTCACTCAAAGGCGGTAATACGGTTATCCACAGAATCAGGGGATAACGCAGGAAAGAACATGTGAGCAAAAGGCCAGCAAAAGGCCAGGAACCGTAAAAAGGCCGCGTTGCTGGCGTTTTTCCATAGGCTCCGCCCCCCTGACGAGCATCACAAAAATCGACGCTCAAGTCAGAGGTGGCGAAACCCGACAGGACTATAAAGATACCAGGCGTTTCCCCCTGGAAGCTCCCTCGTGCGCTCTCCTGTTCCGACCCTGCCGCTTACCGGATACCTGTCCGCCTTTCTCCCTTCGGGAAGCGTGGCGCTTTCTCATAGCTCACGCTGTAGGTATCTCAGTTCGGTGTAGGTCGTTCGCTCCAAGCTGGGCTGTGTGCACGAACCCCCCGTTCAGCCCGACCGCTGCGCCTTATCCGGTAACTATCGTCTTGAGTCCAACCCGGTAAGACACGACTTATCGCCACTGGCAGCAGCCACTGGTAACAGGATTAGCAGAGCGAGGTATGTAGGCGGTGCTACAGAGTTCTTGAAGTGGTGGCCTAACTACGGCTACACTAGAAGAACAGTATTTGGTATCTGCGCTCTGCTGAAGCCAGTTACCTTCGGAAAAAGAGTTGGTAGCTCTTGATCCGGCAAACAAACCACCGCTGGTAGCGGTGGTTTTTTTGTTTGCAAGCAGCAGATTACGCGCAGAAAAAAAGGATCTCAAGAAGATCCTTTGATCTTTTCTACGGGGTCTGACGCTCAGTGGAACGAAAACTCACGTTAAGGGATTTTGGTCATGAGATTATCAAAAAGGATCTTCACCTAGATCCTTTTAAATTAAAAATGAAGTTTTAAATCAATCTAAAGTATATATGAGTAAACTTGGTCTGACAGTTACCAATGCTTAATCAGTGAGGCACCTATCTCAGCGATCTGTCTATTTCGTTCATCCATAGTTGCCTGACTCCCCGTCGTGTAGATAACTACGATACGGGAGGGCTTACCATCTGGCCCCAGTGCTGCAATGATACCGCGAGACCCACGCTCACCGGCTCCAGATTTATCAGCAATAAACCAGCCAGCCGGAAGGGCCGAGCGCAGAAGTGGTCCTGCAACTTTATCCGCCTCCATCCAGTCTATTAATTGTTGCCGGGAAGCTAGAGTAAGTAGTTCGCCAGTTAATAGTTTGCGCAACGTTGTTGCCATTGCTACAGGCATCGTGGTGTCACGCTCGTCGTTTGGTATGGCTTCATTCAGCTCCGGTTCCCAACGATCAAGGCGAGTTACATGATCCCCCATGTTGTGCAAAAAAGCGGTTAGCTCCTTCGGTCCTCCGATCGTTGTCAGAAGTAAGTTGGCCGCAGTGTTATCACTCATGGTTATGGCAGCACTGCATAATTCTCTTACTGTCATGCCATCCGTAAGATGCTTTTCTGTGACTGGTGAGTACTCAACCAAGTCATTCTGAGAATAGTGTATGCGGCGACCGAGTTGCTCTTGCCCGGCGTCAATACGGGATAATACCGCGCCACATAGCAGAACTTTAAAAGTGCTCATCATTGGAAAACGTTCTTCGGGGCGAAAACTCTCAAGGATCTTACCGCTGTTGAGATCCAGTTCGATGTAACCCACTCGTGCACCCAACTGATCTTCAGCATCTTTTACTTTCACCAGCGTTTCTGGGTGAGCAAAAACAGGAAGGCAAAATGCCGCAAAAAAGGGAATAAGGGCGACACGGAAATGTTGAATACTCATACTCTTCCTTTTTCAATATTATTGAAGCATTTATCAGGGTTATTGTCTCATGAGCGGATACATATTTGAATGTATTTAGAAAAATAAACAAATAGGGGTTCCGCGCACATTTCCCCGAAAAGTGCCACCTGACGTCTAAGAAACCATTATTATCATGACATTAACCTATAAAAATAGGCGTATCACGAGGCCCTTTCG

**Table S1 Expression constructs used in this study**

| **Insert** | **Vector backbone** | **Uniprot ID** | **Tag** | **Source** |
| --- | --- | --- | --- | --- |
| MCM5-6-7-Psf2-Cdc45 | pBig1A | P33992, MCM5_HUMAN  Q14566, MCM6_HUMAN  P33993, MCM7_HUMAN  Q9Y248, PSF2_HUMAN  O75419, CDC45_HUMAN | Psf2: Dual Strep, C terminal | This study |
| MCM2-4-Sld5-Psf1-3 | pBig1B | P49736, MCM2_HUMAN  P33991, MCM4_HUMAN  Q9BRT9, SLD5_HUMAN  Q14691, PSF1_HUMAN  Q9BRX5, PSF3_HUMAN | No tag | This study |
| MCM3 | pBig1C | P25205, MCM3_HUMAN | MCM3: FLAG, N terminal | This study |
| Polymerase ε | pACEBac1 | Q07864, POLE1_HUMAN  P56282, POLE2_HUMAN  Q9NRF9, POLE3_HUMAN  Q9NR33, POLE4_HUMAN | POLE2: FLAG, N terminal | This study |
| Tipin-Timeless | pACEBac1 | Q9UNS1, TIM_HUMAN,  Q9BVW5, TIPIN_HUMAN | Timeless: FLAG, N terminal | This study |
| Claspin | pBig1A | Q9HAW4, CLSPN_HUMAN | Dual Strep, C terminal | This study |
| AND-1 | pACEBac1 | O75717, WDHD1_HUMAN | Dual Strep, N terminal | This study |
| RFC1 | pACEBac1 | P35251, RFC1_HUMAN | Dual Strep, N terminal | This study |
| RFC2-3 | pACEBac1 | P35250, RFC2_HUMAN  P40938, RFC3_HUMAN | No tag | This study |
| RFC4-5 | pACEBac1 | P35249, RFC4_HUMAN  P40937, RFC5_HUMAN | No tag | This study |
| CTF18 | pACEBac1 | Q8WVB6, CTF18_HUMAN | Dual strep, N terminal | This study |
| CTF8/DSCC1 | pACEBac1 | P0CG13, CTF8_HUMAN  Q9BVC3, DCC1_HUMAN | No tag | This study |
| POLA2 | pACEBac1 | Q14181, DPOA2_HUMAN | No tag | This study |
| PRIM1 | pACEBac1 | P49642, PRI1_HUMAN | No tag | This study |
| CST | pACEBac1 | Q2NKJ3, CTC1_HUMAN  Q9H668, STN1_HUMAN  Q86WV5, TEN1L_HUMAN | CTC1: FLAG, N terminal  TEN1: His_6_, N terminal | This study |
| FEN1 |  |  | FLAG, C terminal | This study |
| Ligase I |  |  | FLAG, C terminal | This study |
| Polymerase δ | MultiBac |  | POLD4: His_6_, N terminal | Gift from Dr Samir Hamdan |
| POLA1 | pACEBac1 |  | Dual Strep, N terminal | Gift from Dr Joseph Yeeles |
| PRIM2 | pACEBac1 |  | No tag | Gift from Dr Joseph Yeeles |
| RPA | pET11d |  | No tag | Gift from Dr Marc Wold (Addgene, 102613) |
| PCNA | pET16b |  | No tag | Gift from Dr Andrew Deans  (Addgene, 134898) |
| Biotinylated PCNA | pET28a |  | Avitag, Dual His_6_, N terminal | This study |
| BLM WT and helicase dead | pFastBac1 |  | MBP, N terminal  His_10_, C terminal | Gift from Dr Petr Cejka |
| LacI | pD861 |  | His_6_, C terminal | Gift from Dr John Diffley |
| Shelterin WT | pACEBac1 | P54274, TERF1_HUMAN Q15554, TERF2_HUMAN Q9NYB0, TE2IP_HUMAN Q96AP0, ACD_HUMAN Q9NUX5, POTE1_HUMAN Q9BSI4, TINF2_HUMAN | TINF2: Strep N terminal | Gift from Prof Sebastian Guettler |
| Shelterin -TRF1 | pACEBac1 | Q15554, TERF2_HUMAN Q9NYB0, TE2IP_HUMAN Q96AP0, ACD_HUMAN Q9NUX5, POTE1_HUMAN Q9BSI4, TINF2_HUMAN | TINF2: Strep N terminal | Gift from Prof Sebastian Guettler |
| Shelterin -TRF2/RAP1 | pACEBac1 | P54274, TERF1_HUMAN Q9NYB0, TE2IP_HUMAN Q96AP0, ACD_HUMAN Q9NUX5, POTE1_HUMAN Q9BSI4, TINF2_HUMAN | TINF2: Strep N terminal | Gift from Prof Sebastian Guettler |
| Shelterin -TPP1/POT1 | pACEBac1 | P54274, TERF1_HUMAN Q15554, TERF2_HUMAN Q9NYB0, TE2IP_HUMAN Q9BSI4, TINF2_HUMAN | TINF2: Strep N terminal | Gift from Prof Sebastian Guettler |
| Shelterin TRF1 ΔMyb and TRF2 ΔMyb  TRF1 (amino acids 375-431 deletion)  TRF2 (amino acids 484-534 deletion) | pACEBac1 | P54274, TERF1_HUMAN Q15554, TERF2_HUMAN Q9NYB0, TE2IP_HUMAN Q96AP0, ACD_HUMAN Q9NUX5, POTE1_HUMAN Q9BSI4, TINF2_HUMAN | TINF2: Strep N terminal | PCR mutagenesis of shelterin |
| POT1 | pACEBac1 | Q9NUX5, POTE1_HUMAN | No tag | Gift from Prof Sebastian Guettler |
| POT1 F62A | pACEBac1 | Q9NUX5, POTE1_HUMAN | No tag | PCR mutagenesis of POT1 |
| POT1 ΔOB1  (amino acids 1-126 deletion) | pACEBac1 | Q9NUX5, POTE1_HUMAN | No tag | PCR mutagenesis of POT1 |
| TPP1 | pACEBac1 | Q96AP0, ACD_HUMAN | Dual Strep, N terminal | Gift from Prof Sebastian Guettler |

**Table S2 Oligonucleotides used for replication fork construction and initiation**

| **Oligo** | **Use** | **Modifications** | **Sequence (5’-3’)** |
| --- | --- | --- | --- |
| TBL237 | Annealed fork, leading strand oligo, PstI site | 5’ Phospho and 3’ phosphothiroate bonds *  PAGE purified | Phos-GATGTGGTAGGAAGTGAGAATTGGAGAGTGTGTTTTTTTTTTTTTTTTTTTTTTTTTTTTTTTTTTTTTTTTGAGGAAAGAATGTTGGTGAGGGTTGGGAAGTGGAAGGATGGGCTCGAGAGGTTTTTTTTTTTTTTTTTTTTTTTTTTTTT*T*T*T*T*T |
| TBL238 | Annealed fork, lagging strand oligo, PstI site | PAGE purified | TTTTTTTTTTTTTTTTTTTTTTTTTTTTTTTTTTTTTTTTTTTTTTTTTTTTTTTTTTTTCACACTCTCCAATTCTCACTTCCTACCACATCTGCA |
| TBL253 | Replication assay initiating oligo | 5’ Biotin | Bio-CCTCTCGAGCCCATCCTTCCACTTCCCAACCCTCACC |
| TBL272 | Annealed fork, leading strand oligo, HindIII site | 5’ Phospho and 3’ phosphothiroate bonds *  PAGE purified | Phos-AGCTTATGTGGTAGGAAGTGAGAATTGGAGAGTGTGTTTTTTTTTTTTTTTTTTTTTTTTTTTTTTTTTTTTTTTTGAGGAAAGAATGTTGGTGAGGGTTGGGAAGTGGAAGGATGGGCTCGAGAGGTTTTTTTTTTTTTTTTTTTTTTTTTTTTT*T*T*T*T*T |
| TBL273 | Annealed fork, lagging strand oligo, HindIII site | PAGE purified | TTTTTTTTTTTTTTTTTTTTTTTTTTTTTTTTTTTTTTTTTTTTTTTTTTTTTTTTTTTTCACACTCTCCAATTCTCACTTCCTACCACATA |
| TBL296 | Shelterin EMSA | 5’ Biotin  PAGE purified | CGTCTATATTCTATTGTCTCTTAGGGTTAGGGTTAGGGTTAGGGTTAGGGTTAGGGTTAGGGTTAGGGTTAACATCAGTCTCACATAGATTAGCTCACGC |
| TBL297 | Shelterin EMSA | PAGE purified | GCGTGAGCTAATCTATGTGAGACTGATGTTAACCCTAACCCTAACCCTAACCCTAACCCTAACCCTAACCCTAACCCTAAGAGACAATAGAATATAGACG |
| TBL363 | Annealed fork, leading strand oligo, NotI site | 5’ Phospho and 3’ phosphothiroate bonds *  PAGE purified | 5’PhosGGCCGCATGTGGTAGGAAGTGAGAATTGGAGAGTGTGTTTTTTTTTTTTTTTTTTTTTTTTTTTTTTTTTTTTTTTTGAGGAAAGAATGTTGGTGAGGGTTGGGAAGTGGAAGGATGGGCTCGAGAGGTTTTTTTTTTTTTTTTTTTTTTTTTTTTT*T*T*T*T*T |
| TBL364 | Annealed fork, lagging strand oligo, NotI site | PAGE purified | TTTTTTTTTTTTTTTTTTTTTTTTTTTTTTTTTTTTTTTTTTTTTTTTTTTTTTTTTTTTCACACTCTCCAATTCTCACTTCCTACCACATGC |
| CLB353 | BLM helicase assay top strand |  | TTTTTTTTTTTTTTTTTTTTTTTTTTTTTTTTTTTTTTTTCGACCGTGCCAGCCTAAACCAGACTGCTACACACAGGATCGTTCGGTCTC |
| CLB354 | BLM helicase assay bottom strand |  | GAGACCGAACGATCCTGTGTGTAGCAGTCTGGTTTAGGCTGGCACGGTCGTTTTTTTTTTTTTTTTTTTTTTTTTTTTTTTTTTTTTTTT |
